# Effects of Drought on Inflorescence Yield, and Secondary Metabolites in *Cannabis sativa* L.

**DOI:** 10.1101/2025.02.16.638548

**Authors:** Itamar Shenhar, Idan Ifrach, Ohad Barkan, Ohad Guberman, Zohar Kerem, Dvir Taler, David Meiri, Menachem Moshelion

## Abstract

- The use of medical products derived from *Cannabis sativa* L. has increased significantly in recent years. While drought is known to negatively affect the yields of many crops, growers often recommend controlled periods of drought for cannabis cultivation to increase concentrations of secondary metabolites. This is especially pertinent when considering the relationship between medicinal effects and the secondary-metabolite profile.
- We examined the effects of tightly controlled drought treatments on biochemical (111 phytocannabinoids and 132 terpenoids), physiological, and anatomical responses of three Type-I chemotype cultivars, specifically the THCA-dominant cultivars ‘Odem’, ‘MVA’, and ‘187’.
- Our results revealed strong correlations between inflorescence and phytocannabinoid yields, on the one hand, and cumulative transpiration on the other (0.96 < *r*^2^ < 0.99). Drought treatment reduced canopy conductance, with inflorescence weight decreasing by 40% and total fresh weight decreasing by 48%. The concentrations of the major phytocannabinoids, THCA and CBGA, decreased over increasing levels of drought stress (by 26% and 61%, respectively). Interestingly, terpene concentrations showed greater stress-induced variation, and that variation was genotype-dependent.
- Our findings suggest that the decreases in inflorescence weight and concentrations of major phytocannabinoids under drought conditions are mainly due to a lack of biochemical-production processes, as opposed to metabolic degradation.

## Introduction

In recent decades, cannabis (*Cannabis sativa* L.) has been increasingly utilized for both recreational and medicinal purposes, becoming the third most commonly used controlled substance globally (Connor et al., 2021) and used medicinally in both raw flower and extract forms (Kalant, 2001; Ware et al., 2005). The medical use of cannabis products has risen notably due to evidence supporting its benefits for various conditions and its acceptable risk profile (JE et al., 1999; Walsh et al., 2013). The primary compounds of interest in cannabis are phytocannabinoids and terpenoids (Gülck and Møller, 2020), which contribute to its medicinal properties. These compounds are known to interact synergistically, producing what has been termed the **“**entourage effect**”** (Russo, 2011; Koltai and Namdar, 2020; Sommano et al., 2020).

Phytocannabinoids are found in various plants and fungi, with cannabis containing the largest variety of phytocannabinoid compounds (Gülck and Møller, 2020). These compounds are mainly produced in glandular trichomes, with high concentrations in the inflorescence (Livingston et al., 2020), and interact with the endocannabinoid system in mammals to regulate a range of physiological and pathological functions (Zou and Kumar, 2018; Chanda et al., 2019). Cannabis plants are classified into three main chemotypes based on the relative concentrations of the major cannabinoids tetrahydrocannabinolic acid (THCA) and cannabidiolic acid (CBDA). Chemotype Ⅰ is defined by a high THCA:CBDA ratio (>1.0), Chemotype Ⅱ has an intermediate ratio (0.5–2.0) with fairly equivalent levels of THCA and CBDA, and Chemotype Ⅲ is defined by a low ratio (<0.5), with high levels of CBDA and low levels of THCA (Small and Beckstead, 1973).

The biosynthesis of phytocannabinoids begins with cannabigerolic acid (CBGA), which is synthesized from olivetolic acid and geranyl diphosphate (GPP) in the presence of cannabigerolic acid synthase. THCA and CBDA are subsequently produced from CBGA via specific synthases. In addition, under stress conditions, including oxidative degradation, CBGA is decarboxylated to yield tetrahydrocannabinol (THC), cannabidiol (CBD), cannabigerol (CBG), and cannabinol (CBN) (Thomas and ElSohly, 2016).

Terpenoids, the primary constituents of essential oils, are produced by various plants and animals and have health benefits for humans. Terpenoids contribute to the aroma of cannabis and are considered to enhance the activity of its phytocannabinoids (Ben-Shabat et al., 1998; Paduch et al., 2007; Russo, 2011; Koltai and Namdar, 2020). The biosynthesis of terpenoids involves isoprenoid diphosphate precursors and follows two pathways: the plastidial methylerythritol phosphate (MEP) pathway for the production of monoterpenoids and the cytosolic mevalonate (MEV) pathway for the production of sesquiterpenoids (Sommano et al., 2020).

Breeding cannabis to produce inflorescences with high levels of secondary metabolites poses an agronomic challenge. Selective breeding has been used in efforts to increase levels of phytocannabinoids and terpenoids, although traditional methods lack the precision of techniques based on genetic engineering (Parsons et al., 2019).

The growth rate, yield, and secondary-metabolite production of cannabis are all significantly influenced by atmospheric conditions, the growing medium, its mineral composition, and irrigation protocols (Chandra et al., 2008; Burgel et al., 2020; Danziger and Bernstein, 2021a; Danziger and Bernstein, 2021b). Water availability is a critical factor, with water deficit causing stomatal closure, with subsequently reduced CO_2_ assimilation and decreased transpiration. Drought stress typically results in stomatal closure, reducing transpiration and photosynthesis, thereby decreasing the yields of many crops (Sadras and Calviño, 2001; Cetin and Bilgel, 2002; Garan et al., 2016). Temperature can directly affect biochemical activities and also indirectly influences atmospheric vapor pressure deficit (VPD), which controls stomatal aperture and photosynthesis (Yaaran et al., 2019), thereby limiting photosynthesis in response to reduced CO_2_ assimilation and electron-transport-chain activity (Taiz et al., 2015). The resulting decline in energy production can significantly limit the biosynthesis of secondary metabolites (Thoma et al., 2020). Transpiration drives the uptake and transport of mobile minerals and is essential for plant metabolism. Standardizing plant growth and production is challenging due to the close relationship between productivity and environmental conditions, which necessitates an understanding of genotype– environment interactions (G × E), particularly with regard to drought stress (Moshelion, 2020).

Crop plants exhibit various anatomical, physiological, morphological, and biochemical responses to drought stress, including modification of their secondary metabolic profiles to promote the synthesis of terpenoids, phenolic compounds, and nitrogen compounds (Borges et al., 2017). In *Vitis vinifera*, for instance, flavonoid, anthocyanin, and catechin concentrations increase when irrigation is limited (Kennedy et al., 2002). A study by Caplan et al. (2019), found that Chemotype-II cannabis plants subjected to drought had higher concentrations of major phytocannabinoids without any reduction in yield, suggesting a unique mechanism of drought-stress resistance in this chemotype. Additionally, the application of the drought stress-related phytohormone ABA was also reported to increase THC concentrations (Mansouri et al., 2009). However, Morgan et al. (2024) revealed that severe drought significantly reduces the phytocannabinoid and floral yields of Chemotype-Ⅲ cannabis plants. Despite these studies it remains unclear how different levels of drought affect the most commonly used chemotype, Chemotype I.

Our study aimed to investigate the impact of standardized, controlled drought treatments on the inflorescence yield and concentrations of major secondary metabolites under both controlled and non-controlled atmospheric conditions. We utilized three Chemotype-Ⅰ cultivars, grown under three different levels of drought in a highly controlled environment and in a commercial greenhouse with non-controlled conditions. We measure yield in terms of inflorescence dry weight and concentrations of phytocannabinoids and terpenoids.

Our hypotheses were as follows:

1. The inflorescence yield of drought-exposed plants will decrease in correlation with the level of drought stress.
2. The concentrations of major secondary metabolites will increase in correlation with the level of drought level.
3. The trends observed under non-controlled conditions will be similar to those observed under controlled conditions, but with greater variation.

This research is pioneering in its examination of the response of Chemotype-I cannabis plants to controlled levels of drought, its integration of controlled atmospheric conditions with commercial greenhouse conditions, and its potential for providing insights for commercial growers regarding the use of controlled drought under various atmospheric conditions.

## Materials and Methods

### Study locations, plant material, and growing conditions

The research was conducted from September 2020 to June 2022 at two sites: a controlled environment at the Robert H. Smith Faculty of Agriculture, Food and Environment at the Hebrew University of Jerusalem, Rehovot, Israel, and a non-controlled environment at the BOL Pharma Ltd. Facility, Revadim, Israel. Three commercial Chemotype-I cultivars were used in this study. We primarily focused on the ‘Odem’ cultivar, with one repetition in both controlled and non-controlled environments conducted on the ‘MVA’ cultivar and ‘187’ cultivars, respectively (BOL Pharma). The study consisted of three experiments conducted in a controlled environment (growth chamber) and two conducted in a non-controlled environment (greenhouse; see Table 1 and Fig. S1a, b). ‘Odem’ and ‘187’ both exhibited a sativa-like elongated plant structure; whereas ‘MVA’ exhibited indica-like compact growth (Bachir et al., 2022; Fig. 1a).

**Fig. 1.**
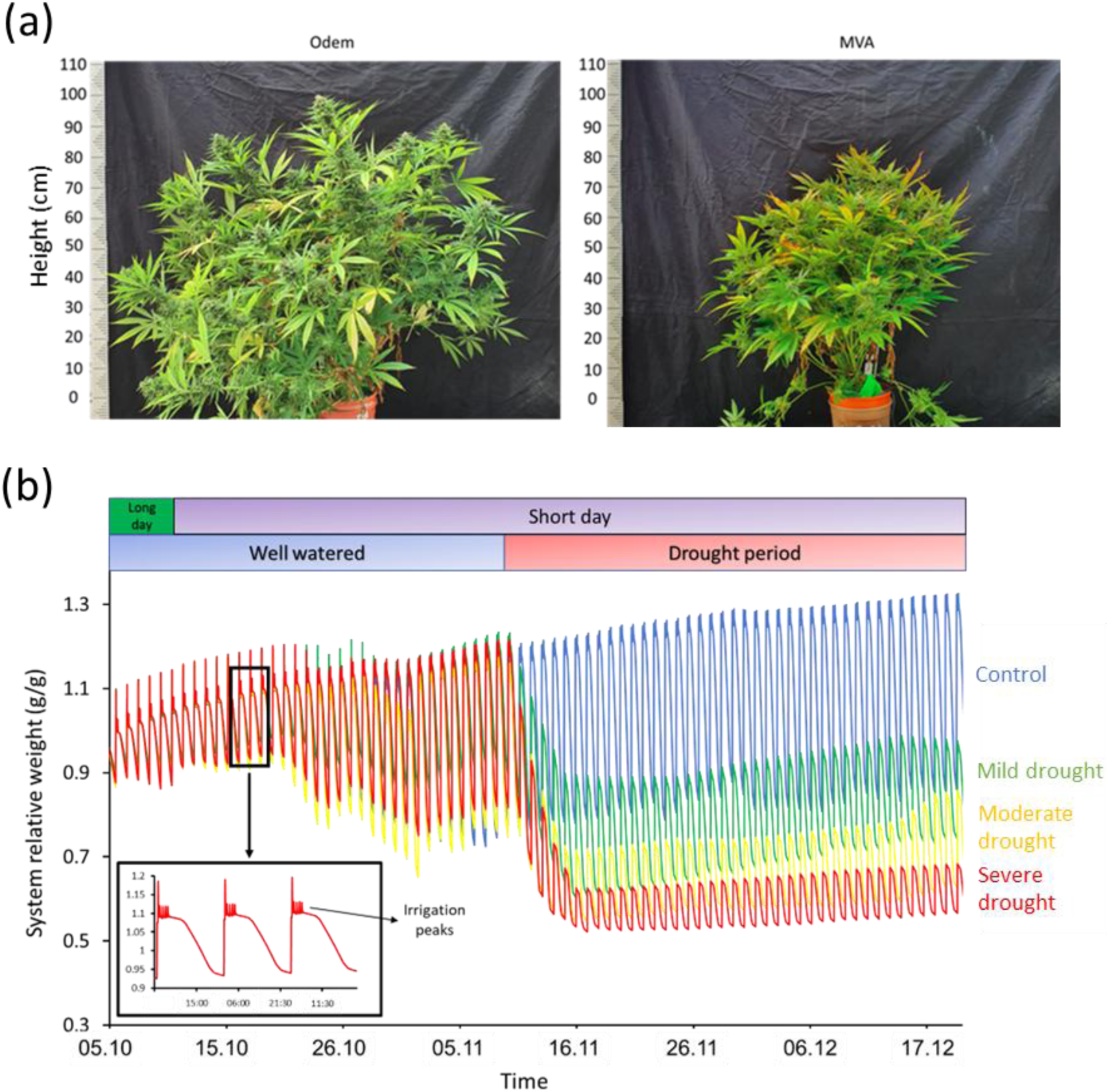
Representative images and experimental scenario. (a) Images of ‘Odem’ and ‘MVA’ plants grown under non-controlled atmospheric conditions. (b) Relative weights of four representative plants under controlled atmospheric conditions: well-watered (blue), mild drought (green, 75% transpiration), moderate drought (yellow, 50% transpiration), and severe drought (red, 25% transpiration). The upper bar indicates the photoperiod (green for vegetative growth, purple for reproductive growth) and the lower bar indicates the drought period (blue for well-watered, red for drought). The inset shows a 3-day weight profile of a well-watered plant, highlighting nightly irrigation to field capacity and daytime weight loss due to transpiration. While all control plants were subjected to this regime, drought-stressed plants received a single irrigation peak tailored to their particular treatment.

**Table 1:** Overview of experiments, figures, and data tables. This table summarizes experiments on different Cannabis cultivars (Odem, MVA, and 187) under controlled and non-controlled atmospheric conditions. Drought levels are categorized as control, low drought (75% transpiration), medium drought (50% transpiration), and severe drought (25% transpiration). Key figures illustrate physiological, biochemical, and yield responses, while supplementary materials provide additional details on experimental conditions and results.

| Experiment | Location | Cultivar | Experiment date | Atmospheric conditions | Related figures | Related tables |
| --- | --- | --- | --- | --- | --- | --- |
| 1 | Growth room | Odem | 05.10.20–<br>20.12.20 | Controlled | 1, 2, 3, 4, 5, 6,<br>8, S1, S2 | S1, S3, S4, S5,<br>S15, S16 |
| 2 | Growth room | Odem | 08.01.21–<br>20.03.21 | Controlled | 6, 8 | S1, S3, S6, S7,<br>S8, S17, S18,<br>S19 |
| 3 | Growth room | MVA | 05.12.21–<br>16.02.22 | Controlled | 7, 9 | S1, S3, S9, S10,<br>S20, S21 |
| 4 | Commercial<br>greenhouse | Odem | 20.04.21–<br>01.07.21 | Non-controlled | 1, 3, S1, S3 | S1, S2, S3, S11,<br>S12 |
| 5 | Commercial<br>greenhouse | 187 | 13.10.21–<br>26.12.21 | Non-controlled | 1, S3 | S1, S2, S3, S13,<br>S14 |
| 6 | Growth room | Odem | 29.03.22–<br>14.06.22 | Controlled | 2 | S1, |

**Table 1: Overview of experiments, figures, and data tables.**
| Experiment | Location | Cultivar | Experiment date | Atmospheric conditions | Related figures | Related tables |
| --- | --- | --- | --- | --- | --- | --- |
| 1 | Growth room | Odem | 05.10.20–<br>20.12.20 | Controlled | 1, 2, 3, 4, 5, 6,<br>8, S1, S2 | S1, S3, S4, S5,<br>S15, S16 |
| 2 | Growth room | Odem | 08.01.21–<br>20.03.21 | Controlled | 6, 8 | S1, S3, S6, S7,<br>S8, S17, S18, S19 |
| 3 | Growth room | MVA | 05.12.21–<br>16.02.22 | Controlled | 7, 9 | S1, S3, S9, S10,<br>S20, S21 |
| 4 | Commercial<br>greenhouse | Odem | 20.04.21–<br>01.07.21 | Non-controlled | 1, 3, S1, S3 | S1, S2, S3, S11,<br>S12 |
| 5 | Commercial<br>greenhouse | 187 | 13.10.21–<br>26.12.21 | Non-controlled | 1, S3 | S1, S2, S3, S13,<br>S14 |
| 6 | Growth room | Odem | 29.03.22–<br>14.06.22 | Controlled | 2 | S1, |

Cuttings from the same mother plant were grown for 21 days before being transplanted into 7000-ml pots filled with a peat-based growth medium (540W, Kekkila Ltd., Finland) and fertilized with 3,3,9+6 solution during the vegetative growth period and 1.3,6,8+6 solution during the short-day flowering growth period (Super Shefer, ICL Ltd., Israel), in addition to a calcium and magnesium solution (5.8,0,0+3+3; CalMag, ICL Ltd., Israel). Fertilizer was applied through two fixed-ratio (1:500) fertigation-dosing pump (D25-F-02, Dosatron Ltd., France) systems with a custom nutrient composition specifically tailored for cannabis cultivation (Table S1).

### Controlled and non-controlled ambient conditions

Controlled atmospheric conditions were achieved using custom-made LED lights (CXB3590, Cree Lighting Ltd., WI, USA) with two spectra: 3000K and 4000K per lamp. Each day started with 2 h of 4000K light, combined 3000K and 4000K light during the main photoperiod, and 2 h of 3000K light at the end of the day. To ensure consistent light intensity at the level of the plant canopy, we manually adjusted the lamps every few days to maintain a target of 1200 µmol m⁻^2^ s⁻¹ photosynthetic photon flux density at the top of the canopy at noon, measured using a LI-COR LI-250A light meter. Artificial lighting was not used in the greenhouse with non-controlled conditions, except to modify the photoperiod for morphological development. Air circulation in the controlled chambers was maintained using fans and atmospheric conditions were continuously monitored. The long-day photoperiod for the vegetative growth stage consisted of 16 h light / 8 h dark; whereas the short-day photoperiod for flowering consisted of 12 h light / 12 h dark.

The controlled drought experiment employed a multilevel multifactorial design, which required continuous measurements to ensure precise replication (Moshelion et al., 2024). Atmospheric conditions were continuously monitored using a meteorological monitoring station (WatchDog 2745, Spectrum Technologies Ltd., Wales; Fig. S1) through the PlantArray platform (Plant-DiTech Ltd., Israel). Drought experiments in pots are inherently challenging due to the pot effect, especially when growing large plants (Gosa et al., 2019a). To address this challenge, automatic feedback irrigation via the PlantArray platform was used to impose precise and standardized drought treatments tailored to each plant’s physiological response. The drought treatments were as follows: mild drought = 75% of the average transpiration of the last 3 days of the well irrigated period; moderate drought = 50% of the average transpiration of the last 3 days of the well irrigated period; severe drought = 25% of the average transpiration of the last 3 days of the well irrigated period; and control = 120% of the previous day’s transpiration, followed by four minimal irrigation pulses to ensure the substrate reached field capacity (Fig. 1b).

### Physiological measurement using the FFP system

The Functional Phenotyping Platform (FFP) system, PlantArray (Plant-DiTech Ltd., Yavne, Israel), incorporates highly sensitive load-cell lysimeters for real-time physiological measurements. The system enabled continuous monitoring of environmental conditions, including photosynthetic photon flux density, temperature, relative humidity, and vapor pressure deficit (VPD). The following whole-plant physiological traits were recorded: daily transpiration (g/day), transpiration rate (g/min), plant net weight (g), water-use efficiency (WUE, g biomass / g water), calculated plant weight (CPW), transpiration normalized to plant weight (E, TR / CPW), canopy stomatal conductance (Gsc, E / VPD), and cumulative transpiration (sum of daily transpiration). The daily light integral (DLI), representing the total number of photons reaching 1 m^2^ per day, was calculated as:

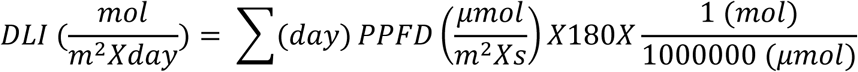

Physiological and environmental parameters were continuously monitored. Data processing and parameter calculations followed the methodologies described by Halperin et al. (2017) and Dalal et al. (2020). Morphological traits, such as new leaf emergence and plant height, were monitored by labeling and numbering leaves and measuring plant height at regular intervals.

### Harvest and biomass measurements

Harvested plants were separated into inflorescences, leaves, and stems. Samples were air-dried for 7 days and then oven-dried at 60°C. The dry weight of each organ was measured.

### Quantification of phytocannabinoids and terpenoids from plants grown under controlled conditions

Primary and secondary inflorescence samples were sampled, dried, and stored under controlled conditions, to preserve volatile terpene content and prevent fungal accumulation. Samples from plants grown under controlled conditions were crushed and digested. The levels of 111phytocannabinoids in those samples were then quantified using U-HPLC with an ultraviolet detector (UltiMate 3000, Thermo Fisher, Germany) and LC-MS with TM coupled with an Q Exactive focus hybrid quadrupole-Orbitrap MS (Q Exactive Plus, Thermo Fisher, Germany), as described by Milay et al. (2020) and Berman et al. (2018). A static headspace injection method followed by gas chromatography-mass spectrometry (Shapira et al., 2019) was used to quantify levels of 132 terpenoids in tissue samples from plants grown under controlled conditions We then used the collected data to analyze and plot the phytocannabinoids and terpenoids that were detected in all of the inflorescence samples.

Levels of major phytocannabinoids in the tissue samples from plants grown in the non-controlled environment were quantified using HPLC in a commercial laboratory that specializes in phytocannabinoid analysis (BOL Pharma Ltd., Revadim, Israel). A coffee grinder was used to grind dried inflorescence samples into a homogenous powder. Five hundred mg of powder were measured and inserted into 50-ml centrifuge tubes with 45 ml HPLC-grade ethanol (UN1170, Bio-Lab Ltd., Jerusalem, Israel). The 50-ml Falcon tubes were placed on a vibrating shaker (Vibramax 100, Heidolph Ltd., Germany) for 25 min and then transferred to an ultrasonic bath (DC-150H, MRC Ltd., England) for an additional 30 min. Later, the centrifuge tubes were centrifuged at 4000 rpm for 10 min (ROTOFIX 32A centrifuge, Hettich Ltd., Germany). One ml of the lowest density upper solution was taken and diluted with 19 ml HPLC-grade methanol (UN 1230, Bio-Lab Ltd., Israel) for 20 ml solution in total. A standard injector was used to filter the solutions through a 0.45-µm polytetrafluoroethylene filter into dedicated glass vials. The vials were inserted into an instrument for high-pressure liquid chromatography (e2695 XE, Waters Ltd., USA), to measure phytocannabinoid concentrations relative to standard solutions (Table S2).

To evaluate the moisture content of the dried samples, 500–600 mg of dried inflorescence powder were weighed and inserted into a moisture analyzer (HE53, Mettler-Toledo Ltd., Greifensee, Switzerland). The cannabinoid concentrations were calculated according to the moisture content of the sample, using the equation:

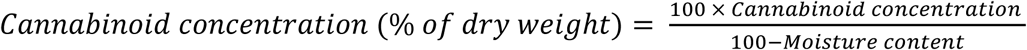

The total THC content was calculated as:

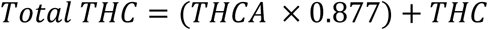

### Statistical analysis

Statistical analyses were performed using JMP, version 16.0 (SAS Institute Inc., NC, USA) and Microsoft Excel. Significance was determined using one-way ANOVA with Tukey’s Honest Significant Difference (HSD) test (*p* < 0.05) and *t*-tests (*p* < 0.05). Pearson correlation coefficients and regression fits were calculated to determine relationships among traits. Graphs, heatmaps, and Venn diagrams were generated using Excel.

## Results

### Physiological behavior of cannabis at different growth stages

We first characterized key physiological patterns of cannabis throughout its growth under well-watered conditions, focusing on the transformation from long-day to short-day photoperiods (Fig. 2). This characterization revealed three distinct physiological phases. Phase I is a vegetative phase that occurs when the plants were exposed to long-day conditions. Phase II is a transitional phase that occurs under short-day conditions and results in period of rapid vegetative growth (in terms of plant weight, leaf number, and height). In our study, Phase II lasted approximately 3 weeks and culminated in the onset of inflorescence development. Phase III is the inflorescence-development phase, which is characterized by a significant reduction in vegetative growth, stabilization of daily transpiration rates, and biomass accumulation that is primarily driven by inflorescence formation, resulting in reduced WUE.

**Fig. 2.**
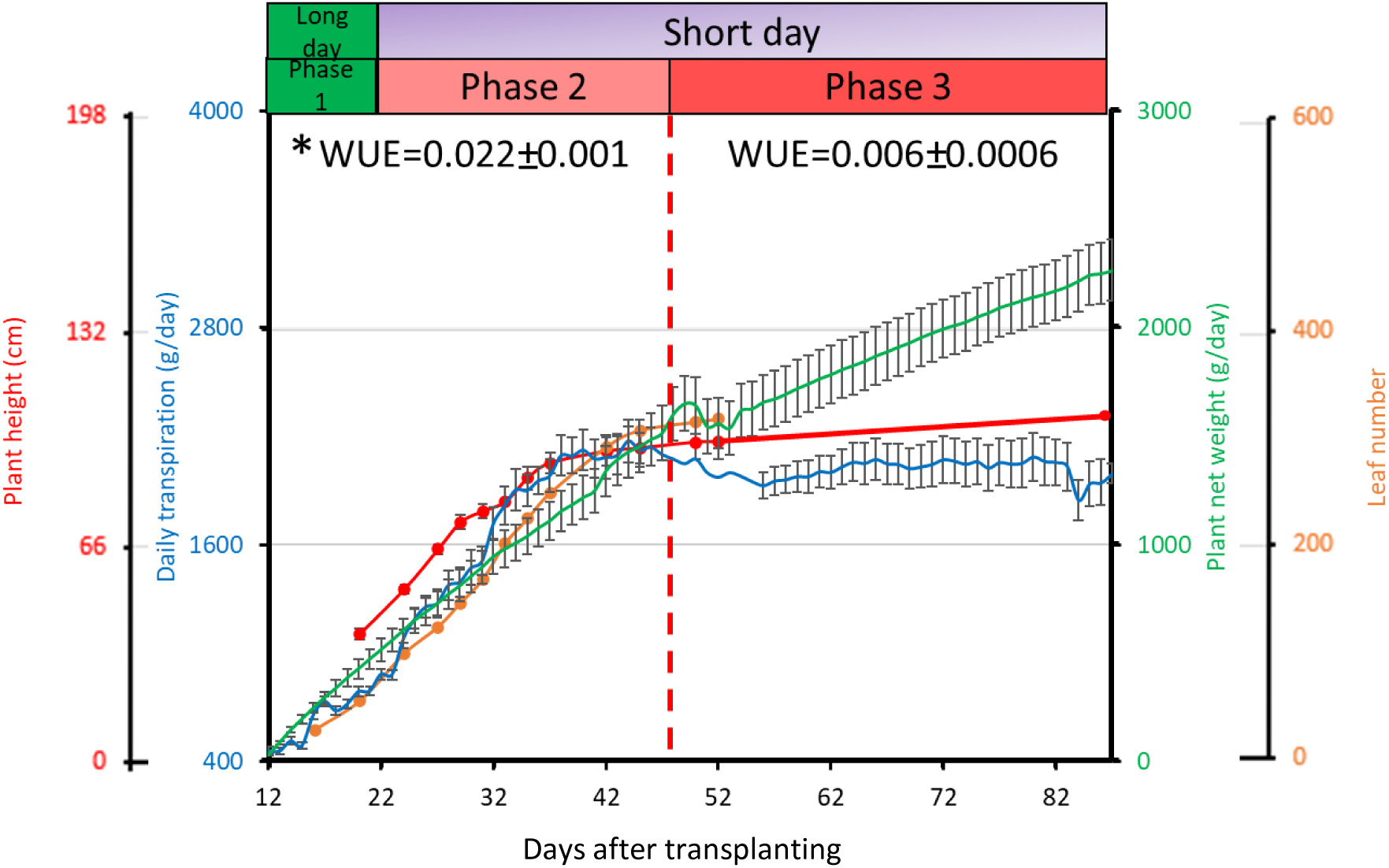
Physiological characteristics of the ‘Odem’ cultivar across growth stages under well-irrigated conditions. Mean ± SE values of daily transpiration (blue), biomass (green), plant height (red), and leaf number (orange) for a representative experiment conducted in the ‘Odem’ cultivar under controlled atmospheric conditions (Experiment 6, Table 1). Plants were grown under well-irrigated conditions with no drought stress. At the top of the figure, the upper bar indicates the duration of the photoperiod (green for long-day conditions, purple for short-day conditions), while the lower bar shows the physiological growth phases: Phase I (vegetative growth), Phase II (transition to flowering, indicated by a red vertical dashed line), and Phase III (inflorescence development). Water-use efficiency (WUE, g plant / g transpired water) was calculated for Phases II and III. The asterisk (*) indicates a significant difference (Student’s t-test; n = 22, p ≤ 0.005).

A close look at the shift to short-day lighting revealed a 28% decrease in the daily light integral (DLI) at the time of that change, which was accompanied by a 15% reduction in daily transpiration. However, this reduction in transpiration was followed by a recovery and a subsequent increase in transpiration over the following days. This recovery was likely due to continued plant growth and the development of new leaves (Fig. S2). During Phase II, we observed a notable surge in daily transpiration, which persisted for approximately 21 days. This increase coincided with rapid increases in fresh biomass, plant height, and the rate at which new leaves emerged. The WUE during this phase was calculated to be 0.022 ± 0.001 g plant / g water. At the conclusion of Phase II and throughout Phase III, daily transpiration stabilized at approximately ~2 L per day. This stabilization was accompanied by a marked decrease in the rates of plant growth and leaf emergence. However, fresh biomass continued to accumulate, driven by inflorescence development. Consequently, the WUE declined significantly to 0.006 ± 0.0006 g plant / g water (Fig. 2).

### The effect of controlled drought on physiological behavior

The drought treatment was applied during the last 39 days of the experiment, coinciding with the inflorescence-development phase (Phase III). Throughout this period, all drought-treated plants exhibited a significant reduction in whole-plant daily transpiration, as compared to the control group, with each treatment group showing distinct and statistically significant differences. This pattern was observed under both controlled conditions (growth chamber; Fig. 3a) and non-controlled conditions (commercial greenhouse; Fig. 3b). The daily transpiration patterns in the non-controlled greenhouse were notably more erratic than those observed in the controlled growth chamber. This variability was likely driven by fluctuating environmental factors such as temperature, relative humidity, and VPD (Fig. S1). Despite their similar size in the two environments (Table S3), the average daily transpiration of ‘Odem’ plants was higher in the greenhouse than it was under controlled conditions. To better understand the physiological adjustments observed under controlled drought conditions, we examined the plants’ momentary water-balance responses, focusing on their dynamic interactions with environmental variables.

**Fig. 3.**
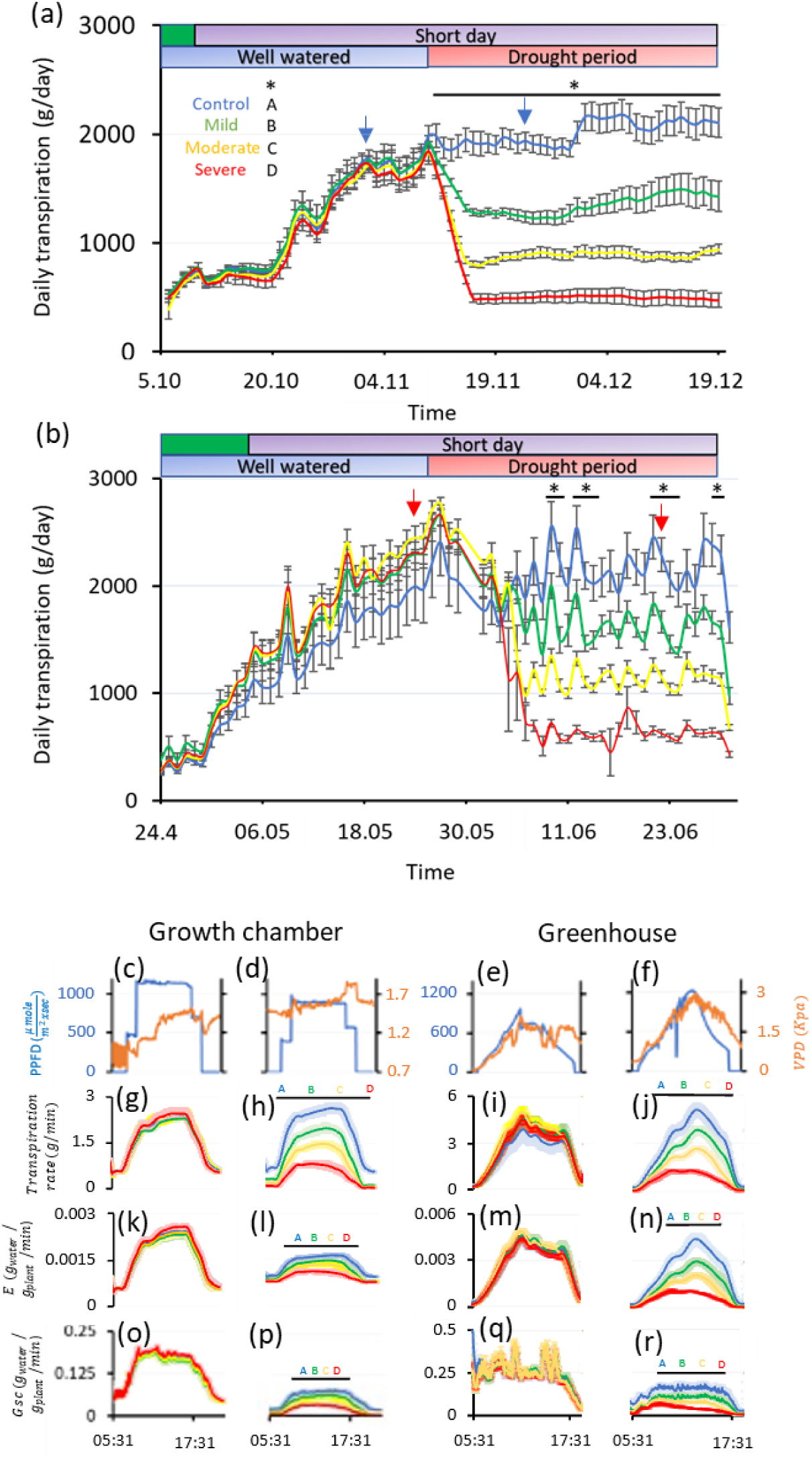
Physiological response of ‘Odem’ to different treatments. Means ± SE of daily transpiration are presented for ‘Odem’ plants under (a) controlled and (b) non-controlled atmospheric conditions across representative experiments (Experiments 1 and 4, Table 1). At the top of the graphs, the upper bar indicates the photoperiod (green for vegetative growth, purple for reproductive growth) and the lower bar indicates the drought period (blue for well-watered, red for drought). Different colors represent the different irrigation levels: blue for well-watered (control), green for mild drought, yellow for moderate drought, and red for severe drought. Panels c–f show the continuous daily plant-environment interactions with photosynthetic photon flux density (PPFD, blue) and vapor pressure deficit (VPD, orange) under well-watered and drought conditions on representative days (indicated by the blue and red arrows in a and b, respectively). (g–j) Mean ± SE transpiration rate (TR), (k–n) normalized transpiration (E), and (o–r) canopy stomatal conductance (Gsc) on the same representative days. Black solid lines denote significant differences, as determined using Tukey’s HSD test (p < 0.05; 5 ≤ n ≤ 6).

### Momentary water-balance responses

We examined momentary physiological traits on two representative days (marked with blue and red arrows in Fig. 3a, b). Under all irrigation conditions in both environments, both the whole-plant transpiration rate and normalized transpiration (E) were closely linked with light intensity (PAR). Notably, plants grown in the greenhouse under well-watered conditions exhibited greater transpiration than those grown in the growth chamber (Fig. 3g, k and Fig. 3i, m, respectively). The canopy stomatal conductance (Gsc) exhibited a similar response to PAR but showed greater sensitivity to instantaneous fluctuations in VPD (Fig. 3o, q). This response highlights the additional atmospheric variability under non-controlled conditions.

### Overall trends in the drought-treated groups

Across all drought treatments, physiological traits exhibited consistent patterns. Signals of limited water availability predominated over the enhancing effects of atmospheric factors, resulting in significantly lower daily integral values for all measured traits in both the controlled and non-controlled environments (Fig. 3h, l, p and Fig. 3j, n, r, respectively).

### Cumulative effects of drought treatment

The drought treatments induced cumulative physiological, anatomical, and yield effects across the four irrigation treatments. These effects will be further analyzed and discussed in relation to transpiration, biomass allocation, and secondary-metabolite production.

### The effect of controlled drought on the total fresh and dry weights of different plant organs

The controlled drought treatments led to significant reductions in calculated plant weight, as observed across the four levels of drought (Fig. 4a). The strong correlation between the harvested fresh weights and oven-dry weights of these plants (*r*^2^ = 0.99, *p* = 0.002; Fig. 4b) validated the precision of the calculated plant weights and underscored the substantial impact of drought on plant biomass. Interestingly, among the well irrigated plants, the biomass gain (calculated plant weight) at the end of the vegetative Phase II (vertical dashed red line; Fig. 4a) was strongly correlated with the oven-dry inflorescence weight (*r*^2^ = 0.54, *p* = 0.001; Fig. 4c). Drought during Phase III had a particularly pronounced influence on inflorescence weight, as indicated by the strong correlation between the dry inflorescence weight and the measured total fresh weight at the experiment’s conclusion (*r*^2^ = 0.98, *p* < 0.0001; Fig. 4d).

**Fig. 4.**
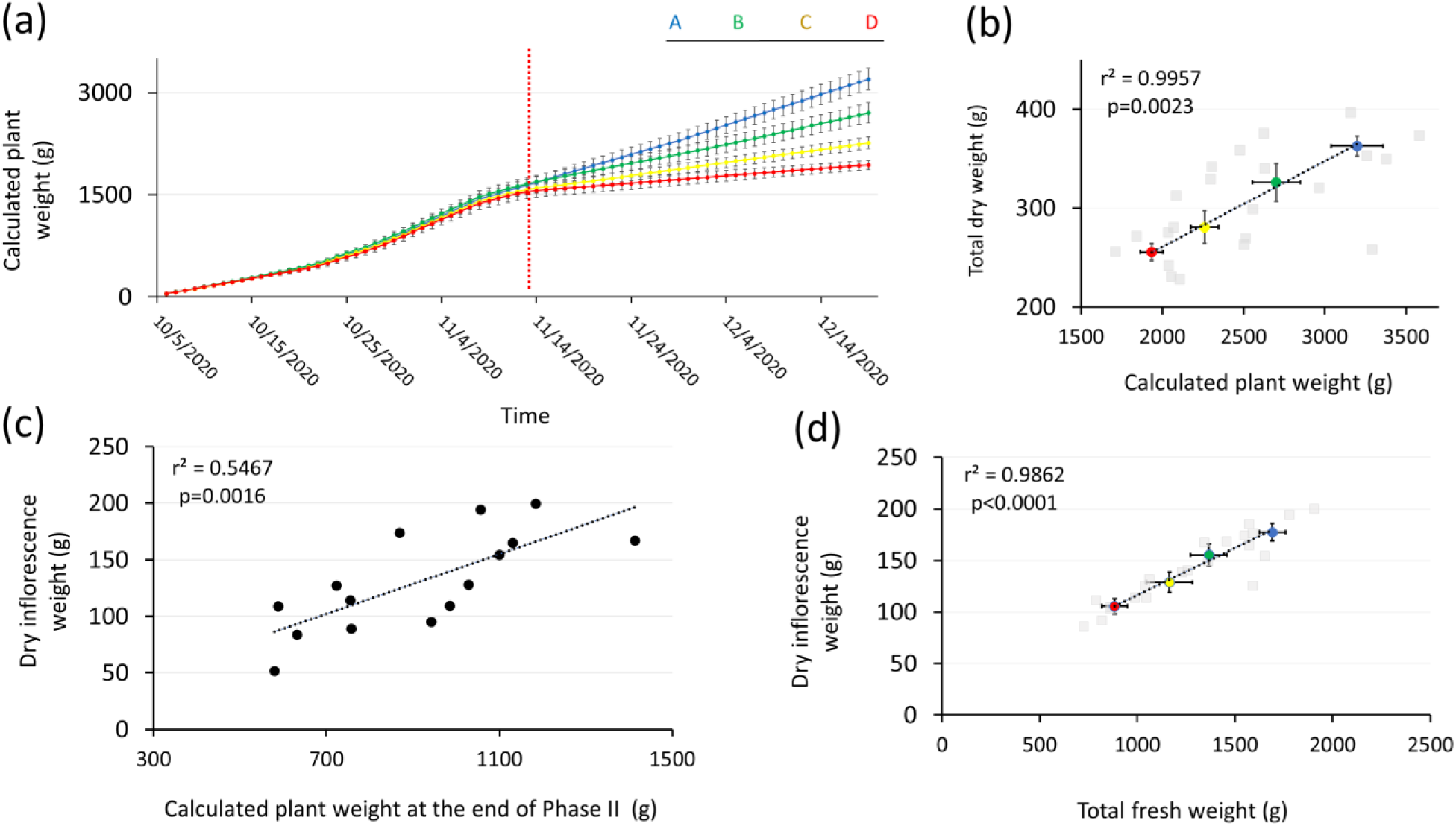
Impact of controlled drought treatments on biomass and correlations between different biomass components in ‘Odem’. (a) Means ± SE of calculated plant weight throughout the experiment (data from Experiment 1, Table 1), in which drought treatments led to a significant reduction in biomass compared to control (colors correspond to those used in Fig. 1). The vertical dashed red line marks the transition to drought treatments following Phase II. Different letters above bars indicate significant differences based on Tukey HSD analysis (p ≤ 0.05; 5 ≤ N ≤ 6). (b) There was a strong positive correlation (r^2^ = 0.99, p = 0.002) between total oven-dry weight and calculated plant weight across treatments (Data from experiment 1, table 1). (c) There was also a significant correlation (r^2^ = 0.54, p = 0.001) between dry inflorescence weight and fresh plant weight at the end of Phase II under control conditions (N = 15). (d) There was a strong correlation (r^2^ = 0.98, p < 0.0001) between dry inflorescence weight and total fresh weight at the experiment’s conclusion, emphasizing the direct effect of drought on inflorescence biomass (data from experiment 1, table 1).

### Organ-specific responses to drought

Further analysis of the oven-dry weights of three different plant organs revealed a differential response to drought treatments on experiment 1 ‘Odem cultivar’. A significant reduction in inflorescence dry weight, relative to the well-watered control, was observed for both the medium and severe drought treatments (Fig. 5a). A similar trend was observed for both leaf dry weight (Fig. 5b) and total oven-dry weight (Fig. 5c). However, stem dry weight did not vary significantly across the treatments (Fig. 5d). These patterns were consistent across replicated experiments and similar trials (Table S3). Stem dry weight accounted for 15% of the total dry biomass (Fig. 5e) and was unaffected by drought (Fig. 5b). Yet, increased drought severity was associated with an increase in the relative proportion of the stem weight to 20% of the total dry biomass in the severe drought treatment (Fig. 5h). Conversely, the inflorescence proportion was smaller under milder drought conditions, reflecting a trade-off between biomass allocation to reproductive versus structural tissues (Fig. 5e–h). We found a strong relationship between the oven-dry weights of the organs and cumulative transpiration, specifically, inflorescence dry weight (*r*^2^ = 0.98, *p* = 0.0005; Fig. 5j), leaf dry weight (*r*^2^ = 0.96, *p* = 0.0028; Fig. 5k), and total dry weight (*r*^2^ = 0.99, *p* = 0.0005; Fig. 5l). Further analysis revealed strong correlations between the dry weights of different organs: inflorescence dry weight and leaf dry weight (*r*^2^ = 0.92, *p* < 0.0001; Fig. 5m), inflorescence dry weight and total dry weight (*r*^2^ = 0.99, *p* < 0.0001; Fig. 5n), and leaf dry weight and total dry weight (*r*^2^ = 0.96, *p* < 0.0001; Fig. 5o). These trends were consistent with findings from ‘Odem’ and ‘187’ plants grown under non-controlled atmospheric conditions, as observed in Experiments 5 and 6 (Fig. S3a–f, Fig. S3g–l, and Table S3).

**Fig. 5.**
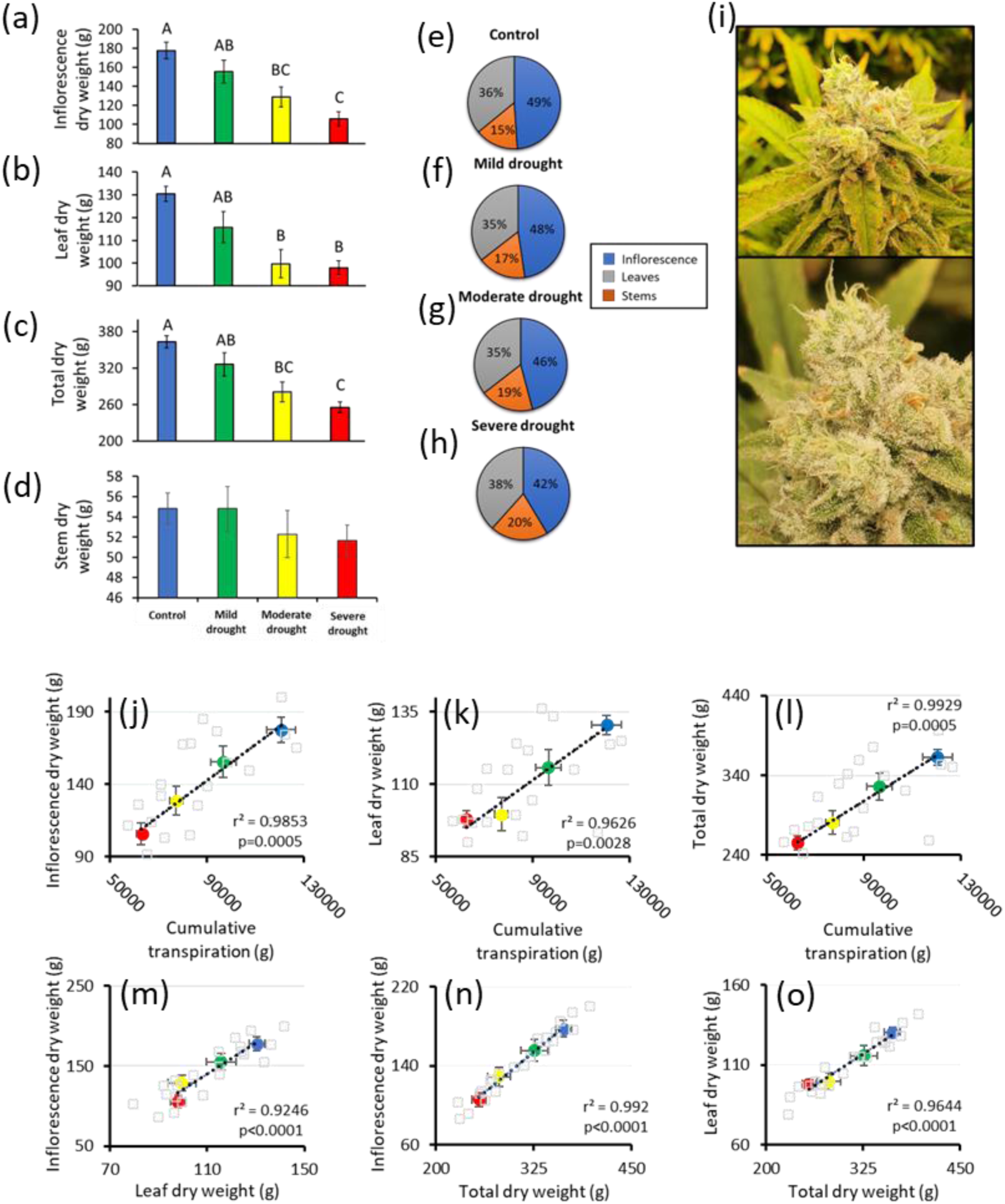
Effects of controlled drought treatments on dry biomass composition of ‘Odem’. Mean ± SE values of oven-dry biomass for (a) inflorescences, (b) leaves, (c) total shoots, and (d) stems under four irrigation treatments (colors as in Fig. 1). Drought treatments led to significant reductions in the dry biomass of inflorescences, leaves, and total shoots, but not stems. Different letters above bars indicate significant differences, as determined using Tukey’s HSD test (p ≤ 0.05; 5 ≤ N ≤6). Relative composition of dry biomass components under (e) control conditions, (f) low drought, (g) medium drought, and (h) severe drought. Drought severity decreased the relative proportion of the inflorescences, while increasing the relative proportion of the stems. (i) Representative images of primary inflorescences during mid and late Phase III (top and bottom, respectively). Pearson correlations between cumulative transpiration and dry weights of (j) inflorescence, (k) leaves, and (l) total shoot, demonstrating strong positive associations. Strong, positive correlations were observed among the weights of the different components of the dry biomass: (m) inflorescences and leaves, (n) inflorescences and total shoots, and (o) leaves and total shoots (data from experiment 1, table 1). Correlation p-values refer to multivariate coefficient probability tests.

### The effect of controlled drought on concentrations of secondary metabolites

To investigate the effects of controlled drought on secondary metabolites, we analyzed phytocannabinoids and terpenoids extracted from the primary and secondary inflorescences of two Chemotype-I cultivars, ‘Odem’ (Figs. 6, 8; Tables S4–S8, S15–S19) and ‘MVA’ (Figs. 7, 9; Tables S9–S10, S20–S21), grown under controlled atmospheric conditions.

**Fig. 6.**
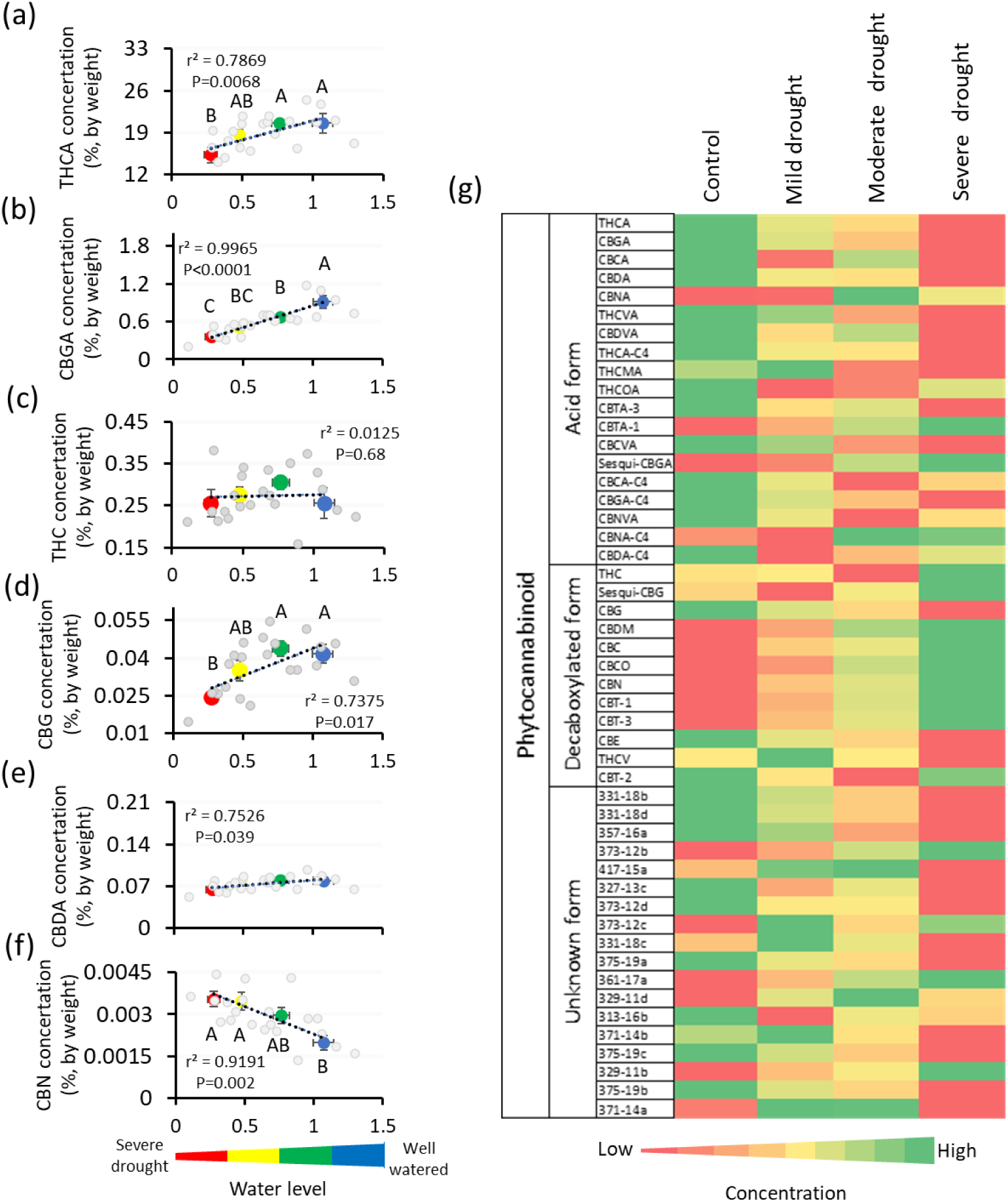
Influence of controlled drought on phytocannabinoid concentrations in ‘Odem’. Correlations of mean ± SE concentrations of major phytocannabinoids: (a) THCA, (b) THC, (c) CBGA, (d) CBG, (e) CBDA, and (f) CBN with the level of drought (colors as indicated in Fig. 1; data from the representative Experiment 1). Different letters indicate significant differences, as determined using Tukey’s HSD test (p < 0.05, 5 ≤ N ≤ 6). The squared value of the Pearson correlation coefficient was derived from regression-fit analysis of the group means. The correlation p-value is based on the multivariate coefficient probability test for all plants. (g) A heatmap showing the grand mean phytocannabinoid concentrations from two independent experiment replications of the ‘Odem’ cultivar (Experiments 1 and 2; Tables S4– S5 and Tables S6–S8, respectively). Green signifies high concentrations; red indicates low concentrations.

**Fig. 7.**
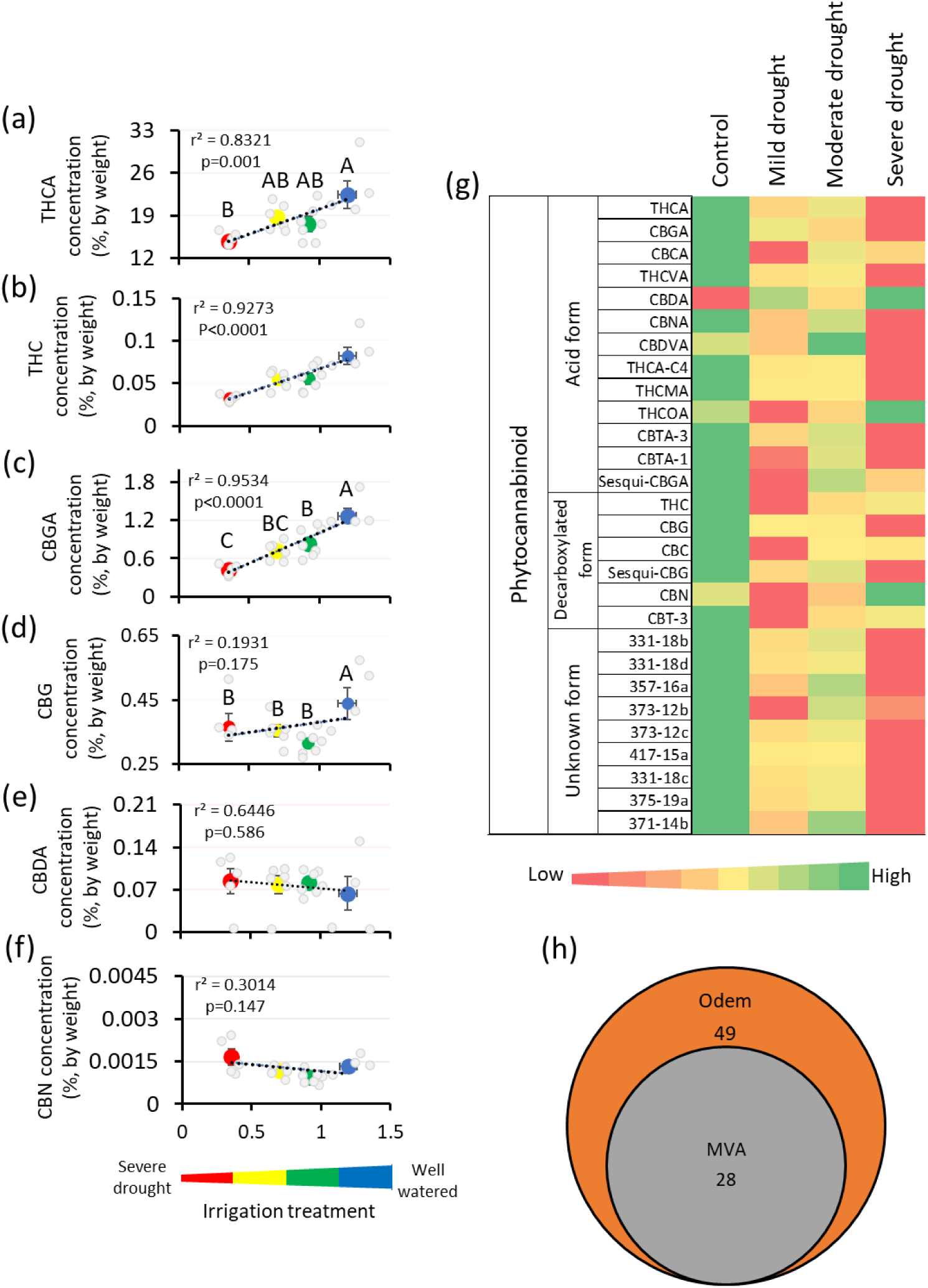
Influence of controlled drought on concentrations of major phytocannabinoids in ‘MVA’. The graph depicts correlations of mean ± SE concentrations of phytocannabinoids: (a) THCA, (b) THC, (c) CBGA, (d) CBG, (e) CBDA, and (f) CBN with the different drought treatments (colors as described in Fig. 1) for the ‘MVA’ cultivar (Experiment 3). Different letters above the data points indicate significant differences, as determined using Tukey’s HSD test (p < 0.05, 5 ≤ n ≤ 6). The squared Pearson correlation coefficient is from the regression-fit analysis for the group means. The correlation p-value is derived from the multivariate coefficient probability test for all plants. (g) A heatmap showing the mean phytocannabinoid concentrations in the ‘MVA’ cultivar (Experiment 3; Tables S9–S10). (h) A Venn diagram comparing ‘Odem’ and ‘MVA’ indicates that 28 phytocannabinoids are common to both cultivars and that ‘Odem’ has an additional 21 phytocannabinoids.

**Fig. 8.**
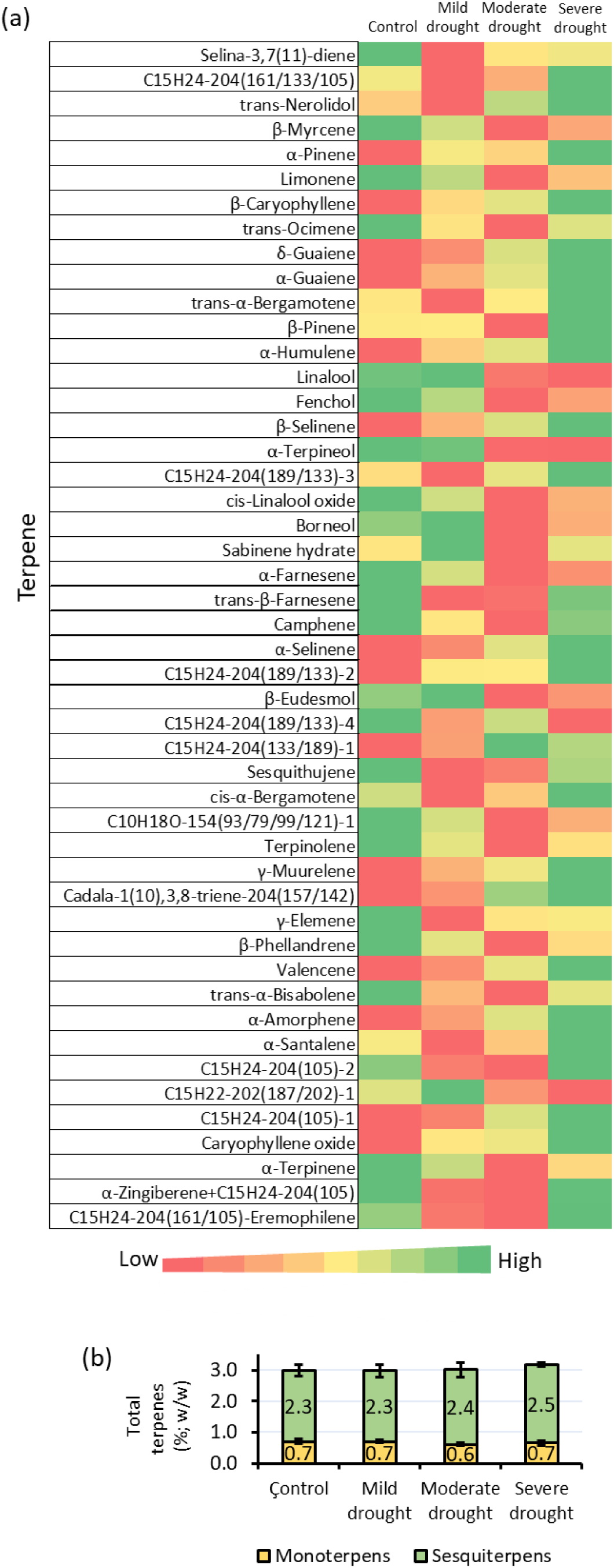
Impact of controlled drought on terpene concentrations (ppm) in ‘Odem’. (a) A heatmap displaying the grand mean concentrations of the top-50 most abundant terpenes across two replicated experiments (Experiments 1 and 2; Tables S15–S16 and S17–S19, respectively). (b) Mean ± SE concentrations of total terpenes, monoterpenes, and sesquiterpenes in the various drought treatments. Different letters above the bars indicate significant differences, as determined using Tukey’s HSD test (p < 0.05, 5 ≤ n ≤ 6).

**Fig. 9.**
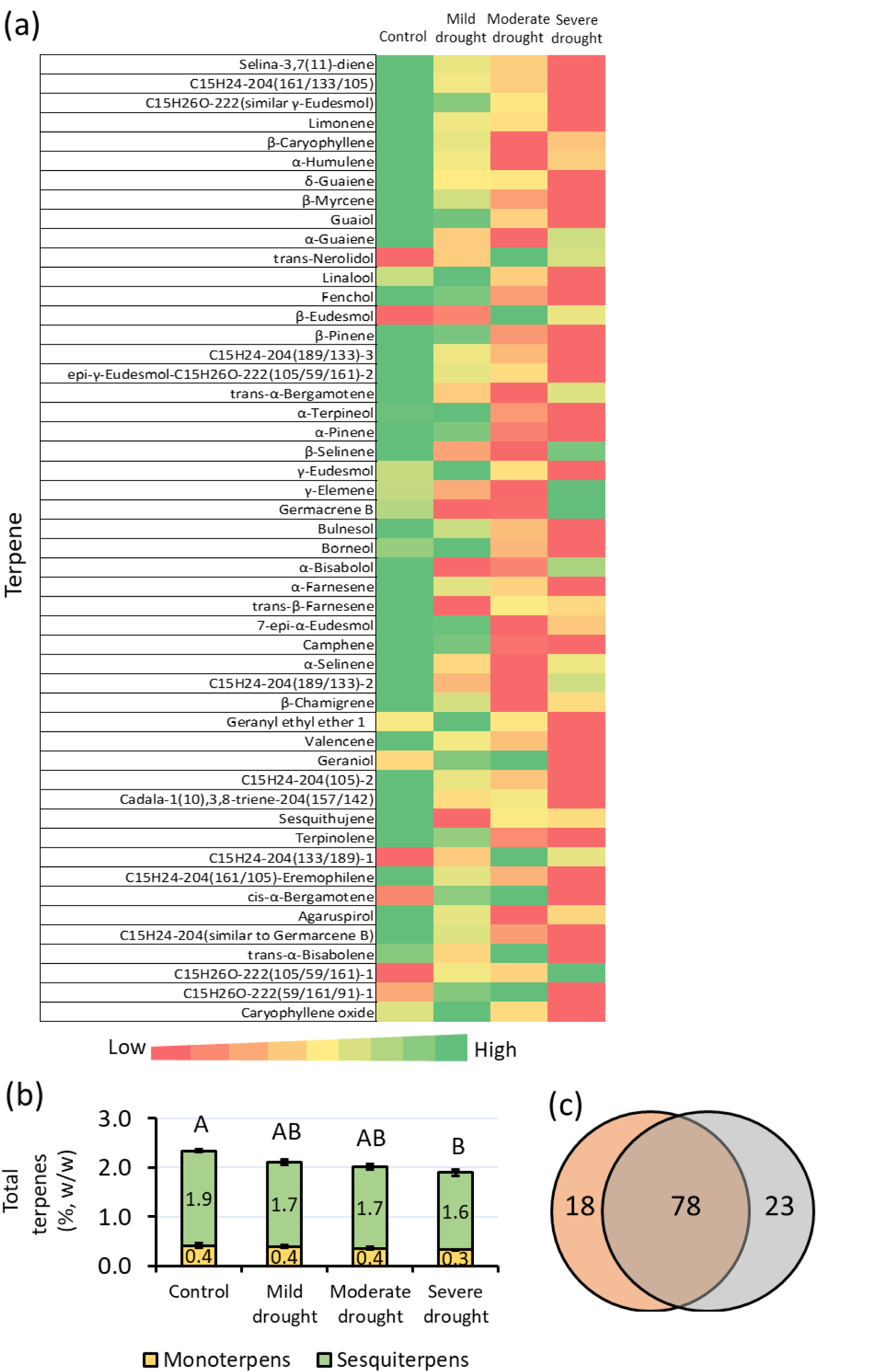
Effect of controlled drought on terpene concentrations (ppm) in ‘MVA’. (a) Heatmap of the means of the top-50 most abundant terpenes in terms of their concentrations in the ‘MVA’ cultivar (Experiment 3, Tables S20–S21). (b) Total terpene, monoterpene, and sesquiterpene concentrations in the different drought treatments. Different letters represent significant differences, according to Tukey’s HSD test (p < 0.05, 5≥ N ≥6). (c) Venn diagram of the terpene distributions of ‘Odem’ (orange) and ‘MVA’ (gray), indicating that 78 phytocannabinoids are common to both cultivars, while ‘Odem’ has an additional 18 unique terpenes and ‘MVA’ has an additional 23 unique terpenes.

### Phytocannabinoid response in ‘Odem’

The concentration of THCA in the ‘Odem’ plants was significantly lower under severe drought conditions and was strongly correlated with the level of irrigation (*r*^2^ = 0.78, *p* = 0.006; Fig. 6a). Similarly, levels of CBGA, the direct precursor of THCA and CBDA, were significantly lower in the moderate and severe drought treatments and also strongly correlated with the level of irrigation (*r*^2^ = 0.99, *p* < 0.0001; Fig. 6b).

The decarboxylated forms of these compounds exhibited varied responses. The THC concentration did not vary across the treatments (Fig. 6c). Concentrations of CBG were significantly lower under severe drought, correlating strongly with the level of irrigation (*r*^2^ = 0.73, *p* = 0.017; Fig. 6d). Conversely, CBN concentrations increased significantly under drought conditions and were strongly correlated with irrigation level (*r*^2^ = 0.92, *p* = 0.002; Fig. 6f). CBDA concentrations did not vary significantly across the different treatments (Fig. 6e). Significant differences in absolute concentrations of several minor phytocannabinoids under drought stress were observed in the ‘Odem’ plants (Tables S4–S8).

Heatmaps summarizing phytocannabinoid concentrations from two independent experiments (Experiments 1 and 2) indicated a general decrease in most acidic phytocannabinoids in response to drought. Exceptions included cannabinolic acid (CBNA), cannabitriolic acid (CBTA-1), and sesqui-CBGA, which exhibited increased concentrations under drought conditions. Additionally, a consistent decreasing trend was observed for major decarboxylated phytocannabinoids as drought severity increased (Fig. 6g). Interestingly, levels of compounds such as sesqui-CBGA, CBN, CBNA, CBTA-1, cannabicitran-1 (CBT-1), and cannabicitran-3 (CBT-3) were higher in the drought-treated plants than they were in the control plants (Fig. 6g).

### Phytocannabinoid response in ‘MVA’

Complementary findings were observed for the ‘MVA’ cultivar (Experiment 3). Phytocannabinoid analysis (Tables S9–S10) revealed strong correlations between irrigation treatment and concentrations of THCA (*r*^2^ = 0.83, *p* = 0.001; Fig. 7a), THC (*r*^2^ = 0.92, *p* < 0.0001; Fig. 7b), and CBGA (*r*^2^ = 0.95, *p* < 0.0001; Fig. 7c). Significantly different concentrations of THCA were observed in the control and severe-drought treatments (Fig. 7a) and significantly different levels of CBGA were observed across all drought levels, with levels in the plants subjected to the severe drought differing significantly from those observed for the mild and moderate drought treatments (Fig. 7c). CBG concentrations across all of the drought treatments were also significantly lower than those observed in the control (Fig. 7d). In contrast, THC levels were unaffected (Fig. 7b) and CBDA levels exhibited no significant drought response (*r*^2^ = 0.64, *p* = 0.5; Fig. 7e). Unlike the situation in ‘Odem’, the CBN concentration in ‘MVA’ was not affected by drought stress. Significant variations were also observed in the absolute concentrations of minor phytocannabinoids (Tables S9–S10).

Heatmaps of relative phytocannabinoid concentrations in ‘MVA’ revealed a general decrease in concentrations in response to drought (Fig. 7g). A Venn diagram comparison showed that the ‘Odem’ cultivar had 49 unique phytocannabinoids and that ‘MVA’ contained 28 unique phytocannabinoids, with 28 phytocannabinoids found in both cultivars (Fig. 7h). These findings were consistent with an independent-repetition experiment conducted using the ‘Odem’ cultivar under controlled conditions (Tables S6–S8) and with phytocannabinoid analyses conducted during the middle of the inflorescence-development phase (Table S6). Comparable results were also obtained under non-controlled atmospheric conditions for ‘Odem’ (Experiment 4; Tables S11–S12) and ‘187’ (Experiment 5; Tables S13–S14).

### Effects of drought on terpenoid levels in ‘Odem’ and ‘MVA’

To investigate the impact of drought on terpenoid levels, heatmaps of the 50 major terpenoids were generated for both ‘Odem’ and ‘MVA’ grown under controlled conditions (Figs. 8 and 9, respectively; Tables S15–S19 and S20–S21, respectively).

In ‘Odem’, no clear pattern of terpenoid response to drought was observed. The concentrations of some terpenoids increased in response to drought conditions, while others decreased (Fig. 8a). Total relative terpenoid concentrations remained constant across the treatments, with no significant differences observed (Fig. 8b). In contrast, in ‘MVA’, there was a consistent decrease in the concentrations of nearly all of the analyzed terpenoids under drought conditions (Fig. 9a). This led to significantly lower total terpenoid contents in the plants subjected to severe drought, as compared to the control plants (Fig. 9b).

Analysis of terpenoid distribution revealed that ‘Odem’ contained 96 unique terpenoids; whereas ‘MVA’ contained 101 unique terpenoids, with 78 terpenoids found in both cultivars (Fig. 9c). Consistent trends were observed in an independent-repetition experiment conducted with the ‘Odem’ cultivar (Tables S17–S19). Terpenoid analyses conducted during the middle of the inflorescence-development phase mirrored these findings (Table S17).

## Discussion

Informed by prior studies, we hypothesized that controlled drought treatments would reduce inflorescence yields while increasing concentrations of secondary metabolites (i.e., phytocannabinoids and terpenoids) under both controlled and non-controlled atmospheric conditions. Our findings confirmed the first part of the hypothesis, demonstrating a decrease in inflorescence yield as drought severity increased (Fig. 5a, j; Fig. S3a, g). However, our results did not support the second part of the hypothesis. Contrary to recent reports, the concentrations of the majority of acid-form phytocannabinoids, including THCA and CBGA, decreased in concentration in response to drought stress in two independent cultivars (Figs. 6 and 7; Tables S4–S10). In contrast, terpene concentrations exhibited a more complex and cultivar-specific response (Figs. 8 and 9; Tables S15–S21).

The discrepancy between our findings and those reported by Caplan et al. (2019), which formed a central part of our rationale, can likely be attributed to significant differences in experimental conditions, specifically radiation levels, the drought-regulation methodology used, and suboptimal growing conditions. In terms of radiation levels, Caplan et al. employed radiation levels that were lower than the ideal intensities recommended for cannabis growth (Chandra et al., 2008). Lower light intensities would be expected to be associated with lower transpiration rates, potentially mitigating the effects of drought stress. In our study, the higher radiation levels aligned with optimal growth conditions, ensuring measurable and reproducible drought-induced effects. In terms of drought-regulation methodology, the drought treatments used by Caplan et al. were manually applied and regulated through a low-throughput system. This approach introduces challenges in maintaining standardized and controlled drought stress across treatments and repetitions, compromising reproducibility (Gosa et al., 2019; Moshelion et al., 2024). Finally, the low yields reported by Caplan et al. for their well-irrigated control plants suggest that these plants may not have been grown under optimal conditions. Such suboptimal growth makes it difficult to establish reliable stress-control comparisons, as baseline productivity and physiological responses are not maximized.

In contrast, our study leveraged a high-throughput, automated drought-regulation system (the PlantArray platform), which provided precise, reproducible, and physiologically relevant drought treatments tailored to each plant’s performance. This enabled us to maintain a consistent drought-stress gradient across all experiments, eliminating variability caused by manual irrigation inconsistencies. Furthermore, the use of optimal radiation levels and rigorous environmental monitoring ensured that both control and drought-treated plants were grown under ideal baseline conditions, enabling a more accurate assessment of the impact of the examined drought treatments on yield and metabolite profiles.

The lower levels of acid-form phytocannabinoids observed under drought stress in our study align with the hypothesis that drought limits primary photosynthetic processes and the availability of metabolic energy, which are critical for synthesizing these metabolites. The cultivar-specific responses observed in terpenoid concentrations further highlight the complexity of secondary-metabolism regulation under drought conditions, as terpene biosynthesis is influenced by both genetic and environmental factors (Russo, 2011; Burgel et al., 2020). Our results underscore the importance of accounting for both experimental precision and environmental factors when interpreting drought-induced responses in cannabis and other crops.

### Cannabis growth phases and water-management behavior

Our findings indicate that the cannabis life cycle is more accurately described by three distinct physiological phases rather than the two phases commonly cited in the literature (Mediavilla et al., 1998; Caplan et al., 2017; Bevan et al., 2021). Phase I is a vegetative stage characterized by long-day photoperiods, vigorous growth, and constant rates of increase in key physiological traits, such as transpiration and biomass accumulation. Phase II is a transitional stage that is triggered by a shift to short-day photoperiods. Rapid growth and high transpiration rates are sustained during Phase II, despite a 25% reduction in photosynthetic radiation time. Phase III is the inflorescence-development stage during which transpiration stabilizes, while the rates of height gain and new leaf emergence decrease dramatically. However, biomass accumulation continues during Phase III, driven primarily by inflorescence development. This phase culminates in a significant reduction in WUE to 0.006 ± 0.0006 g plant / g water (Fig. 2).

These results highlight a sink-source behavior typical of seasonal, terminal flowering species. Comparable growth phases, including stabilization of transpiration and leaf-emergence rates, have been reported in potato (*Solanum tuberosum*), another seasonal, terminal flowering plant (Almekinders and Struik, 1996; Nelson and Hwang, 2008). Understanding these distinct growth phases and their associated physiological traits can provide critical insights for both future physiological studies and the commercial cultivation of cannabis. Specifically, managing irrigation, nutrient supply, and environmental controls in line with the demands of each phase may optimize plant productivity and resource-use efficiency.

### Physiological water-management response under drought conditions

The drought treatments were applied during Phase III, to target inflorescence development with minimal impact on vegetative growth. At the start of Phase III, the plants in all treatment groups were of similar sizes, with similar numbers of leaves per plant and similar levels of transpiration. Plants subjected to drought treatment displayed a reduction in daily transpiration proportional to the severity of the drought stress under both controlled and non-controlled atmospheric conditions (Fig. 3a, b). This reduction aligns with the general behavior of crops transitioning from productive to protective modes under drought conditions, highlighting the necessity of using standardized treatments for reliable results (Moshelion, 2020; Moshelion et al., 2024). The use of automatic-feedback irrigation achieved this standardization by exposing all plants within a treatment group to equivalent stress levels. Under non-controlled conditions, fluctuations in physiological behavior were observed, likely resulting from continuous stomatal adjustments to optimize water balance in response to environmental variations, such as fluctuations in light intensity and VPD (Lewis and Went, 1945). Interestingly, agronomic WUE remained consistent across atmospheric conditions and improved with increasing drought severity (Table S3). While drought is known to modify WUE (Gago et al., 2014; Kang et al., 2017; Liu et al., 2017), this does not necessarily translate into improved yield. In high-yielding plants, drought can act as a counteractive factor (Moshelion, 2020).

These findings suggest that to mitigate the detrimental effects of abiotic drought stress on medical cannabis yield, intelligent irrigation systems tailored to the plant’s real-time physiological needs are essential. Such systems can optimize resource management and preserve productivity under water-limiting conditions.

### Biomass yield response of cannabis under well-watered and drought conditions

Continuous, non-destructive monitoring of plant biomass is critical for assessing the impact of drought, as compared to control conditions(Halperin et al., 2017). The strong correlation between the weights calculated by the platform and the measured dry biomass (*r*^2^ = 0.99, *P* = 0.0023; Fig. 4b) confirms the precision of this non-destructive approach. Our monitoring revealed that the reduction in total biomass under drought conditions during Phase III was primarily due to a decline in inflorescence biomass (Fig. 5e–h). Notably, no significant differences in biomass gain were observed before the onset of the drought stress (Fig. 4a). By this stage, the leaves and stems had largely reached their final developmental states and those organs remained unaffected by drought (Fig. 2a).

The strong correlation between shoot fresh weight and oven-dry inflorescence weight at the experiment’s conclusion (Fig. 4d and Fig. 5n) suggests that yield potential is largely determined by the end of the transitional growth phase. Drought stress subsequently reduces reproductive yield by directly impacting inflorescence development (Fig. 5a, j; Fig. S3a, g; Table S3). A similar dynamic has been observed in maize (*Zea mays*), in which yield potential is established during the pre-flowering vegetative phase (Jacobs and Pearson, 1991). Our findings further indicate that vegetative biomass at the end of Phase II serves as a significant predictor of final inflorescence weight (*r*^2^ = 0.54, *P* = 0.0016; Fig. 4c). Early vigor during growth stages correlated with higher yields in well-irrigated plants, consistent with patterns observed across various crops(Bullock et al., 1998; Peltonen-Sainio et al., 2006; Gosa et al., 2022). This suggests that early-stage vigor can shorten the time frame required for yield-optimization studies.

The similarities in plant size and transpiration levels prior to the onset of drought support the hypothesis that stomatal closure and the subsequent decrease in stomatal conductance of CO_2_ (Lawson and Blatt, 2014) trigger yield loss. This phenomenon has also been reported in other crops(McClelland, 1968; Cassman et al., 1989). The strong correlation between inflorescence dry weight and leaf dry weight (*r*^2^ = 0.98, *P* < 0.0001; Fig. 5m) suggests that reduced photosynthesis is the primary driver of this biomass loss.

Greater plant transpiration is closely linked with elevated CO_2_ assimilation and sugar production (Ibrahim and Jaafar, 2012; Kumar et al., 2017), with sugars subsequently transported to reproductive tissues to support yield production (Rayner, 1969; Lemoine et al., 2013; Julius et al., 2017; Medina et al., 2019). The clear correlation between transpiration and yield (Fig. 5j–l; Fig. S3a, g; Table S3) underscores the substantial impact of even minimal reductions in soil moisture on yield. To maximize yields in medical cannabis cultivation, it is crucial to avoid abiotic drought stress during inflorescence development. Our findings highlight the need for advanced smart irrigation systems that dynamically respond to the plant’s real-time physiological needs, to optimize water use and mitigate yield loss.

### Drought-response profile of secondary metabolites

Phytocannabinoids were categorized into acidic and decarboxylated forms. In the primary inflorescence, the concentrations of most acid-form phytocannabinoids, including THCA and CBGA, decreased with increasing drought severity in both ‘Odem’ and ‘MVA’ (Figs. 6a, b, 7a, c). These results were consistent across all of our experiments (Tables S4–S14). Acid-form phytocannabinoids are naturally synthesized in the glandular trichomes of the cannabis inflorescence during development (Schuurink and Tissier, 2020). These compounds degrade over time and are decarboxylated. Under drought conditions, we observed an upward trend in levels of decarboxylated phytocannabinoids, such as CBN (Fig. 6f) and cannabidiol monoethyl ether (CBDM) (Fig. 6g), suggesting active degradation during drought stress. However, degradation alone cannot explain the dramatic reduction in major phytocannabinoids, particularly THCA, since its primary degradation products, THC and CBN, were detected at much lower concentrations, as compared to control THCA levels (20– 2000 times less; Figs. 6a, f and 7a, f).

Thus, we propose that drought stress triggers two mechanisms that lead to reduced levels of major phytocannabinoids: reduced biosynthesis and degradation. The reduced biosynthesis involves a decline in primary-metabolite production due to reduced stomatal conductance and photosynthesis, which limits the supply of sugar precursors required for phytocannabinoid biosynthesis (via the fatty-acid and MEP pathways; He et al., 2020; Koley et al., 2020; Zhai et al., 2021). This is accompanied by the ongoing degradation of acid-form phytocannabinoids into decarboxylated products under prolonged drought stress.

This trend was also evident in greenhouse experiments conducted under non-controlled conditions, although with greater variability (Tables S11–S14). Phytocannabinoid analyses conducted at the midpoint of the inflorescence-development phase in that environment revealed non-significant similar expression patterns (Table S6), providing further evidence of the continuous detrimental effect of Phase-III drought on the biosynthesis of major phytocannabinoids.

### Terpenoid levels under drought stress

The response of monoterpenoids and sesquiterpenoids to drought stress varied between and within cultivars. In ‘Odem’, the concentrations of some terpenoids increased while others decreased, resulting in no significant change in the total terpene concentration (Fig. 8a, b). These findings were consistent across independent repetitions (Tables S17–S19). This suggests that the decline in phytocannabinoid concentrations in ‘Odem’ may be linked to reduced biosynthesis in the fatty-acid pathway, rather than the MEP pathway, which is shared by both phytocannabinoids and terpenoids (Desaulniers Brousseau et al., 2021). In contrast, in ‘MVA’, we observed a general decrease in terpene concentrations under drought stress (Fig. 9a; Tables S20–S21). In that cultivar, a significant reduction (*p* < 0.05) in total terpene content was observed between the control and severe-drought treatments (Fig. 9b).

Although abiotic stress has been reported to increase terpenoid concentrations in cannabis (Namdar et al., 2019) and other plants (Selmar and Kleinwächter, 2013), the functional significance of such changes remains unclear. Given that both ‘Odem’ and ‘MVA’ are wind-pollinated, these variations are unlikely to serve as signals for pollinator attraction (Small and Antle, 2008; O’Brien and Arathi, 2019). Instead, the expression of certain terpenes may provide protection from pests and diseases (Singh and Sharma, 2015; Aljbory and Chen, 2018; Toffolatti et al., 2021), offering an evolutionary rationale for the observed responses. Our results support this hypothesis, while also suggesting that the cultivar-specific variability in terpenoid response to drought may reflect the cultivars’ distinct genetic origins.

For the ‘MVA’ cultivar, it is plausible that drought stress inhibits both the MEP and MEV pathways, reducing the levels of sugar precursors essential for terpene biosynthesis. Given the potential aromatic and commercial importance of terpenoid profiles (Plumb et al., 2022), targeted irrigation protocols may be developed to achieve desired terpene responses in specific cannabis cultivars.

## Conclusions

We have identified a distinct transition phase (Phase II) in the growth cycle of *Cannabis sativa*, which exhibits physiological patterns similar to the vegetative phase (Phase I) despite occurring under reproductive conditions (Phase III). The early vigor of plants during Phase II is closely linked with key physiological parameters, including daily transpiration, normalized transpiration (E), and canopy stomatal conductance (Gsc). These traits strongly correlate with reproductive traits during inflorescence production in Phase III.

Our findings demonstrate that plant biomass at the end of Phase II serves as a significant determinant of Phase-III productivity. However, any reduction in soil water availability that limits transpiration will directly impair flower weight and secondary-metabolite production, irrespective of initial plant biomass. Unlike previous reports and some common understandings of growers in this field, to optimize yield and metabolite content, it is essential to prevent stress by maintain non-limiting transpiration through appropriate irrigation practices throughout the plant’s life cycle. Even minimal drought stress results in a significant decline in major phytocannabinoids, particularly CBGA and THCA, as well as inflorescence biomass. The terpenoid response to controlled multilevel drought stress is cultivar-specific and non-uniform. This response likely reflects the commercial preference for drought-induced modifications in the terpene profile, particularly in the recreational market, in which aromatic traits are valued by growers and consumers. However, since drought stress negatively impacts inflorescence size and phytocannabinoid content, cultivation practices should be designed to avoid abiotic stress during the inflorescence-development phase. Advanced irrigation systems, tailored to the plant’s real-time physiological needs, are essential to maximize yield and maintain the quality of secondary metabolites.

## Acknowledgements

This research was supported in part by the Israel Science Foundation, grant no. 1043/20. We extend our sincere gratitude to the workers at BOL Pharma—Lior Hadad, Orit Gazovitch, Hen Navon, and Shauli Avraham—for their unique expertise, practical knowledge, and invaluable support throughout the study.

## Competing Interests

None declared.

## Author Contributions

**IS**: Designed and performed the research, analyzed the data, formulated the hypotheses, and co-wrote the manuscript with **MM**.

**II**: Contributed to the experiment set-up and provided feedback and support regarding sensor usage.

**OB**: Assisted in conducting the experiment and performing post-harvest analysis.

**OG**: Managed and conducted the quantification of the secondary metabolites.

**ZK**: Advised on secondary-metabolite analysis, evaluated hypotheses, and reviewed the manuscript.

**DT**: Supported commercial farm operations and seedling preparation for all experimental needs.

**DM**: Managed secondary-metabolite analysis and reviewed the manuscript.

**MM**: Principal investigator who oversaw the project, formulated the hypotheses and experimental design, conducted data analysis, and co-wrote the manuscript with **IS**.

## Data Availability

The data supporting the findings of this study are available from the corresponding author upon reasonable request. Relevant data are also included in the supplementary material provided with this article.

## Notes

### Competing Interest Statement

The authors have declared no competing interest.

## References

Aljbory Z, Chen MS (2018) Indirect plant defense against insect herbivores: a review. Insect Sci 25: 2–23

Almekinders CJM, Struik PC (1996) Shoot development and flowering in potato (Solanum tuberosum L.). Potato Research 1996 39:4 39: 581–607

Bachir F, Eddouks M, Arahou M, Fekhaoui M (2022) Origin, Early History, Cultivation, and Characteristics of the Traditional Varieties of Moroccan Cannabis sativa L.. Cannabis Cannabinoid Res 7: 603–615

Ben-Shabat S, Fride E, Sheskin T, Tamiri T, Rhee MH, Vogel Z, Bisogno T, De Petrocellis L, Di Marzo V, Mechoulam R (1998) An entourage effect: inactive endogenous fatty acid glycerol esters enhance 2-arachidonoyl-glycerol cannabinoid activity. Eur J Pharmacol 353: 23–31

Berman P, Futoran K, Lewitus GM, Mukha D, Benami M, Shlomi T, Meiri D (2018) A new ESI-LC/MS approach for comprehensive metabolic profiling of phytocannabinoids in Cannabis. Sci Rep. doi: 10.1038/S41598-018-32651-4

Bevan L, Jones M, Zheng Y (2021) Optimisation of Nitrogen, Phosphorus, and Potassium for Soilless Production of Cannabis sativa in the Flowering Stage Using Response Surface Analysis. Front Plant Sci 12: 2587

Borges CV, Minatel IO, Gomez-Gomez HA, Lima GPP (2017) Medicinal plants: Influence of environmental factors on the content of secondary metabolites. Medicinal Plants and Environmental Challenges 259–277

Bullock D, Khan S, Rayburn A (1998) Soybean yield response to narrow rows is largely due to enhanced early growth. Crop Sci 38: 1011–1016

Burgel L, Hartung J, Graeff-Hönninger S (2020) Impact of Different Growing Substrates on Growth, Yield and Cannabinoid Content of Two Cannabis sativa L. Genotypes in a Pot Culture. Horticulturae 2020, Vol 6, Page 62 6: 62

Caplan D, Dixon M, Zheng Y (2019) Increasing inflorescence dry weight and cannabinoid content in medical cannabis using controlled drought stress. HortScience 54: 964–969

Caplan D, Dixon M, Zheng Y (2017) Optimal Rate of Organic Fertilizer during the Flowering Stage for Cannabis Grown in Two Coir-based Substrates. HortScience 52: 1796–1803

Cassman KG, Bryant DC, Higashi SL, Roberts BA, Kerby TA (1989) Soil Potassium Balance and Cumulative Cotton Response to Annual Potassium Additions on a Vermiculitic Soil. Soil Science Society of America Journal 53: 805–812

Cetin O, Bilgel L (2002) Effects of different irrigation methods on shedding and yield of cotton. Agric Water Manag 54: 1–15

Chanda D, Neumann D, Glatz JFC (2019) The endocannabinoid system: Overview of an emerging multi-faceted therapeutic target. Prostaglandins Leukot Essent Fatty Acids 140: 51–56

Chandra S, Lata H, Khan IA, Elsohly MA (2008) Photosynthetic response of Cannabis sativa Photosynthetic response of Cannabis sativa L. to variations in photosynthetic photon flux densities, temperature and CO 2 conditions. Physiol Mol Biol Plants 14: 299–306

Connor JP, Stjepanović D, Le Foll B, Hoch E, Budney AJ, Hall WD (2021) Cannabis use and cannabis use disorder. Nature Reviews Disease Primers 2021 7:1 7: 1–24

Danziger N, Bernstein N (2021a) Plant architecture manipulation increases cannabinoid standardization in ‘drug-type’ medical cannabis. Ind Crops Prod 167: 113528

Danziger N, Bernstein N (2021b) Light matters: Effect of light spectra on cannabinoid profile and plant development of medical cannabis (Cannabis sativa L.). Ind Crops Prod 164: 113351

Desaulniers Brousseau V, Wu B Sen, MacPherson S, Morello V, Lefsrud M (2021) Cannabinoids and Terpenes: How Production of Photo-Protectants Can Be Manipulated to Enhance Cannabis sativa L. Phytochemistry. Front Plant Sci 12: 1035

Gago J, Douthe C, Florez-Sarasa I, Escalona JM, Galmes J, Fernie AR, Flexas J, Medrano H (2014) Opportunities for improving leaf water use efficiency under climate change conditions. Plant Science 226: 108–119

Garan H, Çakalogullari U, İştipliler D, Tatar Ö (2016) IS PROLINE ACCUMULATION UNDER WATER DEFICIT REVERSIBLE IN COTTON? Lucrări Ştiinţifice 59:

Gosa SC, Koch A, Shenhar I, Hirschberg J, Zamir D, Moshelion M (2022) The potential of dynamic physiological traits in young tomato plants to predict field-yield performance. Plant Science 315: 111122

Gosa SC, Lupo Y, Moshelion M (2019) Quantitative and comparative analysis of whole-plant performance for functional physiological traits phenotyping: New tools to support pre-breeding and plant stress physiology studies. Plant Science 282: 49–59

Gülck T, Møller BL (2020) Phytocannabinoids: Origins and Biosynthesis. Trends Plant Sci 25: 985–1004

Halperin O, Gebremedhin A, Wallach R, Moshelion M (2017) High-throughput physiological phenotyping and screening system for the characterization of plant– environment interactions. The Plant Journal 89: 839–850

He M, Qin CX, Wang X, Ding NZ (2020) Plant Unsaturated Fatty Acids: Biosynthesis and Regulation. Front Plant Sci 11: 390

Ibrahim MH, Jaafar HZE (2012) Impact of elevated carbon dioxide on primary, secondary metabolites and antioxidant responses of Eleais guineensis Jacq. (oil palm) seedlings. Molecules 17: 5195–5211

JE J, SJ W, JA B (1999) Marijuana and Medicine: Assessing the Science Base. Marijuana and Medicine. doi: 10.17226/6376

Julius BT, Leach KA, Tran TM, Mertz RA, Braun DM (2017) Sugar Transporters in Plants: New Insights and Discoveries. Plant Cell Physiol 58: 1442–1460

Kalant H (2001) Medicinal use of cannabis: History and current status. Pain Res Manag 6: 80–91

Kang S, Hao X, Du T, Tong L, Su X, Lu H, Li X, Huo Z, Li S, Ding R (2017) Improving agricultural water productivity to ensure food security in China under changing environment: From research to practice. Agric Water Manag 179: 5–17

Koley S, Grafahrend-Belau E, Raorane ML, Junker BH (2020) The mevalonate pathway contributes to monoterpene production in peppermint. bioRxiv 2020.05.29.124016

Koltai H, Namdar D (2020) Cannabis Phytomolecule “Entourage”: From Domestication to Medical Use. Trends Plant Sci 25: 976–984

Kumar U, Quick WP, Barrios M, Cruz PCS, Dingkuhn M (2017) Atmospheric CO2 concentration effects on rice water use and biomass production. PLoS One 12: e0169706

Lawson T, Blatt MR (2014) Stomatal Size, Speed, and Responsiveness Impact on Photosynthesis and Water Use Efficiency. Plant Physiol 164: 1556–1570

Lemoine R, La Camera S, Atanassova R, Dédaldéchamp F, Allario T, Pourtau N, Bonnemain JL, Laloi M, Coutos-Thévenot P, Maurousset L, et al (2013) Source-to-sink transport of sugar and regulation by environmental factors. Front Plant Sci 4: 272

Lewis H, Went FW (1945) Plant Growth Under Controlled Conditions. IV. Response of California Annuals to Photoperiod and Temperature. Am J Bot 32: 1

Liu DL, Zeleke KT, Wang B, Macadam I, Scott F, Martin RJ (2017) Crop residue incorporation can mitigate negative climate change impacts on crop yield and improve water use efficiency in a semiarid environment. European Journal of Agronomy 85: 51– 68

Livingston SJ, Quilichini TD, Booth JK, Wong DCJ, Rensing KH, Laflamme-Yonkman J, Castellarin SD, Bohlmann J, Page JE, Samuels AL (2020) Cannabis glandular trichomes alter morphology and metabolite content during flower maturation. The Plant Journal 101: 37–56

Mansouri H, Asrar Z, Szopa J (2009) Effects of ABA on primary terpenoids and Δ9-tetrahydrocannabinol in Cannabis sativa L. at flowering stage. Plant Growth Regul 58: 269–277

McClelland VF (1968) Superphosphate on wheat: The cumulative effect of repeated applications on yield response. Aust J Agric Res 19: 1–8

Mediavilla V, Jonquera M, Schmid-Slembrouck I, Soldati A (1998) Decimal code for growth stages of hemp (Cannabis sativa L.). SOURCE: JOURNAL OF THE INTERNATIONAL HEMP ASSOCIATION 5: 68–74

Medina S, Vicente R, Nieto-Taladriz MT, Aparicio N, Chairi F, Vergara-Diaz O, Araus JL (2019) The Plant-Transpiration Response to Vapor Pressure Deficit (VPD) in Durum Wheat Is Associated With Differential Yield Performance and Specific Expression of Genes Involved in Primary Metabolism and Water Transport. Front Plant Sci. doi: 10.3389/FPLS.2018.01994

Milay L, Berman P, Shapira A, Guberman O, Meiri D (2020) Metabolic Profiling of Cannabis Secondary Metabolites for Evaluation of Optimal Postharvest Storage Conditions. Front Plant Sci. doi: 10.3389/FPLS.2020.583605/FULL

Morgan W, Singh J, Kesheimer K, Davis J, Sanz-Saez A (2024) Severe drought significantly reduces floral hemp (Cannabis sativa L.) yield and cannabinoid content but moderate drought does not. Environ Exp Bot 219: 105649

Moshelion M (2020) The dichotomy of yield and drought resistance. EMBO Rep 21: e51598

Moshelion M, Dietz KJ, Dodd IC, Muller B, Lunn JE (2024) Guidelines for designing and interpreting drought experiments in controlled conditions. J Exp Bot 75: 4671–4679

Namdar D, Charuvi D, Ajjampura V, Mazuz M, Ion A, Kamara I, Koltai H (2019) LED lighting affects the composition and biological activity of Cannabis sativa secondary metabolites. Ind Crops Prod 132: 177–185

Nelson SH, Hwang KE (2008) Water usage by potato plants at different stages of growth. undefined 52: 331–339

O’Brien C, Arathi HS (2019) Bee diversity and abundance on flowers of industrial hemp (Cannabis sativa L.). Biomass Bioenergy 122: 331–335

Paduch R, Kandefer-Szerszeń M, Trytek M, Fiedurek J (2007) Terpenes: Substances useful in human healthcare. Arch Immunol Ther Exp (Warsz) 55: 315–327

Parsons JL, Martin SL, James T, Golenia G, Boudko EA, Hepworth SR (2019) Polyploidization for the genetic improvement of cannabis sativa. Front Plant Sci 10: 476

Peltonen-Sainio P, Kontturi M, Peltonen J (2006) Phosphorus seed coating enhancement on early growth and yield components in oat.

Plumb J, Demirel S, Sackett JL, Russo EB, Wilson-Poe AR (2022) The Nose Knows: Aroma, but Not THC Mediates the Subjective Effects of Smoked and Vaporized Cannabis Flower. Psychoactives 2022, Vol 1, Pages 70-86 1: 70–86

Rayner JN (1969) Climate and Agriculture: An Ecological Survey, by Jen-hu Chang. Geogr Anal 1: 216–217

Russo EB (2011) Taming THC: potential cannabis synergy and phytocannabinoid-terpenoid entourage effects. Br J Pharmacol 163: 1344–1364

Sadras VO, Calviño PA (2001) Quantification of Grain Yield Response to Soil Depth in Soybean, Maize, Sunflower, and Wheat. Agron J 93: 577–583

Schuurink R, Tissier A (2020) Glandular trichomes: micro-organs with model status? New Phytologist 225: 2251–2266

Selmar D, Kleinwächter M (2013) Stress Enhances the Synthesis of Secondary Plant Products: The Impact of Stress-Related Over-Reduction on the Accumulation of Natural Products. Plant Cell Physiol 54: 817–826

Shapira A, Berman P, Futoran K, Guberman O, Meiri D (2019) Tandem Mass Spectrometric Quantification of 93 Terpenoids in Cannabis Using Static Headspace Injections. Anal Chem 91: 11425–11432

Singh B, Sharma RA (2015) Plant terpenes: defense responses, phylogenetic analysis, regulation and clinical applications. 3 Biotech 5: 129–151

Small E, Antle T (2008) A Preliminary Study of Pollen Dispersal in Cannabis sativa in Relation to Wind Direction. http://dx.doi.org/101300/J237v08n02_03 8: 37–50

Small E, Beckstead HD (1973) Cannabinoid Phenotypes in Cannabis sativa. Nature 1973 245:5421 245: 147–148

Sommano SR, Chittasupho C, Ruksiriwanich W, Jantrawut P (2020) The Cannabis Terpenes. Molecules 2020, Vol 25, Page 5792 25: 5792

Taiz L, Zeiger E, Møller IM, Murphy A (2015) Plant physiology and development. Plant physiology and development.

Thoma F, Somborn-Schulz A, Schlehuber D, Keuter V, Deerberg G (2020) Effects of Light on Secondary Metabolites in Selected Leafy Greens: A Review. Front Plant Sci 11: 497

Toffolatti SL, Maddalena G, Passera A, Casati P, Bianco PA, Quaglino F (2021) Role of terpenes in plant defense to biotic stress. Biocontrol Agents and Secondary Metabolites 401–417

Walsh Z, Callaway R, Belle-Isle L, Capler R, Kay R, Lucas P, Holtzman S (2013) Cannabis for therapeutic purposes: Patient characteristics, access, and reasons for use. International Journal of Drug Policy 24: 511–516

Ware MA, Adams H, Guy GW (2005) The medicinal use of cannabis in the UK: results of a nationwide survey. Int J Clin Pract 59: 291–295

Yaaran A, Negin B, Moshelion M (2019) Role of guard-cell ABA in determining steady-state stomatal aperture and prompt vapor-pressure-deficit response. Plant Science 281: 31–40

Zhai Z, Keereetaweep J, Liu H, Xu C, Shanklin J (2021) The Role of Sugar Signaling in Regulating Plant Fatty Acid Synthesis. Front Plant Sci 12: 374

Zou S, Kumar U (2018) Cannabinoid Receptors and the Endocannabinoid System: Signaling and Function in the Central Nervous System. Int J Mol Sci. doi: 10.3390/IJMS19030833

