## Supplementary figures and tables for "Effects of Drought on Inflorescence Yield, and Secondary Metabolites in *Cannabis sativa* L."

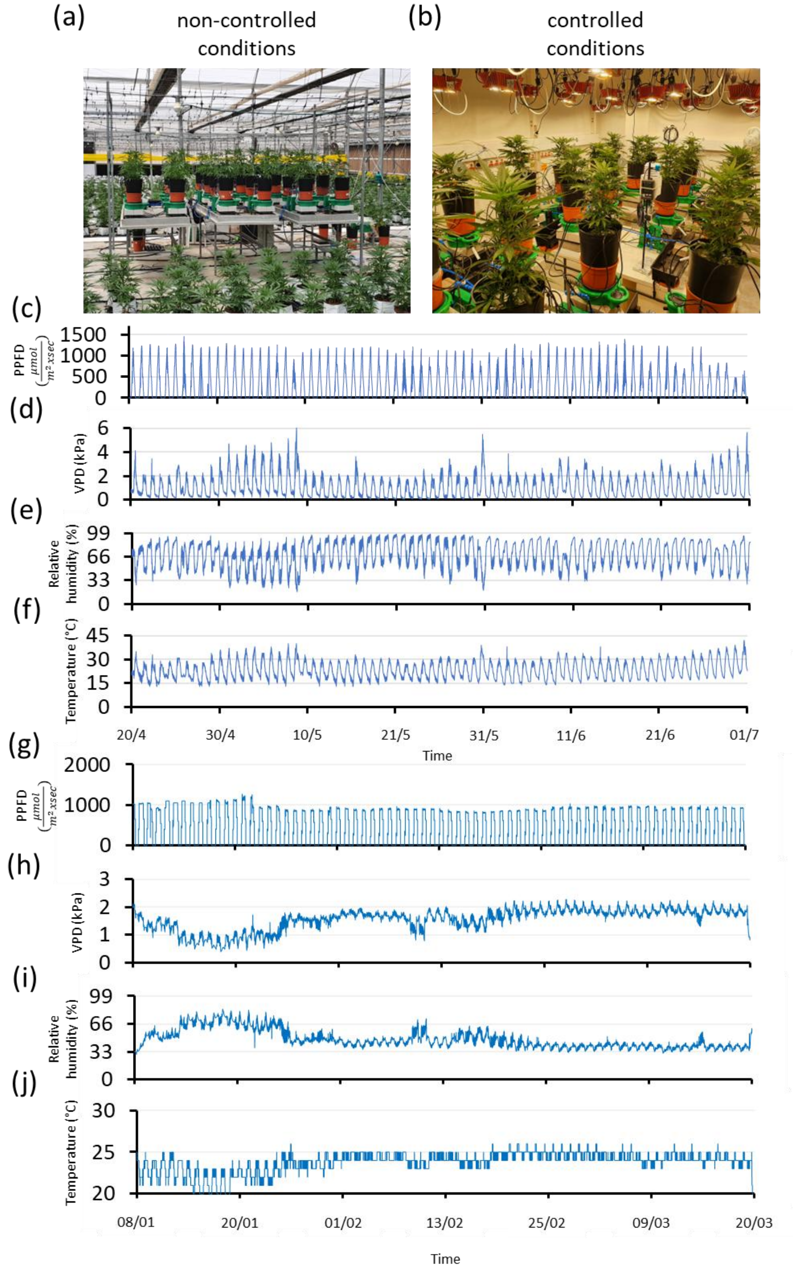


**Fig. S1.** Experimental sites and atmospheric conditions. Representative images of experimental set-ups are shown: (a) non-controlled atmospheric conditions (greenhouse), (b) controlled atmospheric conditions (growth chamber). Panels c–j show continuous meteorological data recorded throughout the experiments under the two examined atmospheric conditions: non-controlled Experiment 4: (c) PPFD radiation, (d) VPD, (e) relative humidity, and (f) temperature, and controlled environment Experiment 2: (g) PPFD radiation, (h) VPD, (i) relative humidity, and (j) temperature. There were minor fluctuations in the data from the controlled chamber, due to variations in day length and manual lamp adjustments to maintain a consistent canopy light intensity (adjustments account for the impact of changes in lamp distance as the plants grew). To ensure uniformity in chamber conditions, annual values were calculated, as described in detail in the Materials and Methods.

| **Mineral** | **Concentration (mg/L; ppm)** |
|---|---|
| N | 130 |
| P | 30 |
| K | 220 |
| Ca | 100 |
| Mg | 60 |
| Fe | 1.4 |
| B | 0.24 |
| Mo | 0.027 |
| Mn | 0.54 |
| Cu | 0.057 |

**Table S1.** Mineral content of the fertilizer solution. Pure mineral concentrations (mg/L; ppm) of the fertigation solution applied during the short-day period, measured using inductively coupled plasma optical emission spectroscopy (ICP-OES). The nitrogen concentration was calculated.

| Compound | CAS number | Company | Country |
|---|---|---|---|
| THCA | 23978-85-0 | USP Ltd | United States |
| THC | 34675-49-5 | Cerilliant Ltd | United States |
| CBGA | 25555-57-1 | Cerilliant Ltd | United States |
| CBDA | 1244-58-2 | Cerilliant Ltd | United States |
| CBD | 25654-31-3 | Cerilliant Ltd | United States |
| THCVA | 31262-37-0 | Cerilliant Ltd | United States |

**Table. S2** Phytocannabinoids HPLC analysis standard solutions**.**

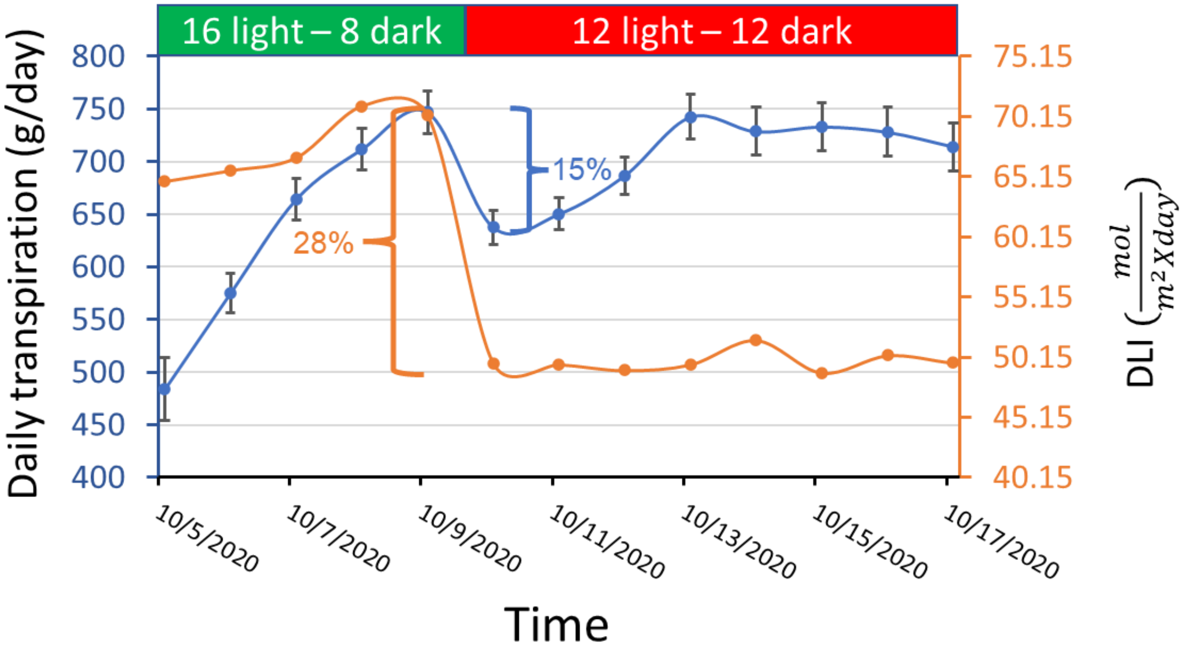


**Fig. S2.** The immediate physiological response of 'Odem' plants to changes in the daily light integral (DLI). Mean ± SE values of the daily transpiration (blue) and DLI (orange) of 'Odem' plants grown under well-irrigated conditions (Experiment 1, Table 1). The upper bar indicates the duration of artificial light. A decrease in DLI during the photoperiod shift was accompanied by an initial reduction in daily transpiration, followed by recovery (see Results section for details).

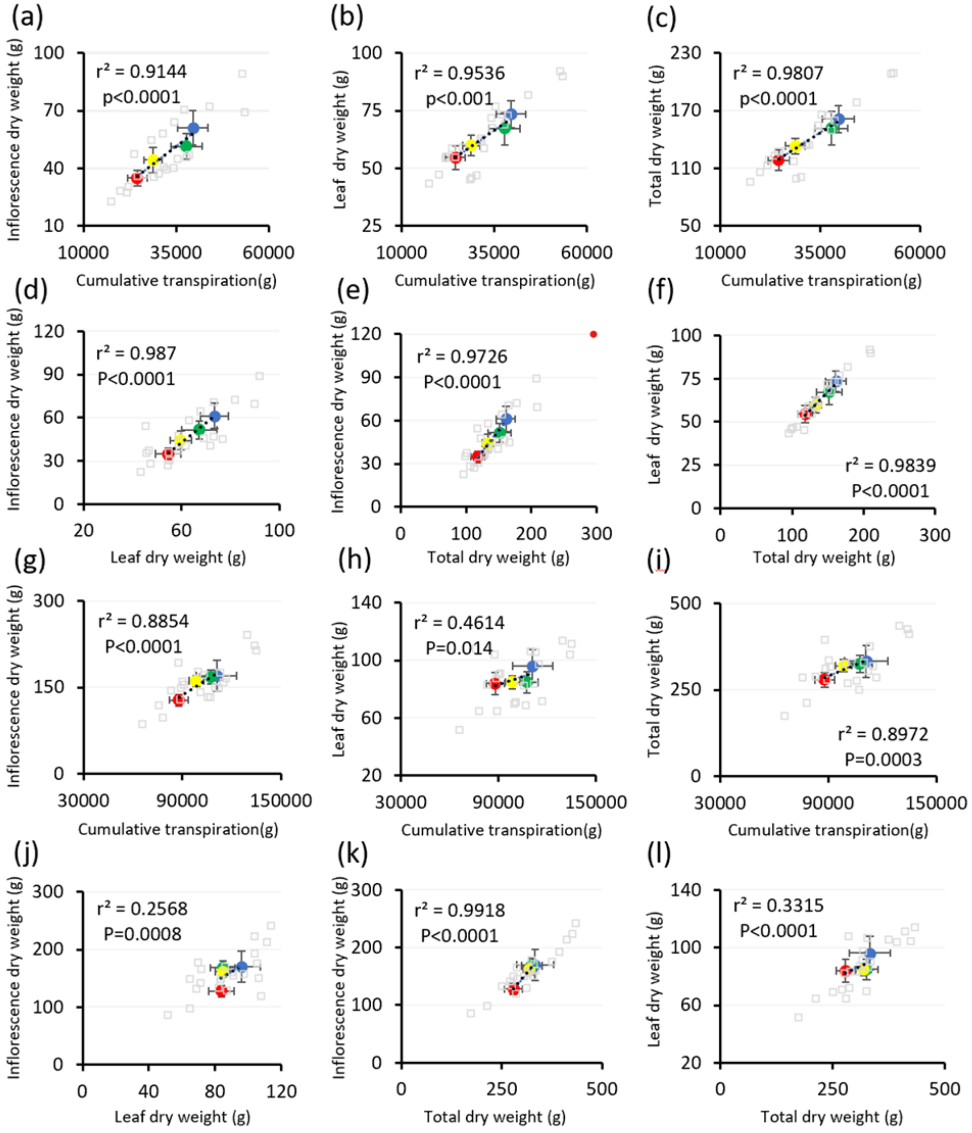


**Fig. S3.** Effects of the drought treatments on the relationship between oven-dry weight biomass and cumulative transpiration under non-controlled atmospheric conditions. Correlation analysis of two lines: Line 187 (Experiment 5; a–f) and 'Odem' (Experiment 4; g–l). (a, g) Correlation between inflorescence dry weight and cumulative transpiration. (b, h) Correlation between leaf dry weight and cumulative transpiration. (c, i) Correlation between total dry weight and cumulative transpiration. (d, j) Correlation between inflorescence dry weight and leaf dry weight. (e, k) Correlation between inflorescence dry weight and total dry weight. (f, l) Correlation between leaf dry weight and total dry weight. Different colors represent the different treatments: blue – control, green – mild drought, yellow – moderate drought, and red – severe drought. Gray points represent all correlated plant data points. R2 represents the mean Pearson correlation coefficient square value. The correlation P-value refers to all plant data points subjected to a multivariate coefficient probability test.

| Experiment | Cultivar | Treatment | Cumulative transpiration (g) | Inflorescence dry weight (g) | Stem dry weight (g) | Leaf dry weight (g) | Total dry weight (g) | Agronomic WUE (g/g) |
|---|---|---|---|---|---|---|---|---|
| 1 | Odem | Control | 120572.5±6023.5 (A) | 177.6±8.6 (A) | 54.8±1.5 (A) | 130.4±3.3 (A) | 362.8±10.1 (A) | 0.0030±0.00014 (B) |
| | | Low drought | 96634.5±5246.1 (B) | 155.4±10.8 (AB) | 54.8±2.0 (A) | 115.8±6.2 (AB) | 325.9±17.4 (AB) | 0.0034±0.00031 (AB) |
| | | Medium drought | 77281.5±2700.3 (C) | 128.8±9.8 (BC) | 52.3±2.1 (A) | 99.7±5.6 (B) | 280.7±14.8 (BC) | 0.0037±0.00023 (AB) |
| | | Severe drought | 62792.8±1735.3 (C) | 105.7±7.4 (C) | 51.6±1.5 (A) | 98.0±2.9 (B) | 255.3±8.6 (C) | 0.0041±0.00017 (A) |
| 2 | Odem | Control | 95189.8±6606.2 (A) | 104.7±8.1 (A) | - | - | - | - |
| | | Low drought | 76952.8±3159.4 (B) | 94.6±3.5 (A) | - | - | - | - |
| | | Medium drought | 71321.2±3638.2 (B) | 83.5±6.8 (AB) | - | - | - | - |
| | | Severe drought | 64803.4±3538.1 (B) | 64.7±5.0 (B) | - | - | - | - |
| 3 | MVA | Control | 109559.7±12521.9 (A) | 143.2±19.0 (A) | 46.9±6.6 (A) | 119.7±18.4 (A) | 309.8±38.5 (A) | 0.0028±0.00016 (A) |
| | | Low drought | 81066.5±12386.1 (A) | 97.8±10.9 (AB) | 36.8±6.4 (A) | 100.0±11.9 (A) | 234.6±27.6 (A) | 0.0030±0.00016 (A) |
| | | Medium drought | 71500.0±13299.8 (A) | 87.6±15.6 (AB) | 40.3±12.3 (A) | 95.4±19.0 (A) | 223.3±46.6 (A) | 0.0031±0.00024 (A) |
| | | Severe drought | 63578.8±10666.7 (A) | 72.9±9.7 (AB) | 41.9±10.4 (A) | 96.8±18.1 (A) | 211.5±37.4 (A) | 0.0033±0.00025 (A) |
| 4 | Odem | Control | 111016.8±12207.3 (A) | 170.1±27.0 (A) | 66.6±8.7 (A) | 96.1±11.5 (A) | 332.8±45.9 (A) | 0.0030±0.00012 (A) |
| | | Low drought | 107491.8±6969.0 (A) | 168.4±12.0 (A) | 72.3±8.0 (A) | 84.8±7.3 (A) | 325.5±25.3 (A) | 0.0030±0.00016 (A) |
| | | Medium drought | 98552.7±4511.2 (A) | 161.5±9.2 (A) | 73.4±5.5 (A) | 84.8±4.5 (A) | 319.7±17.0 (A) | 0.0033±0.00027 (A) |
| | | Severe drought | 87915.0±5535.5 (A) | 128.0±10.1 (A) | 67.3±9.5 (A) | 83.9±7.8 (A) | 279.2±21.7 (A) | 0.0032±0.00029 (A) |
| 5 | 187 | Control | 39524.9±4065.1 (A) | 61.1±8.9 (A) | 26.5±3.9 (A) | 73.5±5.8 (A) * | 161.1±14.2 (A) * | 0.0041±0.00008 (AB) |
| | | Low drought | 37803.2±4146.0 (A) | 51.4±6.5 (AB) | 33.0±6.0 (A) | 67.2±7.2 (A) | 151.6±17.7 (A) | 0.0040±0.00019 (B) |
| | | Medium drought | 28775.6±2396.9 (AB) | 44.2±6.4 (AB) | 29.3±3.4 (A) | 59.7±4.4 (A) | 133.3±8.1 (A) | 0.0047±0.00018 (AB) |
| | | Severe drought | 24560.7±2664.8 (B) | 34.9±4.0 (B) | 28.7±4.2 (A) | 54.6±5.0 (A) * | 118.1±10.5 (A) * | 0.0049±0.00032 (A) |

**Table S3.** Effects of the drought treatments on key physiological traits. Cumulative transpiration, inflorescence dry weight, stem dry weight, leaf dry weight, total dry weight, and agronomic WUE of plants subjected to 4 different drought treatments in 5 different experiments. (The experiments are described in Table 1.) Values are means  SE. Different letters represent significant differences according to one-way ANOVA (between irrigation treatments for a given trait) and Tukey's HSD test ($P < 0.05$, $5 \geq N \geq 6$). An asterisk represents a significant difference between irrigation treatments within each trait, according to Student's t-test ($P < 0.05$).

| Phytocannabinoid | Control | Mild drought | Moderate drought | Severe drought |
|---|---|---|---|---|
| THCA | 20.5743±1.65082 (A) | 20.5689±0.40319 (A) | 18.5980±0.94407 (AB) | 15.2716±1.26778 (B) |
| THC | 0.2561±0.03594 (A) | 0.3046±0.01579 (A) | 0.2759±0.01886 (A) | 0.2553±0.03184 (A) |
| CBDA | 0.0793±0.00640 (A) | 0.0798±0.00244 (A) | 0.0743±0.00361 (A) | 0.0643±0.00473 (A) |
| CBGA | 0.9117±0.10468 (A) | 0.6731±0.01701 (B) | 0.5054±0.03375 (BC) | 0.3532±0.05337 (B) |
| CBG | 0.0416±0.00371 (A) | 0.0437±0.00296 (A) | 0.0350±0.00412 (AB) | 0.0243±0.00248 (B) |
| CBDVA | 0.0338±0.00195 (A) | 0.0356±0.00140 (A) | 0.0350±0.00159 (A) | 0.0295±0.00170 (A) |
| CBNA | 0.0585±0.00422 (A) | 0.0669±0.00224 (A) | 0.0683±0.00426 (A) | 0.0620±0.00398 (A) |
| CBN | 0.0020±0.00027 (B) | 0.0030±0.00029 (AB) | 0.0035±0.00032 (A) | 0.0035±0.00027 (A) |
| CBCA | 0.3948±0.04136 (A) | 0.3893±0.02056 (A) | 0.3960±0.02146 (A) | 0.3583±0.01521 (A) |
| CBC | 0.0138±0.00253 (A) | 0.0142±0.00106 (A) | 0.0116±0.00113 (A) | 0.0104±0.00193 (A) |
| CBGA | 0.7359±0.04484 (A) | 0.5725±0.04329 (B) | 0.3926±0.02501 (C) | 0.2835±0.04509 (C) |
| CBG | 0.0701±0.00422 (A) | 0.0680±0.00519 (A) | 0.0490±0.00460 (B) | 0.0375±0.00416 (B) |
| CBGA-C4 | 0.0004±0.00004 (A) | 0.0003±0.00002 (A) | 0.0002±0.00001 (B) | 0.0002±0.00004 (B) |
| CBGVA | 0.0004±0.00005 (A) | 0.0003±0.00002 (B) | 0.0002±0.00003 (BC) | 0.0001±0.00004 (C) |
| Sesqui-CBGA | 0.0008±0.00007 (A) | 0.0008±0.00008 (A) | 0.0009±0.00007 (A) | 0.0010±0.00011 (A) |
| Sesqui-CBG | 0.0043±0.00028 (A) | 0.0045±0.00025 (A) | 0.0038±0.00039 (A) | 0.0038±0.00040 (A) |
| THC | 0.1634±0.01792 (A) | 0.1953±0.00810 (A) | 0.1784±0.01222 (A) | 0.1611±0.02329 (A) |
| THCA-C4 | 0.0321±0.00258 (A) | 0.0314±0.00117 (A) | 0.0279±0.00147 (AB) | 0.0221±0.00159 (B) |
| THCVA | 0.1275±0.01251 (A) | 0.1200±0.01077 (A) | 0.0940±0.00573 (AB) | 0.0802±0.00757 (B) |
| THCV | 0.0013±0.00019 (A) | 0.0014±0.00009 (A) | 0.0013±0.00008 (A) | 0.0012±0.00011 (A) |
| THCOA | 0.0102±0.00072 (A) | 0.0105±0.00102 (A) | 0.0095±0.00036 (A) | 0.0097±0.00107 (A) |
| THCMA | 0.0148±0.00158 (A) | 0.0164±0.00182 (A) | 0.0126±0.00171 (A) | 0.0136±0.00095 (A) |
| CBDA | 0.0409±0.00211 (A) | 0.0410±0.00220 (A) | 0.0357±0.00172 (AB) | 0.0309±0.00338 (B) |
| CBCA | 0.3179±0.03314 (A) | 0.3205±0.03386 (A) | 0.3105±0.01944 (A) | 0.2770±0.01444 (A) |
| CBC | 0.0047±0.00103 (A) | 0.0049±0.00019 (A) | 0.0054±0.00060 (A) | 0.0039±0.00043 (A) |
| CBCVA | 0.0020±0.00019 (A) | 0.0019±0.00016 (A) | 0.0017±0.00013 (A) | 0.0017±0.00016 (A) |
| CBNA | 0.0555±0.00267 (A) | 0.0616±0.00290 (A) | 0.0597±0.00351 (A) | 0.0568±0.00337 (A) |
| CBN | 0.0014±0.00012 (A) | 0.0018±0.00027 (A) | 0.0017±0.00019 (A) | 0.0020±0.00023 (A) |
| CBNVA | 0.0003±0.00002 (A) | 0.0003±0.00002 (A) | 0.0003±0.00002 (A) | 0.0003±0.00002 (A) |
| CBE | 0.0014±0.00029 (A) | 0.0012±0.00006 (A) | 0.0011±0.00012 (A) | 0.0008±0.00012 (A) |
| CBTA-1 | 0.0032±0.00022 (B) | 0.0043±0.00028 (AB) | 0.0042±0.00037 (AB) | 0.0046±0.00035 (A) |
| CBT-1 | 0.0012±0.00009 (A) | 0.0018±0.00026 (A) | 0.0018±0.00023 (A) | 0.0022±0.00033 (A) |
| CBTA-3 | 0.0033±0.00021 (A) | 0.0036±0.00026 (A) | 0.0033±0.00024 (A) | 0.0032±0.00016 (A) |
| CBT-3 | 0.0009±0.00008 (A) | 0.0012±0.00013 (A) | 0.0010±0.00010 (A) | 0.0011±0.00012 (A) |

**Table. S4** continued on the next page

**Table. S4** continued from the previous page

| Phytocannabinoid | Control | Low drought | Medium drought | Severe drought |
|---|---|---|---|---|
| 329-11b | 0.0002±0.00010 (A) | 0.0004±0.00017 (A) | 0.0007±0.00022 (A) | 0.0009±0.00035 (A) |
| 329-11d | 0.0009±0.00006 (A) | 0.0011±0.00011 (A) | 0.0010±0.00010 (A) | 0.0010±0.00006 (A) |
| 373-12b | 0.0065±0.00137 (A) | 0.0084±0.00241 (A) | 0.0126±0.00372 (A) | 0.0152±0.00558 (A) |
| 373-12c | 0.0040±0.00035 (A) | 0.0047±0.00072 (A) | 0.0044±0.00069 (A) | 0.0040±0.00069 (A) |
| 373-12d | 0.0024±0.00015 (A) | 0.0023±0.00013 (A) | 0.0021±0.00013 (AB) | 0.0017±0.00011 (B) |
| 371-14a | 0.0003±0.00002 (A) | 0.0003±0.00002 (A) | 0.0003±0.00002 (A) | 0.0003±0.00002 (A) |
| 371-14b | 0.0013±0.00008 (AB) | 0.0014±0.00012 (A) | 0.0013±0.00012 (AB) | 0.0010±0.00006 (B) |
| 417-15a | 0.0037±0.00045 (AB) | 0.0051±0.00062 (A) | 0.0049±0.00071 (AB) | 0.0027±0.00025 (B) |
| 357-16a | 0.0110±0.00111 (A) | 0.0120±0.00125 (A) | 0.0093±0.00116 (A) | 0.0097±0.00095 (A) |
| 313-16b | 0.0004±0.00011 (A) | 0.0005±0.00004 (A) | 0.0004±0.00008 (A) | 0.0002±0.00013 (A) |
| 361-17a | 0.0008±0.00006 (A) | 0.0012±0.00015 (A) | 0.0011±0.00014 (A) | 0.0013±0.00017 (A) |
| 331-18b | 0.1256±0.01098 (A) | 0.1248±0.00668 (A) | 0.1054±0.00971 (A) | 0.0932±0.01330 (A) |
| 331-18c | 0.0018±0.00017 (A) | 0.0022±0.00012 (A) | 0.0019±0.00016 (A) | 0.0018±0.00018 (A) |
| 331-18d | 0.1130±0.00629 (A) | 0.1058±0.00579 (AB) | 0.0825±0.00382 (BC) | 0.0640±0.01065 (C) |
| 375-19a | 0.0015±0.00012 (A) | 0.0015±0.00009 (A) | 0.0013±0.00006 (A) | 0.0011±0.00011 (A) |
| 375-19b | 0.0004±0.00002 (A) | 0.0003±0.00003 (B) | 0.0002±0.00002 (BC) | 0.0002±0.00002 (C) |
| 375-19c | 0.0006±0.00005 (A) | 0.0005±0.00004 (AB) | 0.0004±0.00002 (BC) | 0.0003±0.00003 (C) |
| Total THC | 18.2998±1.48035 (A) | 18.3435±0.36085 (A) | 16.5863±0.84447 (AB) | 13.6485±1.13983 (B) |
| Total CBD | 0.0695±0.00561 (A) | 0.0700±0.00214 (A) | 0.0651±0.00317 (A) | 0.0564±0.00415 (A) |
| Total CBG | 0.8411±0.09515 (A) | 0.6340±0.01618 (AB) | 0.4782±0.03213 (BC) | 0.3341±0.04851 (C) |

**Table S4.** Effects of the drought treatments on the phytocannabinoid content of primary inflorescences of 'Odem' plants (Experiment 1). Phytocannabinoid concentrations (%, w/w) in the primary inflorescences of 'Odem' plants subjected to 4 different irrigation treatments, as measured at the end of Experiment 1. Samples were taken from the primary inflorescences. Values are means ± SE. Different letters represent significant differences between irrigation treatments, according to one-way ANOVA and Tukey's HSD test ($P < 0.05$, $5 \leq N \leq 6$).

| Separation method | Phytocannabinoid | Control | Mild drought | Moderate drought | Severe drought |
|---|---|---|---|---|---|
| UHPLC | THCA | 17.71285±1.66946 (A) | 18.81570±0.59626 (AB) | 17.12302±0.50439 (AB) | 13.71875±0.10065 (B) |
| UHPLC | THC | 0.18798±0.01199 (B) | 0.24173±0.01068 (A) | 0.23207±0.01014 (AB) | 0.21888±0.02149 (AB) |
| UHPLC | CBDA | 0.06758±0.00502 (AB) | 0.07161±0.00308 (A) | 0.06600±0.00146 (AB) | 0.05510±0.00033 (B) |
| UHPLC | CBGA | 0.74139±0.06581 (A) | 0.57824±0.02670 (AB) | 0.42036±0.03585 (BC) | 0.26738±0.02796 (C) |
| UHPLC | CBG | 0.03745±0.00567 (AB) | 0.04540±0.00378 (A) | 0.03735±0.00291 (AB) | 0.02494±0.00266 (B) |
| UHPLC | CBDVA | 0.03529±0.00477 (A) | 0.03970±0.00211 (A) | 0.03976±0.00189 (A) | 0.03446±0.00184 (A) |
| UHPLC | CBNA | 0.05867±0.00903 (A) | 0.07123±0.00288 (A) | 0.07413±0.00317 (A) | 0.07111±0.00670 (A) |
| UHPLC | CBN | 0.00175±0.00021 (B) | 0.00268±0.00024 (AB) | 0.00326±0.00025 (A) | 0.00353±0.00050 (A) |
| UHPLC | CBCA | 0.34869±0.03087 (A) | 0.32327±0.02235 (A) | 0.31413±0.01790 (A) | 0.27130±0.01720 (A) |
| UHPLC | CBC | 0.01040±0.00048 (A) | 0.00984±0.00054 (AB) | 0.00888±0.00071 (AB) | 0.00732±0.00071 (B) |
| LCMS | CBGA | 0.48355±0.08757 (A) | 0.37363±0.03469 (A) | 0.31482±0.02580 (A) | 0.30716±0.12739 (A) |
| LCMS | CBG | 0.04602±0.00933 (A) | 0.05328±0.00380 (A) | 0.04504±0.00214 (A) | 0.04058±0.01458 (A) |
| LCMS | CBGA-C4 | 0.00030±0.00007 (A) | 0.00024±0.00005 (A) | 0.00020±0.00001 (A) | 0.00019±0.00012 (A) |
| LCMS | CBGVA | 0.00024±0.00007 (A) | 0.00017±0.00004 (AB) | 0.00003±0.00003 (B) | 0.00012±0.00012 (AB) |
| LCMS | Sesqui-CBGA | 0.00057±0.00008 (A) | 0.00054±0.00004 (A) | 0.00058±0.00003 (A) | 0.00065±0.00005 (A) |
| LCMS | Sesqui-CBG | 0.00292±0.00053 (A) | 0.00334±0.00015 (A) | 0.00314±0.00019 (A) | 0.00328±0.00058 (A) |
| LCMS | THC | 0.13290±0.00965 (A) | 0.15486±0.00606 (A) | 0.15891±0.00568 (A) | 0.15592±0.00276 (A) |
| LCMS | THCA-C4 | 0.02607±0.00312 (A) | 0.02850±0.00129 (A) | 0.02632±0.00058 (A) | 0.02474±0.00384 (A) |
| LCMS | THCVA | 0.08513±0.01136 (A) | 0.08012±0.00460 (A) | 0.07149±0.00281 (A) | 0.07372±0.01974 (A) |
| LCMS | THCV | 0.00099±0.00005 (A) | 0.00115±0.00007 (A) | 0.00105±0.00004 (A) | 0.00108±0.00006 (A) |
| LCMS | THCOA | 0.00826±0.00091 (A) | 0.00802±0.00089 (A) | 0.00746±0.00029 (A) | 0.00797±0.00054 (A) |
| LCMS | THCMA | 0.00929±0.00217 (A) | 0.00973±0.00065 (A) | 0.00891±0.00047 (A) | 0.00913±0.00058 (A) |
| LCMS | CBDA | 0.02946±0.00432 (A) | 0.03086±0.00195 (A) | 0.02915±0.00109 (A) | 0.02717±0.00388 (A) |
| LCMS | CBCA | 0.24485±0.02624 (A) | 0.23142±0.01517 (A) | 0.24075±0.02272 (A) | 0.22550±0.04110 (A) |
| LCMS | CBC | 0.00411±0.00045 (A) | 0.00459±0.00015 (A) | 0.00454±0.00021 (A) | 0.00465±0.00058 (A) |
| LCMS | CBCA-C4 | 0.00025±0.00005 (A) | 0.00030±0.00003 (A) | 0.00018±0.00006 (A) | 0.00021±0.00013 (A) |
| LCMS | CBCVA | 0.00149±0.00018 (A) | 0.00141±0.00010 (A) | 0.00125±0.00008 (A) | 0.00135±0.00037 (A) |
| LCMS | CBNA | 0.05129±0.00810 (A) | 0.05781±0.00267 (A) | 0.05775±0.00144 (A) | 0.06045±0.00294 (A) |
| LCMS | CBN | 0.00180±0.00072 (A) | 0.00166±0.00018 (A) | 0.00170±0.00019 (A) | 0.00205±0.00042 (A) |
| LCMS | CBNA-C4 | 0.00004±0.00002 (A) | 0.00007±0.00001 (A) | 0.00004±0.00002 (A) | 0.00008±0.00000 (A) |
| LCMS | CBNVA | 0.00021±0.00006 (A) | 0.00025±0.00002 (A) | 0.00021±0.00001 (A) | 0.00026±0.00004 (A) |
| LCMS | CBE | 0.00093±0.00015 (A) | 0.00111±0.00006 (A) | 0.00099±0.00004 (A) | 0.00087±0.00028 (A) |
| LCMS | CBTA-1 | 0.00304±0.00059 (A) | 0.00336±0.00017 (A) | 0.00344±0.00011 (A) | 0.00421±0.00050 (A) |
| LCMS | CBT-1 | 0.00079±0.00051 (A) | 0.00130±0.00029 (A) | 0.00151±0.00017 (A) | 0.00211±0.00058 (A) |

**Table. S5** continued on the next page

**Table. S5** continued from the previous page

| Separation method | Phytocannabinoid | Control | Mild drought | Moderate drought | Severe drought |
|---|---|---|---|---|---|
| LCMS | CBTA-3 | 0.00317±0.00060 (A) | 0.00329±0.00022 (A) | 0.00323±0.00012 (A) | 0.00317±0.00035 (A) |
| LCMS | CBT-3 | 0.00078±0.00027 (A) | 0.00078±0.00007 (A) | 0.00077±0.00009 (A) | 0.00092±0.00016 (A) |
| LCMS | 329-11b | 0.00025±0.00011 (A) | 0.00067±0.00022 (A) | 0.00051±0.00025 (A) | 0.00047±0.00008 (A) |
| LCMS | 329-11d | 0.00066±0.00012 (A) | 0.00089±0.00013 (A) | 0.00073±0.00006 (A) | 0.00086±0.00007 (A) |
| LCMS | 373-12b | 0.00646±0.00184 (A) | 0.01181±0.00331 (A) | 0.01091±0.00409 (A) | 0.00732±0.00189 (A) |
| LCMS | 373-12c | 0.00427±0.00100 (A) | 0.00549±0.00070 (A) | 0.00484±0.00096 (A) | 0.00450±0.00140 (A) |
| LCMS | 373-12d | 0.00236±0.00038 (A) | 0.00245±0.00014 (A) | 0.00237±0.00011 (A) | 0.00260±0.00047 (A) |
| LCMS | 371-14a | 0.00030±0.00008 (A) | 0.00036±0.00003 (A) | 0.00033±0.00002 (A) | 0.00036±0.00004 (A) |
| LCMS | 371-14b | 0.00125±0.00021 (A) | 0.00130±0.00006 (A) | 0.00121±0.00008 (A) | 0.00130±0.00027 (A) |
| LCMS | 417-15a | 0.00334±0.00034 (A) | 0.00370±0.00016 (A) | 0.00372±0.00015 (A) | 0.00374±0.00064 (A) |
| LCMS | 357-16a | 0.00659±0.00147 (A) | 0.00561±0.00051 (A) | 0.00550±0.00027 (A) | 0.00546±0.00052 (A) |
| LCMS | 313-16b | 0.00018±0.00011 (A) | 0.00032±0.00010 (A) | 0.00029±0.00009 (A) | 0.00027±0.00014 (A) |
| LCMS | 361-17a | 0.00182±0.00127 (A) | 0.00091±0.00010 (A) | 0.00089±0.00010 (A) | 0.00122±0.00030 (A) |
| LCMS | 331-18b | 0.08722±0.01528 (A) | 0.09356±0.00402 (A) | 0.08368±0.00382 (A) | 0.07199±0.02167 (A) |
| LCMS | 331-18c | 0.00139±0.00021 (A) | 0.00177±0.00010 (A) | 0.00157±0.00005 (A) | 0.00147±0.00023 (A) |
| LCMS | 331-18d | 0.07555±0.00936 (A) | 0.07363±0.00631 (A) | 0.06511±0.00391 (A) | 0.05521±0.01414 (A) |
| LCMS | 375-19a | 0.00112±0.00015 (A) | 0.00125±0.00010 (A) | 0.00111±0.00007 (A) | 0.00105±0.00018 (A) |
| LCMS | 375-19b | 0.00039±0.00007 (A) | 0.00034±0.00005 (A) | 0.00028±0.00003 (A) | 0.00032±0.00011 (A) |
| LCMS | 375-19c | 0.00048±0.00007 (A) | 0.00047±0.00005 (A) | 0.00038±0.00003 (A) | 0.00046±0.00013 (A) |
| HPLC | Total THC | 15.722±1.47478 (AB) | 16.743±0.53056 (A) | 15.248±0.44990 (AB) | 12.250±0.09581 (B) |
| HPLC | Total CBD | 0.0592±0.00440 (AB) | 0.06281±0.00270 (A) | 0.05789±0.00128 (AB) | 0.04832±0.00029 (B) |
| HPLC | Total CBG | 0.687±0.06226 (A) | 0.552±0.02622 (AB) | 0.406±0.03278 (BC) | 0.259±0.02532 (C) |
| HPLC | Sum | 19.202±1.77126 (AB) | 20.199±0.61048 (A) | 18.318±0.53629 (AB) | 14.672±0.12085 (B) |

**Table S5.** Effects of the drought treatments on the phytocannabinoid content of secondary inflorescences of 'Odem' plants (Experiment 1). Phytocannabinoid concentrations (%, w/w) in the secondary inflorescences of 'Odem' plants subjected to 4 different irrigation treatments, as measured at the end of Experiment 1. Samples were taken from the secondary inflorescences. Values are means ± SE. Different letters represent significant differences between irrigation treatments according to one-way ANOVA and Tukey's HSD test ($P < 0.05$, $5 \leq N \leq 6$).

| Separation method | Phytocannabinoid | Control | Mild drought | Moderate drought | Severe drought |
|---|---|---|---|---|---|
| HPLC | THCA | 17.3765±1.00610 (A) | 18.5545±0.82930 (A) | 16.0114±0.67689 (A) | 15.4611±1.07330 (A) |
| HPLC | THC | 0.3671±0.03779 (A) | 0.4585±0.02609 (A) | 0.4373±0.03393 (A) | 0.3641±0.02630 (A) |
| HPLC | CBDA | 0.0668±0.00341 (A) | 0.0697±0.00426 (A) | 0.0607±0.00306 (A) | 0.0591±0.00329 (A) |
| HPLC | CBGA | 0.5229±0.06144 (AB) | 0.5869±0.02422 (A) | 0.4708±0.03043 (AB) | 0.4246±0.03414 (B) |
| HPLC | CBG | 0.0312±0.00460 (A) | 0.0353±0.00354 (A) | 0.0286±0.00257 (A) | 0.0266±0.00602 (A) |
| HPLC | CBDVA | 0.0403±0.00294 (A) | 0.0420±0.00198 (A) | 0.0366±0.00099 (A) | 0.0351±0.00197 (A) |
| HPLC | CBNA | 0.0588±0.00651 (A) | 0.0557±0.00429 (A) | 0.0524±0.00493 (A) | 0.0563±0.00608 (A) |
| HPLC | CBN | 0.0036±0.00025 (A) | 0.0040±0.00051 (A) | 0.0041±0.00030 (A) | 0.0040±0.00057 (A) |
| HPLC | CBCA | 0.3273±0.02867 (A) | 0.3235±0.02531 (A) | 0.2801±0.02313 (A) | 0.3220±0.02609 (A) |
| HPLC | CBC | 0.0081±0.00135 (A) | 0.0103±0.00080 (A) | 0.0097±0.00122 (A) | 0.0084±0.00110 (A) |
| LCMS | CBGA-C4 | 0.0003±0.00004 (A) | 0.0004±0.00004 (A) | 0.0003±0.00001 (A) | 0.0003±0.00003 (A) |
| LCMS | CBGVA | 0.0002±0.00005 (A) | 0.0002±0.00005 (A) | 0.0002±0.00003 (A) | 0.0001±0.00005 (A) |
| LCMS | Sesqui-CBGA | 0.0008±0.00004 (A) | 0.0008±0.00004 (A) | 0.0009±0.00005 (A) | 0.0009±0.00006 (A) |
| LCMS | Sesqui-CBG | 0.0066±0.00064 (A) | 0.0068±0.00063 (A) | 0.0063±0.00047 (A) | 0.0064±0.00099 (A) |
| LCMS | THCA-C4 | 0.0369±0.00271 (A) | 0.0374±0.00264 (A) | 0.0374±0.00190 (A) | 0.0352±0.00400 (A) |
| LCMS | THCVA | 0.1225±0.00931 (A) | 0.1186±0.00926 (A) | 0.1077±0.00572 (A) | 0.1069±0.01173 (A) |
| LCMS | THCOA | 0.0117±0.00121 (A) | 0.0109±0.00049 (A) | 0.0106±0.00058 (A) | 0.0106±0.00056 (A) |
| LCMS | THCMA | 0.0096±0.00042 (A) | 0.0099±0.00030 (A) | 0.0108±0.00061 (A) | 0.0095±0.00074 (A) |
| LCMS | CBCA-C4 | 0.0006±0.00008 (A) | 0.0004±0.00004 (A) | 0.0004±0.00006 (A) | 0.0004±0.00010 (A) |
| LCMS | CBCVA | 0.0016±0.00014 (A) | 0.0015±0.00015 (A) | 0.0014±0.00010 (A) | 0.0016±0.00013 (A) |
| LCMS | CBNA-C4 | 0.0001±0.00002 (A) | 0.0001±0.00002 (A) | 0.0001±0.00001 (A) | 0.0001±0.00002 (A) |
| LCMS | CBNVA | 0.0003±0.00003 (A) | 0.0003±0.00002 (A) | 0.0003±0.00002 (A) | 0.0003±0.00003 (A) |
| LCMS | CBTA-1 | 0.0035±0.00038 (A) | 0.0031±0.00013 (A) | 0.0034±0.00021 (A) | 0.0037±0.00041 (A) |
| LCMS | CBT-1 | 0.0014±0.00014 (A) | 0.0015±0.00011 (A) | 0.0018±0.00021 (A) | 0.0017±0.00027 (A) |
| LCMS | CBTA-3 | 0.0039±0.00030 (A) | 0.0039±0.00019 (A) | 0.0037±0.00010 (A) | 0.0035±0.00023 (A) |
| LCMS | CBT-3 | 0.0007±0.00005 (A) | 0.0008±0.00004 (A) | 0.0009±0.00008 (A) | 0.0009±0.00012 (A) |
| LCMS | 329-11d | 0.0015±0.00021 (A) | 0.0016±0.00005 (A) | 0.0014±0.00003 (A) | 0.0015±0.00018 (A) |
| LCMS | 373-12b | 0.0066±0.00281 (A) | 0.0038±0.00137 (A) | 0.0053±0.00130 (A) | 0.0070±0.00190 (A) |
| LCMS | 373-12c | 0.0017±0.00065 (A) | 0.0012±0.00052 (A) | 0.0014±0.00035 (A) | 0.0017±0.00060 (A) |
| LCMS | 373-12d | 0.0042±0.00068 (A) | 0.0042±0.00030 (A) | 0.0035±0.00015 (A) | 0.0036±0.00049 (A) |
| LCMS | 327-13c | 0.0024±0.00020 (A) | 0.0024±0.00023 (A) | 0.0025±0.00015 (A) | 0.0024±0.00011 (A) |
| LCMS | 371-14a | 0.0005±0.00006 (A) | 0.0004±0.00002 (A) | 0.0004±0.00002 (A) | 0.0004±0.00003 (A) |
| LCMS | 371-14b | 0.0014±0.00019 (A) | 0.0013±0.00008 (A) | 0.0012±0.00005 (A) | 0.0011±0.00010 (A) |
| LCMS | 417-15a | 0.0019±0.00016 (A) | 0.0017±0.00014 (A) | 0.0015±0.00006 (A) | 0.0016±0.00017 (A) |

**Table. S6** continued on the next page

**Table. S6** continued from the previous page

| Separation method | Phytocannabinoid | Control | Mild drought | Moderate drought | Severe drought |
|---|---|---|---|---|---|
| LCMS | 357-16a | 0.0090±0.00088 (A) | 0.0092±0.00023 (A) | 0.0084±0.00057 (A) | 0.0076±0.00046 (A) |
| LCMS | 313-16b | 0.0008±0.00007 (A) | 0.0008±0.00004 (A) | 0.0008±0.00006 (A) | 0.0007±0.00003 (A) |
| LCMS | 361-17a | 0.0008±0.00008 (A) | 0.0008±0.00005 (A) | 0.0010±0.00013 (A) | 0.0010±0.00013 (A) |
| LCMS | 331-18b | 0.1024±0.00927 (A) | 0.1000±0.00590 (A) | 0.0927±0.00508 (A) | 0.0920±0.00925 (A) |
| LCMS | 331-18d | 0.0842±0.00748 (A) | 0.0872±0.00310 (A) | 0.0792±0.00412 (A) | 0.0706±0.00396 (A) |
| LCMS | 375-19a | 0.0017±0.00013 (A) | 0.0018±0.00007 (A) | 0.0017±0.00008 (A) | 0.0016±0.00007 (A) |
| LCMS | 375-19b | 0.0006±0.00008 (A) | 0.0006±0.00002 (A) | 0.0005±0.00004 (A) | 0.0004±0.00004 (A) |
| LCMS | 375-19c | 0.0006±0.00008 (A) | 0.0006±0.00003 (A) | 0.0006±0.00003 (A) | 0.0005±0.00005 (A) |
| HPLC | Total THC | 15.6063±0.89953 (A) | 16.7308±0.74204 (A) | 14.4793±0.61525 (A) | 13.9235±0.94052 (A) |
| HPLC | Total CBD | 0.0586±0.00299 (A) | 0.0611±0.00373 (A) | 0.0532±0.00269 (A) | 0.0518±0.00288 (A) |
| HPLC | Total CBG | 0.4897±0.05565 (AB) | 0.5501±0.02302 (A) | 0.4414±0.02632 (AB) | 0.3991±0.03415 (B) |

**Table S6.** Effects of the drought treatments on the phytocannabinoid content of secondary inflorescences of ‘Odem’ plants (Experiment 2). Phytocannabinoid concentrations (%, w/w) in the secondary inflorescences of ‘Odem’ plants subjected to 4 different irrigation treatments, as measured in the middle of physiological Phase III in Experiment 2. Samples were taken from the secondary inflorescences. Values are means ± SE. Different letters represent significant differences between irrigation treatments, according to one-way ANOVA and Tukey’s HSD test ($P < 0.05$, $5 \leq N \leq 6$).

| Separation method | Phytocannabinoid | Control | Mild drought | Moderate drought | Severe drought |
|---|---|---|---|---|---|
| HPLC | THCA | 20.4975±1.22079 (A) | 16.7406±0.59818 (B) | 17.1472±0.56209 (B) | 15.3464±0.64271 (B) |
| HPLC | THC | 1.0590±0.09598 (A) | 1.0138±0.11521 (A) | 1.0160±0.08725 (A) | 1.2744±0.07398 (A) |
| HPLC | CBDA | 0.0822±0.00504 (A) | 0.0668±0.00257 (AB) | 0.0699±0.00233 (B) | 0.0657±0.00324 (B) |
| HPLC | CBGA | 0.8194±0.05629 (A) | 0.6766±0.03612 (AB) | 0.6218±0.02880 (BC) | 0.4953±0.03465 (C) |
| HPLC | CBG | 0.0949±0.00705 (A) | 0.0778±0.00262 (AB) | 0.0770±0.00341 (B) | 0.0590±0.00363 (C) |
| HPLC | CBDVA | 0.0557±0.00280 (A) | 0.0487±0.00269 (AB) | 0.0524±0.00150 (AB) | 0.0437±0.00211 (B) |
| HPLC | CBNA | 0.0994±0.00511 (A) | 0.0911±0.00373 (A) | 0.1010±0.00331 (A) | 0.0981±0.00446 (A) |
| HPLC | CBN | 0.0052±0.00067 (B) | 0.0067±0.00087 (B) | 0.0082±0.00113 (AB) | 0.0118±0.00095 (A) |
| HPLC | CBCA | 0.3756±0.03916 (A) | 0.3567±0.01959 (A) | 0.3651±0.02500 (A) | 0.3870±0.03534 (A) |
| HPLC | CBC | 0.0396±0.00182 (B) | 0.0462±0.00392 (B) | 0.0529±0.00404 (AB) | 0.0672±0.00697 (A) |
| LCMS | CBGA-C4 | 0.0005±0.00003 (A) | 0.0004±0.00002 (AB) | 0.0004±0.00003 (AB) | 0.0003±0.00002 (B) |
| LCMS | CBGVA | 0.0004±0.00004 (A) | 0.0003±0.00001 (AB) | 0.0002±0.00001 (B) | 0.0001±0.00005 (C) |
| LCMS | Sesqui-CBGA | 0.0009±0.00005 (A) | 0.0009±0.00007 (A) | 0.0010±0.00008 (A) | 0.0011±0.00006 (A) |
| LCMS | Sesqui-CBG | 1.1697±0.12476 (A) | 1.1012±0.12465 (A) | 1.1973±0.11862 (A) | 1.3911±0.07518 (A) |
| LCMS | THCA-C4 | 0.0407±0.00304 (A) | 0.0347±0.00085 (AB) | 0.0371±0.00160 (AB) | 0.0327±0.00151 (B) |
| LCMS | THCVA | 0.0063±0.00046 (A) | 0.0064±0.00076 (A) | 0.0063±0.00052 (A) | 0.0084±0.00087 (A) |
| LCMS | THCOA | 0.0129±0.00083 (A) | 0.0106±0.00065 (A) | 0.0117±0.00076 (A) | 0.0122±0.00077 (A) |
| LCMS | THCMA | 0.0099±0.00049 (A) | 0.0091±0.00048 (A) | 0.0106±0.00092 (A) | 0.0093±0.00056 (A) |
| LCMS | CBDA-C4 | 0.0001±0.00001 (A) | 0.0001±0.00002 (A) | 0.0001±0.00002 (A) | 0.0001±0.00001 (A) |
| LCMS | CBDM | 0.0307±0.00210 (A) | 0.0364±0.00343 (AB) | 0.0491±0.00625 (AB) | 0.0548±0.00617 (A) |
| LCMS | CBCA-C4 | 0.0006±0.00007 (A) | 0.0005±0.00007 (A) | 0.0005±0.00004 (A) | 0.0005±0.00007 (A) |
| LCMS | CBCO | 0.0080±0.00057 (B) | 0.0086±0.00092 (B) | 0.0109±0.00130 (AB) | 0.0131±0.00084 (A) |
| LCMS | CBNA-C4 | 0.0002±0.00001 (A) | 0.0002±0.00001 (A) | 0.0002±0.00001 (A) | 0.0002±0.00001 (A) |
| LCMS | CBNVA | 0.0006±0.00005 (A) | 0.0005±0.00002 (A) | 0.0005±0.00009 (A) | 0.0005±0.00003 (A) |
| LCMS | CBTA-1 | 0.0060±0.00042 (A) | 0.0054±0.00021 (A) | 0.0063±0.00027 (A) | 0.0066±0.00044 (A) |
| LCMS | CBT-1 | 0.0055±0.00033 (B) | 0.0058±0.00050 (B) | 0.0070±0.00075 (AB) | 0.0087±0.00074 (A) |
| LCMS | CBTA-3 | 0.0059±0.00034 (A) | 0.0050±0.00025 (BC) | 0.0055±0.00011 (AB) | 0.0045±0.00020 (C) |
| LCMS | CBT-3 | 0.0036±0.00021 (A) | 0.0036±0.00033 (A) | 0.0042±0.00042 (A) | 0.0050±0.00045 (A) |
| LCMS | CBT-2 | 0.0004±0.00002 (A) | 0.0003±0.00003 (A) | 0.0002±0.00007 (A) | 0.0003±0.00005 (A) |
| LCMS | 329-11b | 0.0004±0.00013 (B) | 0.0005±0.00017 (B) | 0.0007±0.00012 (B) | 0.0019±0.00038 (A) |
| LCMS | 329-11d | 0.0023±0.00020 (A) | 0.0022±0.00018 (A) | 0.0024±0.00014 (A) | 0.0023±0.00022 (A) |
| LCMS | 373-12b | 0.0033±0.00067 (A) | 0.0038±0.00124 (A) | 0.0048±0.00106 (A) | 0.0077±0.00212 (A) |
| LCMS | 373-12c | 0.0008±0.00020 (A) | 0.0010±0.00033 (A) | 0.0010±0.00022 (A) | 0.0016±0.00047 (A) |

**Table. S7** continued on the next page

**Table. S7** continued from the previous page

| Separation method | Phytocannabinoid | Control | Mild drought | Moderate drought | Severe drought |
|---|---|---|---|---|---|
| LCMS | 373-12d | 0.0028±0.00017 (A) | 0.0023±0.00014 (AB) | 0.0025±0.00011 (A) | 0.0020±0.00011 (B) |
| LCMS | 327-13c | 0.0028±0.00017 (A) | 0.0024±0.00010 (B) | 0.0025±0.00009 (AB) | 0.0023±0.00009 (B) |
| LCMS | 371-14a | 0.0003±0.00007 (A) | 0.0003±0.00003 (A) | 0.0002±0.00007 (A) | 0.0002±0.00007 (A) |
| LCMS | 371-14b | 0.0013±0.00008 (A) | 0.0013±0.00005 (A) | 0.0013±0.00005 (A) | 0.0010±0.00007 (B) |
| LCMS | 417-15a | 0.0020±0.00034 (A) | 0.0018±0.00019 (A) | 0.0020±0.00021 (A) | 0.0019±0.00022 (A) |
| LCMS | 357-16a | 0.0111±0.00075 (A) | 0.0092±0.00035 (AB) | 0.0094±0.00048 (AB) | 0.0078±0.00037 (B) |
| LCMS | 313-16b | 0.0014±0.00008 (A) | 0.0012±0.00007 (A) | 0.0013±0.00009 (A) | 0.0012±0.00008 (A) |
| LCMS | 361-17a | 0.0025±0.00008 (A) | 0.0026±0.00022 (A) | 0.0034±0.00038 (A) | 0.0036±0.00030 (A) |
| LCMS | 331-18b | 0.1152±0.01015 (A) | 0.0985±0.00493 (AB) | 0.0993±0.00341 (AB) | 0.0812±0.00420 (B) |
| LCMS | 331-18d | 0.0952±0.00649 (A) | 0.0760±0.00358 (B) | 0.0789±0.00361 (AB) | 0.0639±0.00461 (B) |
| LCMS | 375-19a | 0.0021±0.00015 (A) | 0.0017±0.00009 (AB) | 0.0018±0.00007 (AB) | 0.0016±0.00009 (B) |
| LCMS | 375-19b | 0.0004±0.00002 (A) | 0.0004±0.00005 (AB) | 0.0003±0.00003 (AB) | 0.0002±0.00002 (B) |
| LCMS | 375-19c | 0.0009±0.00008 (A) | 0.0008±0.00004 (A) | 0.0008±0.00004 (AB) | 0.0006±0.00004 (B) |
| HPLC | Total THC | 19.0353±1.05080 (A) | 15.6953±0.57558 (B) | 16.0541±0.56034 (B) | 14.7333±0.60475 (B) |
| HPLC | Total CBD | 0.0721±0.00442 (A) | 0.0586±0.00225 (B) | 0.0613±0.00204 (AB) | 0.0576±0.00284 (B) |
| HPLC | Total CBG | 0.8135±0.05527 (A) | 0.6712±0.03265 (AB) | 0.6223±0.02705 (BC) | 0.4934±0.03334 (C) |

**Table S7.** Effects of the drought treatments on the phytocannabinoid content of primary inflorescences of ‘Odem’ plants (Experiment 2). Phytocannabinoid concentrations (%, w/w) in the primary inflorescences of ‘Odem’ plants subjected to 4 different irrigation treatments, as measured at the end of Experiment 2. Samples were taken from the primary inflorescences. Values are means ± SE. Different letters represent significant differences between irrigation treatments, according to one-way ANOVA and Tukey’s HSD test ($P < 0.05$, $5 \leq N \leq 6$).

| Separation method | Phytocannabinoid | Control | Mild drought | Moderate drought | Severe drought |
|---|---|---|---|---|---|
| HPLC | THCA | 19.0926±0.53493 (A) | 16.978±0.2230(BC) | 17.808±0.4766(AB) | 15.741±0.338 (C) |
| HPLC | THC | 1.1635±0.13184 (A) | 0.9635±0.05838 (A) | 1.1800±0.09829 (A) | 1.2393±0.04843 (A) |
| HPLC | CBDA | 0.0765±0.00181 (A) | 0.0664±0.00121 (B) | 0.0722±0.00215 (AB) | 0.0665±0.00171 (B) |
| HPLC | CBGA | 0.6736±0.04546 (A) | 0.5844±0.01318 (AB) | 0.6154±0.04683 (A) | 0.4499±0.03726 (B) |
| HPLC | CBG | 0.0848±0.00240 (A) | 0.0803±0.00392 (A) | 0.0805±0.00431 (A) | 0.0641±0.00324 (B) |
| HPLC | CBDVA | 0.0536±0.00329 (A) | 0.0513±0.00230 (A) | 0.0544±0.00164 (A) | 0.0468±0.00116 (A) |
| HPLC | CBNA | 0.1009±0.00529 (A) | 0.1020±0.00152 (A) | 0.1085±0.00302 (A) | 0.1061±0.00565 (A) |
| HPLC | CBN | 0.0066±0.00069 (B) | 0.0075±0.00068 (B) | 0.0094±0.00088 (B) | 0.0129±0.00104 (A) |
| HPLC | CBCA | 0.3319±0.02262 (A) | 0.3040±0.01544 (A) | 0.3228±0.01851 (A) | 0.3400±0.02140 (A) |
| HPLC | CBC | 0.0367±0.00283 (B) | 0.0333±0.00110 (B) | 0.0458±0.00447 (AB) | 0.0548±0.00601 (A) |
| LCMS | CBGA-C4 | 0.0005±0.00003 (A) | 0.0004±0.00002 (A) | 0.0004±0.00002 (A) | 0.0004±0.00002 (A) |
| LCMS | CBGVA | 0.0003±0.00002 (A) | 0.0002±0.00001 (A) | 0.0002±0.00004 (A) | 0.00003±0.00003 (B) |
| LCMS | Sesqui-CBGA | 0.0009±0.00005 (AB) | 0.0009±0.00004 (B) | 0.0009±0.00003 (AB) | 0.0011±0.00009 (A) |
| LCMS | Sesqui-CBG | 1.4129±0.17748 (A) | 1.1428±0.08280 (A) | 1.3459±0.11725 (A) | 1.4848±0.09774 (A) |
| LCMS | THCA-C4 | 0.0422±0.00296 (A) | 0.0380±0.00211 (A) | 0.0390±0.00228 (A) | 0.0361±0.00342 (A) |
| LCMS | THCVA | 0.0071±0.00114 (A) | 0.0054±0.00047 (A) | 0.0067±0.00076 (A) | 0.0078±0.00106 (A) |
| LCMS | THCOA | 0.0131±0.00106 (A) | 0.0108±0.00047 (A) | 0.0116±0.00077 (A) | 0.0121±0.00061 (A) |
| LCMS | THCMA | 0.0115±0.00083 (A) | 0.0110±0.00068 (A) | 0.0120±0.00070 (A) | 0.0111±0.00104 (A) |
| LCMS | CBDA-C4 | 0.0001±0.00001 (A) | 0.0001±0.00002 (A) | 0.0001±0.00001 (A) | 0.0001±0.00003 (A) |
| LCMS | CBDM | 0.0361±0.00519 (A) | 0.0304±0.00239 (A) | 0.0429±0.00509 (A) | 0.0507±0.00730 (A) |
| LCMS | CBCA-C4 | 0.0005±0.00005 (A) | 0.0004±0.00005 (A) | 0.0005±0.00007 (A) | 0.0005±0.00007 (A) |
| LCMS | CBCO | 0.0100±0.00089 (B) | 0.0099±0.00046 (B) | 0.0119±0.00086 (AB) | 0.0148±0.00165 (A) |
| LCMS | CBNA-C4 | 0.0002±0.00001 (A) | 0.0002±0.00001 (A) | 0.0002±0.00001 (A) | 0.0002±0.00002 (A) |
| LCMS | CBNVA | 0.0006±0.00004 (A) | 0.0005±0.00001 (A) | 0.0005±0.00002 (A) | 0.0005±0.00004 (A) |
| LCMS | CBTA-1 | 0.0057±0.00028 (A) | 0.0054±0.00013 (A) | 0.0058±0.00033 (A) | 0.0062±0.00062 (A) |
| LCMS | CBT-1 | 0.0064±0.00047 (B) | 0.0064±0.00012 (B) | 0.0074±0.00050 (AB) | 0.0095±0.00106 (A) |
| LCMS | CBTA-3 | 0.0058±0.00039 (A) | 0.0052±0.00014 (AB) | 0.0055±0.00031 (AB) | 0.0045±0.00030 (B) |
| LCMS | CBT-3 | 0.0041±0.00036 (A) | 0.0042±0.00013 (A) | 0.0045±0.00036 (A) | 0.0052±0.00053 (A) |
| LCMS | 329-11b | 0.0006±0.00016 (B) | 0.0008±0.00035 (B) | 0.0010±0.00022 (AB) | 0.0024±0.00070 (A) |
| LCMS | 329-11d | 0.0022±0.00009 (A) | 0.0020±0.00016 (A) | 0.0022±0.00011 (A) | 0.0024±0.00033 (A) |
| LCMS | 373-12b | 0.0038±0.00073 (A) | 0.0050±0.00200 (A) | 0.0058±0.00165 (A) | 0.0092±0.00294 (A) |
| LCMS | 373-12c | 0.0011±0.00026 (A) | 0.0016±0.00080 (A) | 0.0017±0.00049 (A) | 0.0021±0.00070 (A) |
| LCMS | 373-12d | 0.0029±0.00022 (A) | 0.0027±0.00013 (A) | 0.0027±0.00013 (A) | 0.0023±0.00013 (A) |
| LCMS | 327-13a | 0.0002±0.00006 (A) | 0.0002±0.00002 (A) | 0.0002±0.00002 (A) | 0.0002±0.00002 (A) |

**Table. S8** continued on the next page

**Table. S8** continued from the previous page

| Separation method | Phytocannabinoid | Control | Mild drought | Moderate drought | Severe drought |
|---|---|---|---|---|---|
| LCMS | 327-13c | 0.0024±0.00019 (A) | 0.0018±0.00041 (A) | 0.0019±0.00044 (A) | 0.0021±0.00027 (A) |
| LCMS | 371-14a | 0.0004±0.00004 (A) | 0.0004±0.00002 (A) | 0.0004±0.00001 (A) | 0.0003±0.00002 (A) |
| LCMS | 371-14b | 0.0014±0.00011 (A) | 0.0013±0.00008 (A) | 0.0013±0.00005 (A) | 0.0011±0.00008 (A) |
| LCMS | 417-15a | 0.0031±0.00021 (A) | 0.0025±0.00003 (B) | 0.0029±0.00010 (AB) | 0.0024±0.00008 (B) |
| LCMS | 357-16a | 0.0106±0.00059 (A) | 0.0094±0.00032 (AB) | 0.0094±0.00035 (AB) | 0.0078±0.00041 (B) |
| LCMS | 313-16b | 0.0014±0.00013 (A) | 0.0012±0.00008 (A) | 0.0013±0.00011 (A) | 0.0012±0.00009 (A) |
| LCMS | 361-17a | 0.0027±0.00016 (B) | 0.0030±0.00009 (B) | 0.0034±0.00021 (AB) | 0.0041±0.00050 (A) |
| LCMS | 331-18b | 0.1121±0.00707 (A) | 0.1025±0.01020 (A) | 0.0996±0.00592 (A) | 0.0816±0.00574 (A) |
| LCMS | 331-18d | 0.0895±0.00733 (A) | 0.0763±0.00411 (AB) | 0.0799±0.00637 (AB) | 0.0612±0.00161 (B) |
| LCMS | 375-19a | 0.0020±0.00010 (A) | 0.0018±0.00004 (AB) | 0.0018±0.00008 (AB) | 0.0016±0.00005 (B) |
| LCMS | 375-19b | 0.0003±0.00004 (A) | 0.0003±0.00002 (A) | 0.0003±0.00003 (A) | 0.0002±0.00001 (A) |
| LCMS | 375-19c | 0.0008±0.00006 (A) | 0.0007±0.00002 (AB) | 0.0007±0.00005 (AB) | 0.0006±0.00004 (B) |
| HPLC | Total THC | 17.9077±0.34924 (A) | 15.8540±0.24362 (AB) | 16.7977±0.43234 (BC) | 15.0442±0.32887 (C) |
| HPLC | Total CBD | 0.0671±0.00158 (A) | 0.0582±0.00106 (B) | 0.0634±0.00189 (AB) | 0.0583±0.00150 (B) |
| HPLC | Total CBG | 0.6755±0.04062 (A) | 0.5928±0.01495 (AB) | 0.6202±0.04313 (A) | 0.4586±0.03378 (B) |

**Table S8.** Effects of the drought treatments on the phytocannabinoid contents of secondary inflorescences of 'Odem' plants (Experiment 2). Phytocannabinoid concentrations (%, w/w) in the secondary inflorescences of 'Odem' plants subjected to 4 different irrigation treatments, as measured at the end of Experiment 2. Samples were taken from the secondary inflorescences. Values are means ± SE. Different letters represent significant differences between irrigation treatments, according to one-way ANOVA and Tukey's HSD test ($P < 0.05$, $5 \leq N \leq 6$).

| Separation method | Phytocannabinoid | Control | Mild drought | Moderate drought | Severe drought |
|---|---|---|---|---|---|
| HPLC | CBGA | 1.2633±0.11576 (A) | 0.8248±0.09199 (B) | 0.7144±0.05868 (BC) | 0.4197±0.03950 (C) |
| HPLC | CBG | 0.0823±0.01056 (A) | 0.0552±0.00638 (B) | 0.0543±0.00409 (B) | 0.0323±0.00188 (B) |
| HPLC | CBDA | 0.0642±0.02698 (A) | 0.0818±0.00826 (A) | 0.0781±0.01513 (A) | 0.0836±0.02088 (A) |
| HPLC | THCA | 22.405±2.31612 (A) | 17.567±1.24962 (AB) | 18.686±0.88501 (AB) | 14.673±0.89214 (B) |
| HPLC | THC | 0.4389±0.04858 (A) | 0.3155±0.01580 (A) | 0.3543±0.01853 (A) | 0.3654±0.04252 (A) |
| HPLC | CBDVA | 0.0178±0.00155 (A) | 0.0167±0.00112 (A) | 0.0194±0.00119 (A) | 0.0151±0.00133 (A) |
| HPLC | CBCA | 0.1817±0.02318 (A) | 0.1394±0.00916 (A) | 0.1618±0.00492 (A) | 0.1557±0.00797 (A) |
| HPLC | CBC | 0.0169±0.00198 (A) | 0.0115±0.00079 (AB) | 0.0132±0.00083 (B) | 0.0131±0.0012 (AB) |
| HPLC | CBNA | 0.0338±0.00223 (A) | 0.0286±0.00097 (AB) | 0.0312±0.00133 (AB) | 0.0252±0.00151 (B) |
| HPLC | CBN | 0.0013±0.0001 (AB) | 0.0009±0.00010 (B) | 0.0012±0.00007 (AB) | 0.0017±0.00028 (A) |
| LCMS | THCA-C4 | 0.0207±0.00238 (A) | 0.0154±0.00172 (AB) | 0.0157±0.00095 (AB) | 0.0119±0.00078 (B) |
| LCMS | THCVA | 0.0378±0.00562 (A) | 0.0272±0.00262 (AB) | 0.0282±0.00110 (AB) | 0.0213±0.00076 (B) |
| LCMS | THCOA | 0.0142±0.00205 (A) | 0.0114±0.00148 (A) | 0.0134±0.00163 (A) | 0.0147±0.00161 (A) |
| LCMS | THCMA | 0.0210±0.00247 (A) | 0.0179±0.00236 (A) | 0.0180±0.00136 (A) | 0.0148±0.00135 (A) |
| LCMS | Sesqui-CBGA | 0.0006±0.00007 (A) | 0.0004±0.00004 (A) | 0.0005±0.00005 (A) | 0.0005±0.00008 (A) |
| LCMS | Sesqui-CBG | 0.0074±0.00090 (A) | 0.0058±0.00075 (AB) | 0.0063±0.00071 (AB) | 0.0041±0.00032 (B) |
| LCMS | CBTA-1 | 0.0026±0.00014 (A) | 0.0020±0.00008 (B) | 0.0022±0.00007 (AB) | 0.0020±0.00011 (B) |
| LCMS | CBTA-3 | 0.0016±0.00015 (A) | 0.0014±0.00008 (A) | 0.0015±0.00009 (A) | 0.0012±0.00012 (A) |
| LCMS | CBT-3 | 0.0007±0.00010 (A) | 0.0005±0.00003 (A) | 0.0006±0.00002 (A) | 0.0006±0.00008 (A) |
| LCMS | 373-12b | 0.0011±0.00015 (A) | 0.0008±0.00005 (A) | 0.0009±0.00007 (A) | 0.0008±0.00006 (A) |
| LCMS | 373-12c | 0.0002±0.00002 (A) | 0.0001±0.00001 (A) | 0.0001±0.00001 (A) | 0.0001±0.00001 (A) |
| LCMS | 371-14b | 0.0005±0.00003 (A) | 0.0004±0.00002 (AB) | 0.0004±0.00003 (A) | 0.0003±0.00003 (B) |
| LCMS | 417-15a | 0.0014±0.00017 (A) | 0.0011±0.00006 (A) | 0.0011±0.00011 (A) | 0.0008±0.00009 (A) |
| LCMS | 373-15b | 0.0188±0.00219 (A) | 0.0148±0.00142 (AB) | 0.0162±0.00066 (AB) | 0.0109±0.00072 (B) |
| LCMS | 357-16a | 0.0020±0.00029 (A) | 0.0017±0.00019 (A) | 0.0019±0.00015 (A) | 0.0015±0.00022 (A) |
| LCMS | 331-18b | 0.0546±0.00699 (A) | 0.0443±0.00476 (AB) | 0.0474±0.00154 (AB) | 0.0321±0.00206 (B) |
| LCMS | 331-18c | 0.0014±0.00020 (A) | 0.0010±0.00009 (AB) | 0.0011±0.00005 (AB) | 0.0007±0.00005 (B) |
| LCMS | 331-18d | 0.0553±0.00854 (A) | 0.0414±0.00386 (AB) | 0.0434±0.00166 (AB) | 0.0305±0.00249 (B) |
| LCMS | 375-19a | 0.0012±0.00017 (A) | 0.0009±0.00006 (AB) | 0.0010±0.00004 (AB) | 0.0008±0.00005 (B) |
| LCMS | Total THC | 20.088±2.07385 (A) | 15.721±1.11057 (AB) | 16.742±0.78117 (AB) | 13.234±0.77499 (B) |
| LCMS | Total CBD | 0.0563±0.02366 (A) | 0.0718±0.00724 (A) | 0.0685±0.01327 (A) | 0.0734±0.01831 (A) |
| LCMS | Total CBG | 1.1902±0.11175 (A) | 0.7786±0.08425 (B) | 0.6808±0.05466 (BC) | 0.4004±0.03651 (C) |

**Table S9.** Effects of the drought treatments on the phytocannabinoid contents of primary inflorescences of ‘MVA’ plants (Experiment 3). Phytocannabinoid concentrations (%, w/w) in the primary inflorescences of ‘MVA’ plants subjected to 4 different irrigation treatments, as measured at the end of Experiment 3. Samples were taken from the primary inflorescences. Values are means ± SE. Different letters represent significant differences between irrigation treatments, according to one-way ANOVA and Tukey’s HSD test ($P < 0.05$, $5 \leq N \leq 6$).

| Separation method | Phytocannabinoid | Control | Mild drought | Moderate drought | Severe drought |
|---|---|---|---|---|---|
| HPLC | CBGA | 1.2161±0.09770 (A) | 0.8077±0.09782 (B) | 0.5703±0.07907 (BC) | 0.4346±0.04083 (C) |
| HPLC | CBG | 0.0737±0.00615 (A) | 0.0525±0.00504 (B) | 0.0406±0.00599 (B) | 0.0332±0.00168 (B) |
| HPLC | THCA | 21.7113±1.6123 (A) | 17.3936±0.96036 (AB) | 15.1615±1.18447 (B) | 14.4774±0.63477 (B) |
| HPLC | THC | 0.4120±0.04089 (A) | 0.3347±0.00805 (AB) | 0.2955±0.01033 (B) | 0.3685±0.03954 (AB) |
| HPLC | THCA-C4 | 0.0177±0.00104 (A) | 0.0134±0.00120 (AB) | 0.0116±0.00128 (B) | 0.0117±0.00053 (B) |
| HPLC | CBDA | 0.0958±0.00573 (A) | 0.0812±0.00625 (A) | 0.0733±0.00565 (A) | 0.1011±0.01142 (A) |
| HPLC | CBDVA | 0.0175±0.00136 (A) | 0.0164±0.00092 (A) | 0.0163±0.00135 (A) | 0.0143±0.00107 (A) |
| HPLC | CBCA | 0.1749±0.01188 (A) | 0.1440±0.00594 (AB) | 0.1272±0.00699 (B) | 0.1568±0.00795 (AB) |
| HPLC | CBC | 0.0172±0.00087 (A) | 0.0131±0.00107 (B) | 0.0116±0.00035 (B) | 0.0150±0.00120 (AB) |
| HPLC | CBNA | 0.0344±0.00245 (A) | 0.0289±0.00119 (AB) | 0.0265±0.00175 (B) | 0.0256±0.00168 (B) |
| HPLC | CBN | 0.0012±0.00010 (A) | 0.0011±0.00015 (A) | 0.0012±0.00009 (A) | 0.0014±0.00020 (A) |
| LCMS | Sesqui-CBGA | 0.0005±0.00008 (A) | 0.0004±0.00004 (A) | 0.0004±0.00003 (A) | 0.0005±0.00005 (A) |
| LCMS | Sesqui-CBG | 0.0073±0.00135 (A) | 0.0054±0.00056 (AB) | 0.0051±0.00079 (AB) | 0.0037±0.00013 (B) |
| LCMS | THCVA | 0.0319±0.00166 (A) | 0.0250±0.00179 (B) | 0.0201±0.00195 (B) | 0.0209±0.00085 (B) |
| LCMS | THCOA | 0.0135±0.00081 (A) | 0.0111±0.00149 (A) | 0.0108±0.00135 (A) | 0.0136±0.00216 (A) |
| LCMS | THCMA | 0.0216±0.00228 (A) | 0.0162±0.00081 (AB) | 0.0152±0.00076 (B) | 0.0127±0.00169 (B) |
| LCMS | CBTA-1 | 0.0023±0.00015 (A) | 0.0020±0.00011 (A) | 0.0019±0.00011 (A) | 0.0019±0.00012 (A) |
| LCMS | CBTA-3 | 0.0015±0.00017 (A) | 0.0013±0.00007 (A) | 0.0015±0.00018 (A) | 0.0010±0.00018 (A) |
| LCMS | CBT-3 | 0.0006±0.00003 (A) | 0.0005±0.00004 (A) | 0.0004±0.00002 (A) | 0.0006±0.00012 (A) |
| LCMS | 373-12b | 0.0010±0.00006 (A) | 0.0008±0.00004 (A) | 0.0007±0.00007 (A) | 0.0008±0.00013 (A) |
| LCMS | 373-12c | 0.0001±0.00001 (A) | 0.0001±0.00001 (A) | 0.0001±0.00002 (A) | 0.0001±0.00002 (A) |
| LCMS | 371-14b | 0.0004±0.00003 (A) | 0.0004±0.00002 (A) | 0.0004±0.00003 (AB) | 0.0003±0.00003 (B) |
| LCMS | 417-15a | 0.0014±0.00013 (A) | 0.0012±0.00006 (AB) | 0.0010±0.00009 (B) | 0.0008±0.00005 (B) |
| LCMS | 373-15b | 0.0178±0.00204 (A) | 0.0141±0.00079 (AB) | 0.0119±0.00069 (B) | 0.0101±0.00071 (B) |
| LCMS | 357-16a | 0.0020±0.00023 (A) | 0.0017±0.00012 (AB) | 0.0017±0.00009 (AB) | 0.0013±0.00014 (B) |
| LCMS | 331-18b | 0.0505±0.00548 (A) | 0.0394±0.00209 (AB) | 0.0333±0.00268 (B) | 0.0285±0.00191 (B) |
| LCMS | 331-18c | 0.0013±0.00013 (A) | 0.0010±0.00006 (B) | 0.0008±0.00008 (B) | 0.0007±0.00006 (B) |
| LCMS | 331-18d | 0.0484±0.00245 (A) | 0.0383±0.00315 (AB) | 0.0313±0.00306 (B) | 0.0289±0.00302 (B) |
| LCMS | 375-19a | 0.0011±0.00009 (A) | 0.0009±0.00006 (B) | 0.0008±0.00005 (B) | 0.0007±0.00005 (B) |
| HPLC | Total THC | 19.4528±1.4420 (A) | 15.5889±0.84233 (AB) | 13.5921±1.04511 (B) | 13.0651±0.56815 (B) |
| HPLC | Total CBD | 0.0840±0.00502 (A) | 0.0712±0.00548 (A) | 0.0643±0.00495 (A) | 0.0887±0.01002 (A) |
| HPLC | Total CBG | 1.1402±0.09113 (A) | 0.7608±0.09042 (B) | 0.5407±0.07465 (BC) | 0.4143±0.03725 (C) |

**Table S10.** Effects of the drought treatment on the phytocannabinoid contents of secondary inflorescences of ‘MVA’ plants (Experiment 3). Phytocannabinoid concentrations (%, w/w) in the secondary inflorescences of ‘MVA’ plants subjected to 4 different irrigation treatments, as measured at the end of Experiment 3. Samples were taken from the secondary inflorescences. Values are means ± SE. Different letters represent significant differences between irrigation treatments, according to one-way ANOVA and Tukey’s HSD test ($P < 0.05$, $5 \leq N \leq 6$).

| Separation method | Phytocannabinoid | Control | Mild drought | Moderate drought | Severe drought |
|---|---|---|---|---|---|
| HPLC | THCA | 21.529±0.790 (A) | 19.912±1.408 (AB) | 17.687±0.502 (B) | 16.999±0.118 (B) |
| HPLC | THC | 0.279±0.025 (A) | 0.245±0.035 (A) | 0.218±0.014 (A) | 0.248±0.023 (A) |
| HPLC | CBGA | 0.570±0.049 (A) | 0.411±0.049 (AB) | 0.313±0.022 (B) | 0.280±0.052 (B) |
| HPLC | CBDA | 0.027±0.007 (A) | 0.025±0.008 (A) | 0.006±0.006 (A) | 0.007±0.008 (A) |
| HPLC | THCV | 0.146±0.016 (A) | 0.148±0.012 (A) | 0.171±0.014 (A) | 0.123±0.016 (A) |
| HPLC | Total THC | 21.808±0.812 (A) | 20.157±1.439 (AB) | 17.905±0.513 (B) | 17.758±0.561 (B) |

**Table S11.** Effects of the drought treatments on the phytocannabinoid contents of primary inflorescences of ‘Odem’ plants (Experiment 4). Phytocannabinoid concentrations (%, w/w) in the primary inflorescences of ‘Odem’ plants subjected to 4 different irrigation treatments, as measured at the end of Experiment 4. Samples were taken from the primary inflorescences. Values are means ± SE. Different letters represent significant differences between irrigation treatments, according to one-way ANOVA and Tukey’s HSD test ($P < 0.05$, $5 \leq N \leq 6$).

| Separation method | Phtocannabinoid | Control | Mild drought | Moderate drought | Severe drought |
|---|---|---|---|---|---|
| HPLC | THCA | 20.60±0.75 (A) | 19.69±0.66 (AB) | 17.31±0.76 (B) | 17.95±1.08 (AB) |
| HPLC | THC | 0.23±0.01 (A) | 0.24±0.03 (A) | 0.22±0.02 (A) | 0.26±0.03 (A) |
| HPLC | CBGA | 0.54±0.04 (A) | 0.41±0.04 (AB) | 0.29±0.02 (B) | 0.31±0.04 (B) |
| HPLC | CBDA | 0.03±0.00 (A) | 0.02±0.01 (A) | 0.01±0.01 (A) | 0.01±0.01 (A) |
| HPLC | THCV | 0.13±0.02 (A) | 0.15±0.02 (A) | 0.15±0.01 (A) | 0.12±0.01 (A) |
| HPLC | TOTAL THC | 20.84±0.76 (A) | 19.93±0.68 (A) | 17.53±0.77 (A) | 18.21±1.10 (A) |

**Table S12.** Effects of the drought treatments on the phytocannabinoid contents of secondary inflorescences of ‘Odem’ plants (Experiment 4). Phytocannabinoid concentrations (%, w/w) in the secondary inflorescences of ‘Odem’ plants subjected to 4 different irrigation treatments, as measured at the end of Experiment 4. Samples were taken from the secondary inflorescences. Values are means ± SE. Different letters represent significant differences between irrigation treatments, according to one-way ANOVA and Tukey’s HSD test ($P < 0.05$, $5 \leq N \leq 6$).

| Separation method | Phytocannabinoid | Control | Mild drought | Moderate drought | Severe drought |
|---|---|---|---|---|---|
| HPLC | THCA | 19.17±0.720 (A) | 19.98±0.511 (A) | 18.60±0.741 (A) | 17.50±0.826 (A) |
| HPLC | THC | 0.13±0.026 (A) | 0.13±0.011 (A) | 0.12±0.011 (A) | 0.11±0.007 (A) |
| HPLC | CBGA | 0.56±0.032 (A) | 0.52±0.040 (A) | 0.48±0.036 (A) | 0.43±0.043 (A) |
| HPLC | CBG | 0.11±0.007 (AB) | 0.12±0.006 (A) | 0.11±0.010 (AB) | 0.08±0.008 (B) |
| HPLC | Total THC | 19.30±0.742 (A) | 20.11±0.510 (A) | 18.72±0.747 (A) | 17.61±0.830 (A) |

**Table S13.** Effects of the drought treatments on the phytocannabinoid content of primary inflorescences of Line 187 plants (Experiment 5). Phytocannabinoid concentrations (%, w/w) in the primary inflorescences of plants of Line 187 that were subjected to 4 different irrigation treatments, as measured at the end of Experiment 5. Samples were taken from the primary inflorescences. Values are means ± SE. Different letters represent significant differences between irrigation treatments, according to one-way ANOVA and Tukey's HSD test ($P < 0.05$, $5 \leq N \leq 6$).

| Separation method | Phytocannabinoid | Control | Mild drought | Moderate drought | Severe drought |
|---|---|---|---|---|---|
| HPLC | THCA | 21.40±0.67 (A) | 21.49±0.23 (A) | 19.08±0.92 (A) | 18.88±1.23 (A) |
| HPLC | THC | 0.13±0.01 (A) | 0.14±0.01 (A) | 0.12±0.01 (A) | 0.13±0.01 (A) |
| HPLC | CBGA | 0.62±0.04 (A) | 0.59±0.02 (A) | 0.51±0.04 (A) | 0.49±0.06 (A) |
| HPLC | CBG | 0.13±0.01 (A) | 0.13±0.01 (A) | 0.11±0.01 (A) | 0.10±0.01 (A) |
| HPLC | Total THC | 21.53±0.67 (A) | 21.62±0.24 (A) | 19.19±0.92 (A) | 19.02±1.24 (A) |

**Table S14.** Effects of the drought treatments on the phytocannabinoid contents of secondary inflorescences of Line 187 plants (Experiment 5). Phytocannabinoid concentrations (%, w/w) in the secondary inflorescences of plants of Line 187 that were subjected to 4 different irrigation treatments, as measured at the end of Experiment 5. Samples were taken from the secondary inflorescences. Values are means ± SE. Different letters represent significant differences between irrigation treatments, according to one-way ANOVA and Tukey's HSD test ($P < 0.05$, $5 \leq N \leq 6$).

| Terpenoid | Control | Mild drought | Moderate Drought | Severe drought |
|---|---|---|---|---|
| α-Pinene | 1367.64±192.99 (A) | 1410.20±220.24 (A) | 1521.45±325.96 (A) | 1773.80±342.20 (A) |
| Camphene | 108.97±15.53 (A) | 104.70±7.94 (A) | 95.84±13.37 (A) | 109.85±16.04 (A) |
| β-Pinene | 576.96±71.34 (A) | 573.56±73.76 (A) | 595.49±105.73 (A) | 662.88±115.05 (A) |
| β-Myrcene | 1370.99±194.10 (A) | 1278.84±155.87 (A) | 1087.20±95.45 (A) | 980.50±53.19 (A) |
| α-Phellandrene | 4.93±0.79 (A) | 4.59±0.56 (A) | 3.74±0.23 (A) | 3.30±0.49 (A) |
| α-Terpinene | 14.35±1.61 (A) | 14.22±1.26 (A) | 12.41±0.80 (A) | 12.80±1.21 (A) |
| Limonene | 1156.42±148.64 (A) | 1105.19±61.45 (A) | 867.35±65.04 (A) | 918.56±110.41 (A) |
| β-Phellandrene | 28.67±3.58 (A) | 27.37±3.43 (A) | 23.11±2.23 (A) | 21.34±3.29 (A) |
| cis-Ocimene | 10.44±1.52 (A) | 11.13±1.15 (A) | 9.12±0.67 (A) | 9.81±1.58 (A) |
| Eucalyptol | 10.98±2.00 (A) | 14.28±2.15 (A) | 12.62±1.19 (A) | 13.61±2.22 (A) |
| trans-Ocimene | 626.55±112.86 (A) | 630.20±22.66 (A) | 509.98±33.24 (A) | 595.77±77.10 (A) |
| γ-Terpinene | 16.22±1.93 (A) | 15.73±1.55 (A) | 13.86±1.06 (A) | 14.83±1.63 (A) |
| Terpinolene | 55.78±6.86 (A) | 54.15±3.30 (A) | 43.83±3.00 (A) | 50.40±4.33 (A) |
| cis-Linalool oxide | 178.94±14.05 (A) | 170.43±14.87 (A) | 135.85±14.76 (A) | 156.65±10.17 (A) |
| Sabinene hydrate | 126.77±10.08 (A) | 134.91±5.22 (A) | 99.98±14.93 (A) | 129.12±16.02 (A) |
| trans-Linalool oxide | 12.69±1.26 (A) | 11.67±1.52 (A) | 9.40±1.03 (A) | 9.62±1.43 (A) |
| Fenchone | 29.18±3.38 (A) | 25.01±2.37 (A) | 20.73±1.57 (A) | 23.69±0.79 (A) |
| Linalool | 521.53±53.65 (AB) | 562.87±38.49 (A) | 399.79±24.17 (B) | 370.77±42.00 (B) |
| C10H18O-154(93/79/99/121)-1 | 75.83±9.92 (A) | 59.51±10.53 (AB) | 37.81±6.64 (B) | 47.61±4.31 (AB) |
| Fenchol | 364.31±35.09 (A) | 344.16±23.86 (A) | 269.84±16.83 (A) | 291.00±16.49 (A) |
| Camphor | 2.55±0.30 (A) | 2.96±0.27 (A) | 2.49±0.29 (A) | 2.91±0.39 (A) |
| Terpinen-4-ol | 22.53±3.01 (A) | 26.90±3.06 (A) | 21.51±2.86 (A) | 21.04±2.58 (A) |
| Borneol | 129.99±8.51 (A) | 134.69±7.80 (A) | 118.13±5.67 (A) | 119.37±6.25 (A) |
| α-Terpineol | 209.84±21.63 (A) | 213.54±19.66 (A) | 165.75±11.81 (A) | 159.89±16.02 (A) |
| Nerol | 4.59±1.01 (A) | 6.17±0.63 (A) | 3.55±0.48 (A) | 3.87±0.92 (A) |
| Geraniol | 27.63±6.33 (A) | 22.02±5.08 (A) | 9.76±3.76 (A) | 19.93±2.10 (A) |
| Bornyl acetate | 10.39±1.47 (B) | 13.81±1.70 (AB) | 13.72±2.44 (AB) | 19.84±2.17 (A) |
| α-Cubebene | 3.56±0.42 (A) | 5.22±0.68 (AB) | 5.04±0.73 (AB) | 7.02±0.64 (A) |
| Ylangene | 9.42±1.10 (A) | 11.97±1.62 (A) | 11.79±1.54 (A) | 14.39±1.05 (A) |
| α-Copaene | 6.46±0.70 (A) | 7.92±1.32 (A) | 7.93±1.12 (A) | 9.56±0.76 (A) |
| C15H24-204(105/(120+119)/161) | 4.03±0.51 (A) | 5.67±0.75 (AB) | 5.47±0.76 (AB) | 7.50±0.58 (A) |
| 7-epi-Sesquithujene | 12.78±1.21 (A) | 14.56±2.33 (A) | 14.23±2.01 (A) | 19.73±1.60 (A) |
| β-Isocomene | 24.51±2.45 (A) | 20.71±2.32 (A) | 25.51±3.37 (A) | 24.25±1.52 (A) |
| Sesquithujene | 84.65±4.99 (A) | 69.68±10.73 (A) | 68.87±12.41 (A) | 71.74±12.77 (A) |
| α-Cedrene | 2.45±0.04 (A) | 7.81±3.41 (A) | 2.56±0.13 (A) | 3.21±0.36 (A) |

**Table. S15** continued on the next page

**Table. S15** continued from the previous page

| Terpenoid | Control | Mild drought | Moderate Drought | Severe drought |
|---|---|---|---|---|
| cis-α-Bergamotene | 59.27±4.78 (A) | 56.29±3.16 (A) | 57.49±1.96 (A) | 62.01±3.37 (A) |
| C15H24-204(105/91/147/189) | 16.50±1.06 (A) | 18.40±1.92 (A) | 18.61±2.01 (A) | 22.22±1.13 (A) |
| α-Santalene | 40.28±1.84 (A) | 41.09±2.99 (A) | 40.52±3.18 (A) | 47.15±2.56 (A) |
| β-Caryophyllene | 1447.18±163.69 (A) | 1758.19±194.09 (A) | 1835.88±170.53 (A) | 1960.19±145.44 (A) |
| C15H24-204(120/91/79/69) | 4.33±0.74 (A) | 5.30±0.61 (A) | 5.45±0.62 (A) | 7.03±1.20 (A) |
| γ-Elemene | 79.23±1.87 (A) | 39.76±1.54 (B) | 56.60±7.36 (B) | 55.09±4.02 (B) |
| trans-α-Bergamotene | 817.22±63.61 (A) | 803.43±107.83 (A) | 841.27±85.45 (A) | 984.50±59.68 (A) |
| α-Guaiene | 798.33±77.24 (A) | 865.04±115.73 (A) | 940.64±121.67 (A) | 1062.03±74.07 (A) |
| Guaia-6,9-diene | 10.50±0.53 (A) | 10.42±1.65 (A) | 11.49±1.28 (A) | 12.73±0.71 (A) |
| C15H24-204(69/91/105/161) | 5.84±0.18 (A) | 5.57±0.70 (A) | 6.04±0.66 (A) | 7.70±0.44 (A) |
| β-Santalene | 16.68±1.02 (A) | 16.12±2.08 (A) | 16.00±1.93 (A) | 20.23±1.40 (A) |
| α-Himachalene | 5.10±0.42 (A) | 5.45±0.69 (A) | 5.97±0.61 (A) | 6.33±0.39 (A) |
| C15H24-204(105/91/133/161/189)-1 | 12.01±1.02 (A) | 13.34±1.63 (A) | 13.63±1.71 (A) | 16.54±0.97 (A) |
| α-Humulene | 750.98±78.55 (A) | 851.29±92.08 (A) | 899.67±90.09 (A) | 1006.39±63.29 (A) |
| trans-β-Farnesene | 154.70±8.73 (A) | 136.73±15.60 (A) | 135.93±15.56 (A) | 157.88±8.00 (A) |
| C15H24-204(105)-1 | 41.33±2.91 (A) | 42.98±5.06 (A) | 46.19±5.86 (A) | 54.52±5.19 (A) |
| C15H24-204(133/189)-1 | 56.28±3.87 (A) | 60.64±6.30 (A) | 65.57±7.93 (A) | 75.33±7.05 (A) |
| γ-Muurelene | 73.11±4.99 (A) | 75.96±8.73 (A) | 79.01±9.45 (A) | 95.19±5.38 (A) |
| α-Amorphene | 47.24±2.96 (A) | 49.55±4.90 (A) | 53.00±5.78 (A) | 62.94±3.05 (A) |
| γ-Curcumene | 21.31±0.75 (A) | 20.89±1.26 (A) | 21.41±1.64 (A) | 23.99±0.92 (A) |
| C15H24-204(189/133)-2 | 95.39±6.44 (A) | 103.28±11.54 (A) | 114.18±13.94 (A) | 131.63±11.95 (A) |
| 4,5-di-epi-aristolochene | 23.15±2.49 (A) | 20.11±1.71 (A) | 19.43±2.09 (A) | 22.34±3.01 (A) |
| Sesquisabinene | 23.52±3.60 (A) | 22.12±3.43 (A) | 21.41±2.91 (A) | 21.47±1.63 (A) |
| α-Selinene | 105.33±7.27 (A) | 108.32±10.61 (A) | 116.50±13.59 (A) | 134.70±11.54 (A) |
| α-Curcumene | 9.60±0.41 (A) | 9.01±0.84 (A) | 9.68±0.78 (A) | 9.80±0.41 (A) |
| Valencene | 44.73±2.36 (A) | 45.75±3.93 (A) | 47.82±4.78 (A) | 53.92±2.36 (A) |
| β-Selinene | 442.49±39.62 (A) | 479.18±53.05 (A) | 533.41±63.30 (A) | 585.59±42.11 (A) |
| α-Zingiberene+C15H24-204(105) | 32.87±1.03 (A) | 31.29±2.22 (A) | 30.92±2.50 (A) | 33.31±0.66 (A) |
| δ-Guaiene | 1161.79±103.05 (A) | 1184.72±120.43 (A) | 1304.05±144.22 (A) | 1399.24±87.06 (A) |
| β-Bisabolene | 28.65±0.59 (A) | 27.42±0.70 (A) | 26.94±1.12 (A) | 28.35±0.76 (A) |
| C15H24-204(similar to Germarcene B) | 32.28±1.36 (A) | 30.72±2.06 (A) | 30.48±2.58 (A) | 33.26±0.81 (A) |
| α-Farnesene | 148.99±12.39 (A) | 138.44±17.15 (A) | 127.56±12.75 (A) | 136.42±5.31 (A) |
| C15H24-204(161/105)-Eremophilene | 32.12±1.40 (A) | 31.16±1.99 (A) | 30.98±2.58 (A) | 33.80±0.85 (A) |
| δ-Cadinene-C15H24-204(119/161/105/134) | 16.18±0.48 (A) | 16.29±0.51 (A) | 17.00±0.47 (A) | 17.49±0.57 (A) |
| Sesquicineole | 13.53±0.71 (A) | 12.21±1.09 (A) | 12.78±1.42 (A) | 15.25±0.64 (A) |

**Table. S15** continued on the next page

**Table. S15** continued from the previous page

| Terpenoid | Control | Mild drought | Moderate Drought | Severe drought |
|---|---|---|---|---|
| C15H24-204(105)-2 | 36.82±1.67 (A) | 35.16±2.27 (A) | 35.19±3.04 (A) | 38.07±0.90 (A) |
| cis-Calamenene-204(159) | 3.34±0.12 (A) | 3.31±0.24 (A) | 3.47±0.25 (A) | 3.74±0.20 (A) |
| β-Sesquiphellandrene | 24.65±0.52 (A) | 23.80±0.63 (A) | 23.35±0.66 (A) | 24.11±0.66 (A) |
| C15H24-204(189/133)-3 | 193.33±12.20 (A) | 186.03±17.36 (A) | 200.89±22.09 (A) | 206.80±8.81 (A) |
| C15H24-204(161/133/105) | 4798.09±377.89 (A) | 4782.21±454.86 (A) | 4857.75±443.96 (A) | 5202.20±203.91 (A) |
| Selina-3,7(11)-diene | 7431.03±632.60 (A) | 7213.84±635.47 (A) | 7712.23±722.61 (A) | 7587.64±251.43 (A) |
| trans-α-Bisabolene | 45.13±1.17 (A) | 39.27±2.38 (A) | 38.92±2.95 (A) | 43.76±1.51 (A) |
| C15H24-202(202/131/145/159) | 4.66±0.17 (A) | 5.12±0.36 (A) | 4.77±0.49 (A) | 5.41±0.54 (A) |
| Cadala-1(10),3,8-triene-204(157/142) | 48.87±2.62 (A) | 53.16±2.20 (A) | 54.72±4.02 (A) | 60.19±4.28 (A) |
| Germacrene B | 45.91±6.13 (B) | 42.49±3.33 (B) | 52.72±7.34 (B) | 77.88±6.08 (A) |
| C15H24-204(189/133)-4 | 48.94±2.73 (A) | 46.40±2.20 (A) | 44.00±3.35 (A) | 44.49±1.66 (A) |
| trans-Nerolidol | 2856.84±277.10 (A) | 2791.83±185.67 (A) | 2920.34±346.94 (A) | 2976.48±267.32 (A) |
| Caryophyllene oxide | 33.75±0.52 (A) | 38.76±1.99 (A) | 36.59±2.02 (A) | 39.83±1.87 (A) |
| C15H22-202(178/163) | 13.09±0.39 (A) | 12.19±1.24 (A) | 11.16±1.51 (A) | 13.10±0.82 (A) |
| C15H22-202(187/202)-1 | 36.12±0.88 (A) | 40.34±2.45 (A) | 33.40±3.51 (A) | 35.34±1.06 (A) |
| C15H26O-222(93/69/41) | 9.51±0.28 (A) | 10.62±1.40 (A) | 9.86±1.50 (A) | 11.91±0.90 (A) |
| C15H24 -204(123/81) | 5.70±0.42 (B) | 6.90±0.72 (AB) | 6.68±0.96 (AB) | 9.10±0.89 (A) |
| Humulene oxide II | 12.77±0.18 (A) | 14.55±1.52 (A) | 13.70±1.75 (A) | 16.15±0.98 (A) |
| C15H26O-222(59/161/91)-1 | 35.51±3.88 (A) | 31.91±2.58 (A) | 26.55±2.94 (A) | 25.74±0.90 (A) |
| C15H26O-222(105/59/161)-1 | 19.52±1.72 (A) | 15.65±0.86 (AB) | 12.28±2.68 (B) | 13.20±0.91 (AB) |
| C15H22-202(187) | 2.46±0.28 (A) | 3.17±0.41 (A) | 2.72±0.38 (A) | 3.52±0.48 (A) |
| C15H22-202(187/202)-2 | 8.60±0.89 (A) | 11.28±1.53 (A) | 9.86±1.61 (A) | 13.07±1.68 (A) |
| β-Eudesmol | 72.56±5.03 (A) | 76.88±3.97 (A) | 73.34±6.76 (A) | 76.45±2.06 (A) |
| C15H24O-(similar Caryophyllene oxide)-1 | 22.42±2.09 (A) | 22.40±1.20 (A) | 20.87±1.96 (A) | 21.05±0.16 (A) |
| α-Bisabolol | 30.86±2.61 (A) | 34.23±2.13 (A) | 34.29±4.70 (A) | 42.03±3.04 (A) |
| Juniper camphor | 6.65±0.55 (B) | 9.02±0.46 (AB) | 8.16±0.83 (AB) | 9.25±0.52 (A) |
| Total MONO,% | 0.71±0.08 (A) | 0.70±0.05 (A) | 0.61±0.05 (A) | 0.65±0.07 (A) |
| Total Sesqui,% | 2.27±0.18 (A) | 2.29±0.20 (A) | 2.40±0.22 (A) | 2.52±0.06 (A) |
| Total terpenoids % | 2.98±0.25 (A) | 2.98±0.22 (A) | 3.01±0.24 (A) | 3.17±0.09 (A) |

**Table S15.** Effects of the drought treatments on terpenoid concentrations in primary inflorescences of ‘Odem’ plants (Experiment 1). Terpenoid concentrations (mg/L; ppm) in the primary inflorescences of ‘Odem’ plants subjected to 4 different irrigation treatments, as measured at the end of Experiment 1. Samples were taken from the primary inflorescences. Values are means ± SE. Different letters represent significant differences between irrigation treatments, according to one-way ANOVA and Tukey’s HSD test ($P < 0.05$, $5 \leq N \leq 6$).

| Terpenoid | Control | Mild drought | Moderate drought | Severe drought' |
|---|---|---|---|---|
| α-Pinene | 1498.61±100.762 (A) | 1716.47±303.555 (A) | 1649.66±385.795 (A) | 2208.50±428.464 (A) |
| Camphene | 128.92±7.378 (A) | 125.93±17.742 (A) | 122.35±20.658 (A) | 170.53±38.624 (A) |
| β-Pinene | 707.52±51.170 (A) | 688.34±107.428 (A) | 797.65±124.740 (A) | 869.63±141.997 (A) |
| β-Myrcene | 1295.44±235.333 (A) | 1174.57±75.074 (A) | 1059.47±75.212 (A) | 951.48±47.376 (A) |
| α-Phellandrene | 4.25±0.676 (A) | 4.23±0.174 (A) | 3.61±0.386 (A) | 3.68±0.712 (A) |
| α-Terpinene | 14.35±0.929 (A) | 14.02±0.628 (A) | 12.91±1.037 (A) | 13.54±2.163 (A) |
| Limonene | 1051.36±132.125 (A) | 967.97±42.054 (A) | 845.50±69.916 (A) | 880.25±98.194 (A) |
| β-Phellandrene | 25.74±3.914 (A) | 24.93±1.215 (A) | 20.36±2.875 (A) | 52.15±26.300 (A) |
| cis-Ocimene | 8.52±1.572 (A) | 9.34±0.626 (A) | 8.05±0.867 (A) | 6.86±1.035 (A) |
| Eucalyptol | 9.80±1.391 (A) | 11.59±0.768 (A) | 11.13±1.990 (A) | 10.63±1.609 (A) |
| trans-Ocimene | 505.05±125.041 (A) | 542.22±38.533 (A) | 447.77±43.602 (A) | 343.05±58.843 (A) |
| γ-Terpinene | 15.90±1.085 (A) | 15.24±0.759 (A) | 14.43±1.141 (A) | 14.73±2.145 (A) |
| Terpinolene | 54.48±3.878 (A) | 52.30±3.177 (A) | 45.33±4.090 (A) | 49.33±7.466 (A) |
| cis-Linalool oxide | 182.85±17.047 (A) | 155.93±13.003 (A) | 133.71±16.249 (A) | 152.35±22.891 (A) |
| Sabinene hydrate | 133.81±8.039 (A) | 125.14±7.174 (A) | 92.34±21.876 (A) | 123.79±20.322 (A) |
| trans-Linalool oxide | 13.59±1.007 (A) | 11.47±0.863 (A) | 9.99±1.202 (A) | 11.40±1.858 (A) |
| Fenchone | 30.07±1.952 (A) | 24.26±1.287 (AB) | 20.63±1.839 (B) | 23.88±3.513 (AB) |
| Linalool | 469.54±35.062 (A) | 517.58±34.540 (A) | 439.07±35.150 (A) | 401.02±58.626 (A) |
| C10H18O-154(93/79/99/121)-1 | 75.56±8.510 (A) | 47.13±8.114 (A) | 49.84±7.220 (A) | 58.29±9.059 (A) |
| Fenchol | 362.91±22.698 (A) | 333.92±19.730 (A) | 283.50±24.888 (A) | 327.69±50.181 (A) |
| Terpinen-4-ol | 21.14±2.020 (A) | 23.66±1.920 (A) | 21.52±3.533 (A) | 22.98±5.076 (A) |
| Borneol | 128.65±4.992 (A) | 130.24±5.576 (A) | 127.87±8.275 (A) | 127.92±12.006 (A) |
| α-Terpineol | 200.61±11.248 (A) | 204.36±17.180 (A) | 195.94±16.384 (A) | 184.61±25.649 (A) |
| Geraniol | 12.05±6.810 (A) | 5.85±0.541 (A) | 3.99±0.536 (A) | 10.42±2.678 (A) |
| Bornyl acetate | 11.39±0.888 (A) | 13.24±0.757 (A) | 16.16±3.291 (A) | 22.49±4.898 (A) |
| α-Cubebene | 4.09±0.604 (A) | 5.01±0.556 (A) | 5.59±0.826 (A) | 7.52±1.268 (A) |
| Ylangene | 10.28±1.090 (A) | 11.91±1.032 (A) | 13.12±1.419 (A) | 15.62±2.217 (A) |
| α-Copaene | 7.03±0.736 (A) | 7.95±0.929 (A) | 9.11±0.993 (A) | 10.63±1.498 (A) |
| C15H24-204(105/(120+119)/161) | 4.45±0.696 (A) | 5.50±0.654 (A) | 6.11±0.757 (A) | 7.66±1.060 (A) |
| 7-epi-Sesquithujene | 19.42±3.930 (A) | 13.49±1.345 (A) | 20.74±3.752 (A) | 20.65±3.179 (A) |
| β-Isocomene | 24.98±2.458 (A) | 24.40±2.576 (A) | 25.78±3.672 (A) | 23.65±1.260 (A) |
| Sesquithujene | 81.79±9.845 (A) | 98.05±9.369 (A) | 77.24±13.713 (A) | 64.42±8.729 (A) |
| cis-α-Bergamotene | 67.23±10.613 (A) | 57.52±3.297 (A) | 55.96±3.404 (A) | 59.44±3.339 (A) |
| C15H24-204(105/91/147/189) | 17.71±1.129 (A) | 19.02±1.684 (A) | 20.59±1.586 (A) | 23.18±2.138 (A) |

**Table. S16** continued on the next page

**Table. S16** continued from the previous page

| Terpenoid | Control | Mild drought | Moderate drought | Severe drought |
|---|---|---|---|---|
| α-Santalene | 41.08±2.138 (A) | 39.98±2.105 (A) | 42.39±2.613 (A) | 45.45±2.493 (A) |
| β-Caryophyllene | 1386.14±164.244 (A) | 1660.17±151.076 (A) | 1804.11±112.220 (A) | 1873.58±148.571 (A) |
| C15H24-204(120/91/79/69) | 4.53±0.557 (A) | 5.19±0.463 (A) | 5.52±0.418 (A) | 5.55±0.549 (A) |
| γ-Elemene | 86.37±15.246 (A) | 46.08±4.241 (B) | 58.80±9.745 (AB) | 47.99±4.612 (AB) |
| trans-α-Bergamotene | 791.88±53.249 (A) | 779.20±82.427 (A) | 901.04±102.549 (A) | 1079.59±108.278 (A) |
| α-Guaiene | 885.91±84.465 (A) | 939.53±151.114 (A) | 1059.30±123.085 (A) | 1152.55±117.317 (A) |
| Guaia-6,9-diene | 11.08±0.535 (A) | 11.61±1.175 (A) | 12.53±1.325 (A) | 13.05±0.946 (A) |
| C15H24-204(69/91/105/161) | 5.88±0.334 (A) | 5.67±0.576 (A) | 5.99±0.787 (A) | 6.48±0.261 (A) |
| β-Santalene | 16.18±1.363 (A) | 15.12±1.482 (A) | 16.19±1.726 (A) | 17.81±0.992 (A) |
| α-Himachalene | 5.16±0.430 (A) | 5.83±0.483 (A) | 6.32±0.644 (A) | 6.26±0.536 (A) |
| C15H24-204(105/91/133/161/189)-1 | 12.19±1.032 (A) | 13.52±1.335 (A) | 14.46±1.658 (A) | 15.77±1.170 (A) |
| α-Humulene | 712.47±85.629 (A) | 876.58±132.582 (A) | 862.97±95.715 (A) | 917.19±67.501 (A) |
| trans-β-Farnesene | 146.72±13.054 (A) | 124.73±11.376 (A) | 129.85±13.930 (A) | 129.53±4.949 (A) |
| C15H24-204(105)-1 | 40.99±3.384 (A) | 42.78±4.351 (A) | 48.91±4.356 (A) | 49.32±3.604 (A) |
| C15H24-204(133/189)-1 | 55.67±4.236 (A) | 60.91±5.470 (A) | 69.93±6.290 (A) | 67.97±4.887 (A) |
| γ-Muurelene | 76.39±5.929 (A) | 77.75±7.854 (A) | 86.50±9.996 (A) | 92.53±4.430 (A) |
| α-Amorphene | 47.26±3.633 (A) | 50.95±4.326 (A) | 56.41±6.022 (A) | 57.61±2.303 (A) |
| γ-Curcumene | 20.49±1.033 (A) | 20.83±0.999 (A) | 20.85±2.192 (A) | 21.57±0.506 (A) |
| C15H24-204(189/133)-2 | 95.49±7.901 (A) | 103.90±9.885 (A) | 120.37±11.151 (A) | 118.86±8.017 (A) |
| 4,5-di-epi-aristolochene | 22.36±1.051 (A) | 18.37±1.296 (A) | 18.88±2.272 (A) | 20.17±2.723 (A) |
| Sesquisabinene | 21.12±1.884 (A) | 25.81±2.672 (A) | 21.06±1.885 (A) | 20.09±2.948 (A) |
| α-Selinene | 102.76±8.360 (A) | 107.52±9.402 (A) | 123.43±10.727 (A) | 118.40±7.050 (A) |
| α-Curcumene | 9.42±0.676 (A) | 9.12±0.491 (A) | 9.68±0.919 (A) | 9.75±0.480 (A) |
| Valencene | 46.03±2.866 (A) | 46.81±3.738 (A) | 50.27±5.069 (A) | 50.75±1.621 (A) |
| β-Selinene | 454.55±39.853 (A) | 459.30±47.608 (A) | 551.26±84.646 (A) | 574.13±52.249 (A) |
| α-Zingiberene+C15H24-204(105) | 32.15±2.002 (A) | 30.95±1.933 (A) | 30.46±3.348 (A) | 31.30±0.418 (A) |
| δ-Guaiene | 1152.12±95.127 (A) | 1191.96±127.848 (A) | 1472.56±166.722 (A) | 1459.16±106.067 (A) |
| β-Bisabolene | 28.24±1.446 (A) | 27.05±0.678 (A) | 25.29±2.657 (A) | 25.85±0.387 (A) |
| C15H24-204(similar to Germarcene B) | 32.98±1.829 (A) | 31.11±2.285 (A) | 31.22±3.596 (A) | 31.78±0.409 (A) |
| α-Farnesene | 162.50±16.881 (A) | 155.03±18.179 (A) | 142.22±22.312 (A) | 140.91±6.602 (A) |
| C15H24-204(161/105)-Eremophilene | 32.87±1.838 (A) | 31.56±2.300 (A) | 31.95±3.664 (A) | 32.83±0.664 (A) |
| δ-Cadinene-C15H24-204(119/161/105/134) | 16.37±0.629 (A) | 17.38±0.480 (A) | 16.42±1.703 (A) | 16.83±0.479 (A) |
| Sesquicineole | 12.26±1.294 (A) | 11.45±1.105 (A) | 12.46±1.532 (A) | 12.14±0.698 (A) |
| C15H24-204(105)-2 | 37.53±2.270 (A) | 35.90±2.763 (A) | 36.39±4.265 (A) | 36.92±0.803 (A) |

**Table. S16** continued on the next page

**Table. S16** continued from the previous page

| Terpenoid | Control | Mild drought | Moderate drought | Severe drought |
|---|---|---|---|---|
| cis-Calamenene-204(159) | 3.45±0.200 (A) | 3.30±0.210 (A) | 3.60±0.378 (A) | 3.73±0.173 (A) |
| β-Sesquiphellandrene | 23.84±1.265 (A) | 23.32±0.862 (A) | 21.21±2.367 (A) | 21.78±0.334 (A) |
| C15H24-204(189/133)-3 | 289.90±101.773 (A) | 189.20±20.331 (A) | 293.68±82.623 (A) | 299.01±93.824 (A) |
| C15H24-204(161/133/105) | 4906.87±316.834 (A) | 4774.57±568.200 (A) | 5419.09±658.803 (A) | 5536.19±239.335 (A) |
| Selina-3,7(11)-diene | 7582.89±481.675 (A) | 7257.63±829.004 (A) | 8265.28±806.416 (A) | 8128.25±349.382 (A) |
| trans-α-Bisabolene | 41.46±3.356 (A) | 37.62±2.182 (A) | 36.06±4.487 (A) | 36.91±1.930 (A) |
| C15H24-202(202/131/145/159) | 4.32±0.111 (A) | 4.62±0.236 (A) | 4.65±0.598 (A) | 5.03±0.234 (A) |
| Cadala-1(10),3,8-triene-204(157/142) | 51.94±3.608 (A) | 52.71±2.470 (A) | 58.91±6.432 (A) | 61.90±4.304 (A) |
| Germacrene B | 39.10±2.235 (A) | 46.52±3.666 (A) | 51.72±8.440 (A) | 62.04±7.854 (A) |
| C15H24-204(189/133)-4 | 48.37±3.933 (A) | 46.40±3.712 (A) | 46.48±6.010 (A) | 46.80±1.491 (A) |
| trans-Nerolidol | 2493.75±168.216 (A) | 2613.28±175.103 (A) | 2913.49±364.888 (A) | 2682.04±272.088 (A) |
| Caryophyllene oxide | 38.18±3.459 (A) | 38.47±1.309 (A) | 36.40±1.959 (A) | 35.71±1.531 (A) |
| C15H22-202(178/163) | 13.98±0.747 (A) | 11.36±1.413 (A) | 13.65±1.034 (A) | 13.65±0.743 (A) |
| C15H22-202(187/202)-1 | 33.84±1.841 (B) | 32.67±2.127 (AB) | 35.58±2.217 (AB) | 35.77±1.733 (A) |
| C15H26O-222(93/69/41) | 9.39±0.869 (A) | 8.39±1.172 (A) | 11.41±0.886 (A) | 11.62±1.074 (A) |
| C15H24 -204(123/81) | 5.08±0.197 (B) | 5.83±0.416 (AB) | 6.63±0.896 (AB) | 8.10±0.632 (A) |
| Humulene oxide II | 12.81±0.757 (A) | 12.25±1.146 (A) | 14.66±0.939 (A) | 14.52±0.808 (A) |
| C15H26O-222(59/161/91)-1 | 35.41±3.280 (A) | 28.02±2.797 (AB) | 26.47±2.066 (AB) | 23.49±1.322 (B) |
| C15H22-202(187/202)-2 | 7.44±0.144 (A) | 8.57±0.576 (A) | 9.54±1.179 (A) | 11.08±0.934 (A) |
| β-Eudesmol | 72.63±5.973 (A) | 69.57±3.936 (A) | 74.62±4.302 (A) | 67.45±2.193 (A) |
| C15H24O-(similar Caryophyllene oxide)-1 | 22.61±2.252 (A) | 20.09±1.455 (A) | 20.67±1.414 (A) | 18.58±0.659 (A) |
| α-Bisabolol | 25.02±2.703 (A) | 30.78±3.559 (A) | 30.10±4.863 (A) | 32.88±1.580 (A) |
| Juniper camphor | 6.14±0.903 (A) | 7.57±0.680 (A) | 8.30±0.671 (A) | 8.05±0.778 (A) |
| Total MONO,% | 0.70±0.061 (A) | 0.70±0.043 (A) | 0.64±0.069 (A) | 0.71±0.085 (A) |
| Total Sesqui,% | 2.27±0.153 (A) | 2.27±0.225 (A) | 2.56±0.245 (A) | 2.57±0.122 (A) |
| Total % | 2.96±0.191 (A) | 2.97±0.224 (A) | 3.20±0.251 (A) | 3.28±0.187 (A) |

**Table S16.** Effects of the drought treatments on terpenoid concentrations in secondary inflorescences of 'Odem' plants (Experiment 1). Terpenoid concentrations (mg/L; ppm) in the secondary inflorescences of 'Odem' plants subjected to 4 different irrigation treatments, as measured at the end Experiment 1. Samples were taken from the secondary inflorescences. Values are means ± SE. Different letters represent significant differences between irrigation treatments, according to one-way ANOVA and Tukey's HSD test ($P < 0.05$, $5 \leq N \leq 6$).

| Terpenoid | Control | Mild drought | Moderate drought | Severe drought |
|---|---|---|---|---|
| α-Pinene | 1597.088±256.014 (A) | 1363.783±280.698 (A) | 1342.196±73.837 (A) | 1781.305±211.387 (A) |
| Camphene | 168.196±26.347 (A) | 145.050±27.638 (A) | 129.973±9.657 (A) | 154.481±25.207 (A) |
| β-Pinene | 608.752±99.145 (A) | 453.702±76.897 (A) | 483.100±31.729 (A) | 647.545±125.351 (A) |
| β-Myrcene | 1585.400±171.551 (A) | 1143.746±141.054 (AB) | 1017.559±88.420 (B) | 1004.222±136.558 (B) |
| α-Phellandrene | 12.768±2.310 (A) | 11.125±1.330 (A) | 8.704±0.784 (A) | 9.752±0.967 (A) |
| α-Terpinene | 57.253±6.006 (A) | 54.744±4.531 (AB) | 37.255±3.898 (B) | 53.129±4.392 (AB) |
| Limonene | 1077.084±169.728 (A) | 847.150±125.150 (A) | 671.781±53.727 (A) | 826.419±113.099 (A) |
| β-Phellandrene | 58.466±9.286 (A) | 44.197±5.974 (AB) | 30.757±3.946 (B) | 41.911±5.082 (AB) |
| cis-Ocimene | 13.198±1.163 (A) | 12.097±1.320 (A) | 9.068±1.009 (A) | 13.212±2.413 (A) |
| Eucalyptol | 10.390±0.898 (A) | 10.612±1.798 (A) | 6.553±0.508 (A) | 16.143±5.717 (A) |
| trans-Ocimene | 971.224±139.602 (A) | 823.439±91.598 (A) | 746.362±71.474 (A) | 822.977±127.011 (A) |
| γ-Terpinene | 40.686±2.877 (A) | 38.227±2.719 (A) | 29.162±2.668 (A) | 39.117±4.295 (A) |
| Terpinolene | 51.917±5.803 (A) | 47.809±4.098 (A) | 37.284±3.871 (A) | 46.092±2.999 (A) |
| cis-Linalool oxide | 93.817±15.188 (A) | 88.994±16.921 (A) | 75.184±10.791 (A) | 84.678±14.836 (A) |
| trans-Linalool oxide | 7.440±1.018 (A) | 7.123±0.931 (A) | 5.173±0.666 (A) | 5.682±0.473 (A) |
| Fenchone | 16.280±2.076 (A) | 15.294±1.074 (A) | 11.629±1.120 (A) | 12.451±0.851 (A) |
| Linalool | 458.699±62.894 (A) | 460.614±15.430 (A) | 354.674±34.252 (A) | 357.755±27.333 (A) |
| C10H18O-154(93/79/99/121)-1 | 24.158±3.843 (A) | 24.099±4.414 (A) | 16.268±2.103 (A) | 19.214±2.733 (A) |
| Fenchol | 458.446±57.217 (A) | 436.412±21.210 (AB) | 311.547±31.919 (B) | 359.038±27.106 (AB) |
| Terpinen-4-ol | 18.639±2.281 (AB) | 19.891±1.088 (A) | 12.551±1.660 (B) | 16.735±1.121 (AB) |
| Borneol | 159.692±8.151 (A) | 154.707±4.672 (A) | 147.655±3.260 (A) | 136.877±7.764 (A) |
| α-Terpineol | 204.594±16.630 (A) | 203.219±6.978 (A) | 174.174±7.292 (A) | 167.989±11.417 (A) |
| Geraniol | 26.292±3.299 (A) | 31.658±6.006 (A) | 16.667±3.123 (A) | 25.445±9.560 (A) |
| Bornyl acetate | 25.794±6.288 (A) | 25.076±6.861 (A) | 17.591±8.360 (A) | 20.734±6.109 (A) |
| β-Isocomene | 7.654±0.794 (A) | 6.175±0.530 (A) | 7.525±0.844 (A) | 7.148±1.048 (A) |
| Sesquithujene | 19.397±3.184 (A) | 17.421±1.463 (A) | 21.688±3.688 (A) | 21.895±3.691 (A) |
| cis-α-Bergamotene | 37.333±3.708 (A) | 33.143±3.409 (A) | 37.467±4.014 (A) | 37.004±4.907 (A) |
| α-Santalene | 22.239±0.652 (A) | 20.962±0.823 (A) | 22.484±1.151 (A) | 20.439±0.804 (A) |
| β-Caryophyllene | 359.398±28.456 (A) | 319.991±18.909 (A) | 403.787±47.411 (A) | 359.627±47.427 (A) |
| γ-Elemene | 11.083±2.876 (A) | 8.184±1.109 (A) | 13.693±2.325 (A) | 13.271±2.582 (A) |
| trans-α-Bergamotene | 143.413±11.318 (A) | 137.368±9.742 (A) | 150.657±16.950 (A) | 145.290±17.795 (A) |
| α-Guaiene | 216.391±12.354 (A) | 207.232±16.141 (A) | 244.292±31.622 (A) | 216.747±30.927 (A) |
| β-Santalene | 4.186±0.390 (A) | 4.013±0.378 (A) | 4.603±0.671 (A) | 4.228±0.708 (A) |
| α-Humulene | 142.546±11.254 (A) | 131.377±7.458 (A) | 147.941±12.329 (A) | 154.588±24.718 (A) |

**Table. S17** continued on the next page

**Table. S17** continued from the previous page

| Terpenoid | Control | Mild drought | Moderate drought | Severe drought |
|---|---|---|---|---|
| trans-β-Farnesene | 55.397±4.082 (A) | 55.383±3.251 (A) | 58.471±5.477 (A) | 51.627±3.872 (A) |
| C15H24-204(105)-1 | 17.437±0.391 (AB) | 16.981±0.679 (AB) | 18.398±0.710 (A) | 15.994±0.396 (B) |
| C15H24-204(133/189)-1 | 75.240±5.933 (A) | 78.433±7.252 (A) | 76.789±8.975 (A) | 54.916±5.888 (A) |
| γ-Muurelene | 19.725±0.768 (A) | 18.842±0.709 (A) | 20.528±1.445 (A) | 17.928±0.703 (A) |
| α-Amorphene | 19.218±0.639 (A) | 18.238±0.649 (A) | 19.708±1.003 (A) | 17.094±0.533 (A) |
| γ-Curcumene | 19.617±1.430 (A) | 18.598±1.187 (A) | 21.308±1.959 (A) | 17.995±1.860 (A) |
| C15H24-204(189/133)-2 | 71.551±7.895 (A) | 66.375±6.746 (A) | 74.774±12.073 (A) | 75.228±13.794 (A) |
| 4,5-di-epi-aristolochene | 12.386±1.330 (A) | 12.962±1.363 (A) | 15.452±1.231 (A) | 10.685±1.078 (A) |
| Sesquisabinene | 12.294±1.470 (A) | 12.013±1.042 (A) | 14.051±2.380 (A) | 12.699±2.205 (A) |
| α-Selinene | 49.791±2.944 (A) | 50.984±3.130 (A) | 53.138±7.016 (A) | 48.393±4.626 (A) |
| α-Curcumene | 2.905±0.345 (A) | 3.145±0.193 (A) | 3.268±0.470 (A) | 2.752±0.406 (A) |
| Valencene | 24.690±1.215 (A) | 23.775±0.903 (A) | 27.080±2.137 (A) | 22.829±1.125 (A) |
| β-Selinene | 108.529±7.498 (A) | 110.820±7.416 (A) | 114.221±7.803 (A) | 105.171±12.052 (A) |
| α-Zingiberene+C15H24-204(105) | 21.965±1.149 (A) | 21.537±0.757 (A) | 22.721±1.568 (A) | 19.638±0.793 (A) |
| δ-Guaiene | 307.778±23.826 (A) | 303.283±20.807 (A) | 337.916±24.619 (A) | 285.309±37.478 (A) |
| β-Bisabolene | 19.988±1.204 (A) | 20.588±0.953 (A) | 21.553±2.229 (A) | 19.066±2.544 (A) |
| C15H24-204(similar to Germarcene B) | 21.669±1.440 (A) | 20.876±0.974 (A) | 21.562±1.437 (A) | 18.925±0.938 (A) |
| α-Farnesene | 79.659±14.295 (A) | 78.583±8.679 (A) | 72.661±10.035 (A) | 65.555±11.097 (A) |
| C15H24-204(161/105)-Eremophilene | 21.765±1.281 (A) | 20.564±0.868 (A) | 21.434±1.941 (A) | 18.813±0.839 (A) |
| δ-Cadinene-C15H24-204(119/161/105/134) | 19.266±1.112 (A) | 18.497±0.895 (A) | 18.397±1.612 (A) | 14.605±1.005 (A) |
| Sesquicineole | 3.678±0.353 (A) | 3.867±0.182 (A) | 4.367±0.701 (A) | 4.000±0.603 (A) |
| C15H24-204(105)-2 | 24.380±1.195 (A) | 23.709±1.037 (A) | 24.621±1.827 (A) | 21.327±0.833 (A) |
| β-Sesquiphellandrene | 23.260±1.581 (A) | 25.704±1.079 (A) | 26.381±2.357 (A) | 19.691±1.573 (A) |
| C15H24-204(189/133)-3 | 90.551±8.576 (A) | 90.994±6.535 (A) | 98.704±11.909 (A) | 82.247±8.899 (A) |
| C15H24-204(161/133/105) | 1241.87±132.167 (A) | 1304.95±123.461 (A) | 1416.51±166.773(A) | 1118.45±141.532 (A) |
| Selina-3,7(11)-diene | 2122.28±248.269 (A) | 2127.53±176.748 (A) | 1989.58±190.87 (A) | 1901.95±280.153 (A) |
| trans-α-Bisabolene | 25.449±2.271 (A) | 23.552±1.204 (A) | 21.938±2.816 (A) | 20.825±1.666 (A) |
| Cadala-1(10),3,8-triene-204(157/142) | 31.192±2.250 (A) | 32.075±1.557 (A) | 31.256±3.857 (A) | 28.943±2.700 (A) |
| Germacrene B | 22.067±6.206 (A) | 15.213±2.161 (A) | 25.615±7.025 (A) | 26.577±6.981 (A) |
| C15H24-204(189/133)-4 | 72.275±10.346 (A) | 64.425±4.181 (A) | 51.704±6.732 (A) | 62.357±9.589 (A) |
| trans-Nerolidol | 463.000±43.669 (A) | 546.978±37.610 (A) | 480.397±110.37 (A) | 439.661±67.597 (A) |
| Caryophyllene oxide | 26.284±1.517 (A) | 25.665±1.062 (A) | 23.808±3.443 (A) | 21.398±2.007 (A) |
| C15H22-202(178/163) | 5.666±0.738 (A) | 6.274±0.454 (A) | 4.650±1.010 (A) | 4.867±0.646 (A) |
| C15H22-202(187/202)-1 | 26.284±1.517 (A) | 25.665±1.062 (A) | 23.808±3.443 (A) | 21.398±2.007 (A) |

**Table. S17** continued on the next page

**Table. S17** continued from the previous page

| Terpenoid | Control | Mild drought | Moderate drought | Severe drought |
|---|---|---|---|---|
| C15H26O-222(93/69/41) | 18.310±3.309 (A) | 19.884±2.341 (A) | 18.845±3.349 (A) | 17.943±2.156 (A) |
| C15H24 -204(123/81) | 4.895±0.799 (A) | 4.296±0.339 (A) | 3.803±0.570 (A) | 4.463±0.739 (A) |
| Humulene oxide II | 6.149±0.767 (A) | 6.080±0.295 (A) | 5.473±1.118 (A) | 5.924±0.785 (A) |
| C15H26O-222(59/161/91)-1 | 6.590±0.856 (A) | 7.263±0.672 (A) | 4.532±0.760 (A) | 5.022±0.627 (A) |
| γ-Eudesmol | 24.362±2.290 (A) | 19.825±2.611 (AB) | 13.500±2.647 (B) | 14.886±3.135 (AB) |
| C15H26O-222(105/59/161)-1 | 5.335±0.694 (A) | 4.818±0.532 (A) | 3.512±0.367 (A) | 3.629±0.457 (A) |
| C15H22-202(187/202)-2 | 12.379±1.911 (A) | 12.955±1.205 (A) | 8.044±0.940 (A) | 10.451±1.540 (A) |
| β-Eudesmol | 72.647±2.665 (AB) | 80.312±5.982 (A) | 58.515±4.609 (B) | 61.580±5.156 (AB) |
| α-Bisabolol | 23.010±1.855 (A) | 22.480±1.457 (A) | 20.008±1.955 (A) | 20.542±1.218 (A) |
| Juniper camphor | 18.344±0.717 (A) | 18.047±0.903 (A) | 17.027±1.096 (A) | 15.590±0.895 (A) |
| Total Mono, ppm | 7756.016±886.194 (A) | 6472.551±703.407 (A) | 5700.431±314.665 (A) | 6670.560±682.166 (A) |
| Total Mono, % | 0.776±0.089 (A) | 0.647±0.070 (A) | 0.570±0.031 (A) | 0.667±0.068 (A) |
| Total Sesqui, ppm | 6357.773±570.536 (A) | 6411.498±423.005 (A) | 6505.522±657.991 (A) | 5850.335±699.860 (A) |
| Total Sesqui, % | 0.636±0.057 (A) | 0.641±0.042 (A) | 0.651±0.066 (A) | 0.585±0.070 (A) |
| Total Terpenoid (ppm) | 14113.789±1140.158 (A) | 12884.048±1011.678 (A) | 12205.953±763.345 (A) | 12520.895±1096.806 (A) |
| Total Terpenoid % | 1.411±0.114 (A) | 1.288±0.101 (A) | 1.221±0.076 (A) | 1.252±0.110 (A) |

**Table S17.** Effects of the drought treatments on terpenoid concentrations in secondary inflorescences of 'Odem' plants (Experiment 2). Terpenoid concentrations (mg/L; ppm) in the secondary inflorescences of 'Odem' plants subjected to 4 different irrigation treatments, as measured in the middle of physiological Phase III in Experiment 2. Samples were taken from the secondary inflorescences. Values are means ± SE. Different letters represent significant differences between irrigation treatments, according to one-way ANOVA and Tukey's HSD test ($P < 0.05$, $5 \leq N \leq 6$).

| Terpenoid | Control | Mild drought | Moderate drought | Severe drought |
|---|---|---|---|---|
| α-Pinene | 1071.813±244.187 (A) | 1227.718±227.011 (A) | 1056.449±223.859 (A) | 1332.717±320.043 (A) |
| Camphene | 100.904±18.790 (A) | 102.796±19.942 (A) | 82.816±13.974 (A) | 99.682±21.054 (A) |
| β-Pinene | 364.759±76.925 (A) | 368.605±70.859 (A) | 316.768±68.019 (A) | 431.090±96.607 (A) |
| β-Myrcene | 1699.079±387.545 (A) | 1352.927±275.426 (A) | 968.311±155.801 (A) | 1257.570±208.314 (A) |
| α-Phellandrene | 11.214±1.913 (A) | 10.209±1.705 (A) | 7.117±1.014 (A) | 9.780±1.254 (A) |
| α-Terpinene | 44.360±7.066 (A) | 41.547±6.648 (A) | 29.786±3.652 (A) | 39.535±6.217 (A) |
| Limonene | 978.111±238.257 (A) | 860.403±179.104 (A) | 579.739±121.587 (A) | 798.539±166.579 (A) |
| β-Phellandrene | 46.938±11.249 (A) | 39.669±8.335 (A) | 27.344±6.030 (A) | 41.946±8.678 (A) |
| cis-Ocimene | 12.644±3.048 (A) | 11.014±1.920 (A) | 8.166±1.802 (A) | 11.981±2.703 (A) |
| Eucalyptol | 11.681±2.027 (A) | 11.449±1.577 (A) | 7.912±1.512 (A) | 10.429±1.969 (A) |
| trans-Ocimene | 946.956±274.514 (A) | 825.370±183.616 (A) | 617.138±147.501 (A) | 902.186±270.749 (A) |
| γ-Terpinene | 32.671±5.790 (A) | 28.764±4.427 (A) | 22.096±2.851 (A) | 28.957±4.621 (A) |
| Terpinolene | 40.368±7.625 (A) | 35.158±5.367 (A) | 26.589±3.535 (A) | 36.041±5.423 (A) |
| cis-Linalool oxide | 118.851±18.795 (A) | 88.307±17.039 (A) | 65.243±9.301 (A) | 67.042±12.778 (A) |
| trans-Linalool oxide | 8.186±1.461 (A) | 6.406±1.142 (A) | 4.446±0.602 (A) | 4.904±0.812 (A) |
| Fenchone | 15.617±2.510 (A) | 12.513±2.008 (A) | 9.457±0.820 (A) | 11.317±1.654 (A) |
| Linalool | 358.234±45.837 (A) | 326.420±41.782 (A) | 256.829±19.144 (A) | 270.227±28.167 (A) |
| C10H18O-154(93/79/99/121)-1 | 24.628±4.810 (A) | 18.649±4.088 (A) | 13.225±1.269 (A) | 13.515±3.110 (A) |
| Fenchol | 363.489±57.930 (A) | 315.499±52.235 (A) | 223.491±20.491 (A) | 249.552±39.267 (A) |
| Terpinen-4-ol | 14.950±2.064 (A) | 13.286±1.807 (A) | 10.050±0.980 (A) | 12.407±1.436 (A) |
| Borneol | 140.951±9.139 (A) | 139.240±5.200 (A) | 131.733±2.963 (A) | 138.130±3.233 (A) |
| α-Terpineol | 180.492±13.464 (A) | 173.560±10.949 (A) | 155.625±4.703 (A) | 161.110±5.234 (A) |
| Geraniol | 25.886±5.406 (A) | 17.875±2.496 (AB) | 12.551±1.077 (B) | 14.054±2.014 (AB) |
| Bornyl acetate | 16.486±7.953 (A) | 10.379±4.200 (A) | 6.841±3.432 (A) | 16.628±7.103 (A) |
| β-Isocomene | 9.075±1.704 (A) | 7.968±1.653 (A) | 8.157±1.313 (A) | 10.729±2.257 (A) |
| Sesquithujene | 27.756±4.920 (A) | 25.022±5.165 (A) | 27.008±4.336 (A) | 35.427±7.788 (A) |
| cis-α-Bergamotene | 44.727±4.790 (A) | 41.990±7.198 (A) | 44.164±3.734 (A) | 44.582±7.670 (A) |
| α-Santalene | 25.003±2.531 (A) | 22.173±2.525 (A) | 23.918±1.490 (A) | 25.327±3.799 (A) |
| β-Caryophyllene | 438.177±73.047 (A) | 433.374±66.344 (A) | 450.769±63.887 (A) | 541.490±103.968 (A) |
| trans-α-Bergamotene | 206.945±33.676 (A) | 178.695±26.908 (A) | 185.145±22.136 (A) | 220.280±44.746 (A) |
| α-Guaiene | 265.428±43.788 (A) | 254.965±42.848 (A) | 264.269±46.262 (A) | 335.332±89.533 (A) |
| Guaia-6,9-diene | 4.175±0.630 (A) | 4.053±0.871 (A) | 4.210±0.620 (A) | 6.573±1.073 (A) |
| β-Santalene | 5.851±0.981 (A) | 5.297±0.867 (A) | 5.557±0.760 (A) | 6.773±1.281 (A) |
| α-Humulene | 179.864±28.754 (A) | 183.281±27.186 (A) | 199.415±30.574 (A) | 227.406±53.853 (A) |

**Table. S18** continued on the next page

**Table. S18** continued from the previous page

| Terpenoid | Control | Mild drought | Moderate drought | Severe drought |
|---|---|---|---|---|
| trans-β-Farnesene | 77.610±10.266 (A) | 67.669±8.644 (A) | 69.300±6.665 (A) | 72.256±10.895 (A) |
| C15H24-204(105)-1 | 17.789±1.322 (A) | 16.743±1.794 (A) | 18.354±0.940 (A) | 17.556±2.452 (A) |
| C15H24-204(133/189)-1 | 64.087±11.062 (A) | 65.501±10.129 (A) | 84.402±6.360 (A) | 66.385±12.263 (A) |
| γ-Muurelene | 21.529±2.126 (A) | 21.144±1.902 (A) | 21.861±1.504 (A) | 22.183±2.409 (A) |
| α-Amorphene | 18.841±1.350 (A) | 18.329±1.465 (A) | 19.925±0.926 (A) | 19.473±1.801 (A) |
| γ-Curcumene | 16.497±1.140 (A) | 16.926±0.198 (A) | 16.543±0.666 (A) | 17.457±0.648 (A) |
| C15H24-204(189/133)-2 | 62.563±6.386 (A) | 70.750±3.340 (A) | 59.963±5.898 (A) | 79.974±8.017 (A) |
| 4,5-di-epi-aristolochene | 13.945±1.939 (A) | 13.162±2.249 (A) | 14.281±1.091 (A) | 14.357±2.064 (A) |
| Sesquisabinene | 18.694±2.976 (A) | 15.796±2.636 (A) | 16.624±2.066 (A) | 17.757±3.770 (A) |
| α-Selinene | 59.528±7.201 (A) | 58.475±7.373 (A) | 61.170±6.185 (A) | 64.428±8.525 (A) |
| α-Curcumene | 5.465±0.968 (A) | 4.660±0.667 (A) | 4.754±0.443 (A) | 4.696±0.697 (A) |
| Valencene | 27.399±2.716 (A) | 26.981±2.459 (A) | 27.743±1.924 (A) | 28.942±2.814 (A) |
| β-Selinene | 160.512±31.551 (A) | 156.414±7.545 (A) | 148.875±23.569 (A) | 167.481±27.853 (A) |
| α-Zingiberene+C15H24-204(105) | 24.168±2.073 (A) | 23.244±1.437 (A) | 23.526±0.984 (A) | 23.714±1.550 (A) |
| δ-Guaiene | 386.265±64.959 (A) | 382.098±49.254 (A) | 354.035±32.599 (A) | 391.955±45.161 (A) |
| β-Bisabolene | 24.542±2.800 (A) | 23.446±3.036 (A) | 25.940±1.074 (A) | 22.066±4.061 (A) |
| C15H24-204(similar to Germarcene B) | 24.381±2.414 (A) | 23.196±1.792 (A) | 23.058±1.038 (A) | 23.276±1.826 (A) |
| α-Farnesene | 87.549±15.322 (A) | 76.673±9.417 (A) | 67.266±4.834 (A) | 62.267±10.119 (A) |
| C15H24-204(161/105)-Eremophilene | 24.583±2.435 (A) | 23.451±1.814 (A) | 23.486±1.389 (A) | 23.399±1.794 (A) |
| δ-Cadinene-C15H24-204(119/161/105/134) | 18.588±1.224 (A) | 15.899±0.525 (A) | 20.459±1.985 (A) | 16.789±1.796 (A) |
| Sesquicineole | 6.078±0.663 (A) | 5.682±0.573 (A) | 6.051±0.462 (A) | 6.330±1.042 (A) |
| C15H24-204(105)-2 | 26.682±2.467 (A) | 26.129±1.576 (A) | 25.880±1.355 (A) | 25.795±1.843 (A) |
| β-Sesquiphellandrene | 24.145±1.579 (A) | 23.271±0.894 (A) | 25.775±0.637 (A) | 22.850±3.290 (A) |
| C15H24-204(189/133)-3 | 107.190±12.381 (A) | 102.555±10.421 (A) | 102.472±7.458 (A) | 103.924±12.614 (A) |
| C15H24-204(161/133/105) | 1588.790±220.30 (A) | 1494.599±173.99 (A) | 1459.36±145.058 (A) | 1562.345±112.96 (A) |
| Selina-3,7(11)-diene | 2813.480±416.37 (A) | 2672.527±272.05 (A) | 2413.602±164.52 (A) | 2559.762±241.85 (A) |
| trans-α-Bisabolene | 26.567±3.176 (A) | 25.778±1.091 (A) | 24.575±1.834 (A) | 23.269±1.743 (A) |
| Cadala-1(10),3,8-triene-204(157/142) | 43.052±4.382 (A) | 40.408±4.046 (A) | 45.401±2.982 (A) | 41.930±4.456 (A) |
| Germacrene B | 5.007±1.086 (A) | 5.217±1.052 (A) | 7.113±1.509 (A) | 10.491±2.278 (A) |
| C15H24-204(189/133)-4 | 76.380±10.984 (A) | 68.853±10.748 (A) | 76.377±8.081 (A) | 68.988±11.767 (A) |
| trans-Nerolidol | 426.926±51.507 (A) | 396.498±71.112 (A) | 421.369±35.420 (A) | 404.115±46.335 (A) |
| Caryophyllene oxide | 24.972±2.063 (A) | 23.752±1.959 (A) | 26.141±0.713 (A) | 23.784±2.536 (A) |
| C15H22-202(178/163) | 9.211±1.329 (A) | 7.849±1.149 (A) | 8.581±0.650 (A) | 7.012±0.965 (A) |
| C15H22-202(187/202)-1 | 24.972±2.063 (A) | 23.752±1.959 (A) | 26.141±0.713 (A) | 23.784±2.536 (A) |

**Table. S18** continued on the next page

**Table. S18** continued from the previous page

| Terpenoid | Control | Mild drought | Moderate drought | Severe drought |
|---|---|---|---|---|
| C15H26O-222(93/69/41) | 28.094±1.759 (A) | 26.110±2.413 (A) | 28.204±1.685 (A) | 26.525±3.793 (A) |
| C15H24 -204(123/81) | 4.987±0.766 (A) | 4.424±0.593 (A) | 4.892±0.408 (A) | 5.131±0.811 (A) |
| Humulene oxide II | 8.008±0.973 (A) | 7.485±0.882 (A) | 8.491±0.830 (A) | 7.571±1.264 (A) |
| C15H26O-222(59/161/91)-1 | 5.531±0.536 (A) | 5.160±0.485 (A) | 4.711±0.294 (A) | 3.976±0.565 (A) |
| γ-Eudesmol | 23.532±2.069 (A) | 20.212±2.371 (A) | 18.480±2.088 (A) | 16.150±2.512 (A) |
| C15H22-202(187/202)-2 | 9.358±1.513 (A) | 8.482±1.080 (A) | 8.116±0.676 (A) | 7.676±1.116 (A) |
| β-Eudesmol | 58.913±5.532 (A) | 55.815±5.285 (A) | 50.959±3.826 (A) | 49.362±5.747 (A) |
| α-Bisabolol | 19.803±1.603 (A) | 18.668±0.807 (A) | 18.491±0.865 (A) | 18.887±1.431 (A) |
| Juniper camphor | 13.557±1.001 (A) | 13.395±0.296 (A) | 13.556±0.424 (A) | 13.870±0.423 (A) |
| Total Mono, ppm | 6629.265±1397.575 (A) | 6037.764±1092.102 (A) | 4639.721±755.501 (A) | 5959.338±1178.843 (A) |
| Total Mono, % | 0.663±0.140 (A) | 0.604±0.109 (A) | 0.464±0.076 (A) | 0.596±0.118 (A) |
| Total Sesqui, ppm | 7757.727±1046.775 (A) | 7373.721±802.145 (A) | 7158.740±618.723 (A) | 7670.294±866.327 (A) |
| Total Sesqui, % | 0.776±0.105 (A) | 0.737±0.080 (A) | 0.716±0.062 (A) | 0.767±0.087 (A) |
| Toal Terpenoid (ppm) | 14386.992±2404.922 (A) | 13411.484±1880.280 (A) | 11798.461±1337.625 (A) | 13629.633±1968.497 (A) |
| Total Terpenoid (%) | 1.439±0.240 (A) | 1.341±0.188 (A) | 1.180±0.134 (A) | 1.363±0.197 (A) |

**Table S18.** Effects of the drought treatment on terpenoid concentrations in primary inflorescences of ‘Odem’ plants (Experiment 2). Terpenoid concentrations (mg/L; ppm) in the primary inflorescences of ‘Odem’ plants subjected to 4 different irrigation treatments, as measured at the end of Experiment 2. Samples were taken from the primary inflorescences. Values are means ± SE. Different letters represent significant differences between irrigation treatments, according to one-way ANOVA and Tukey’s HSD test ($P < 0.05$, $5 \leq N \leq 6$).

| Terpenoid | Control | Mild drought | Moderate drought | Severe drought |
|---|---|---|---|---|
| α-Pinene | 1113.20±364.582 (A) | 1173.09±253.175 (A) | 974.19±178.404 (A) | 1294.78±211.434 (A) |
| Camphene | 109.08±23.674 (A) | 101.14±17.449 (A) | 81.48±12.275 (A) | 103.26±15.788 (A) |
| β-Pinene | 354.19±81.306 (A) | 340.32±51.275 (A) | 312.60±50.488 (A) | 395.67±68.613 (A) |
| β-Myrcene | 1572.85±346.372 (A) | 1044.21±104.871 (A) | 1007.62±154.680 (A) | 1012.42±181.626 (A) |
| α-Phellandrene | 12.25±2.388 (A) | 8.37±0.710 (A) | 7.21±0.843 (A) | 7.64±1.363 (A) |
| α-Terpinene | 46.73±7.139 (A) | 33.90±2.401 (A) | 29.78±3.134 (A) | 32.70±5.676 (A) |
| Limonene | 948.94±230.559 (A) | 660.41±82.126 (A) | 600.15±97.540 (A) | 706.25±140.114 (A) |
| β-Phellandrene | 46.76±11.491 (A) | 29.58±3.141 (A) | 27.70±4.626 (A) | 30.13±6.130 (A) |
| cis-Ocimene | 12.07±3.425 (A) | 8.34±1.101 (A) | 8.11±1.389 (A) | 9.27±2.017 (A) |
| Eucalyptol | 10.85±2.593 (A) | 7.69±1.103 (A) | 7.77±1.534 (A) | 9.35±1.664 (A) |
| trans-Ocimene | 852.75±220.133 (A) | 626.75±105.533 (A) | 632.42±114.332 (A) | 710.21±207.153 (A) |
| γ-Terpinene | 32.28±5.004 (A) | 23.92±1.759 (A) | 21.93±2.299 (A) | 24.56±3.296 (A) |
| Terpinolene | 39.92±6.587 (A) | 30.47±2.379 (A) | 26.94±2.704 (A) | 30.42±4.351 (A) |
| cis-Linalool oxide | 115.93±9.217 (A) | 70.43±11.193 (B) | 63.82±8.639 (B) | 60.16±12.077 (B) |
| trans-Linalool oxide | 8.27±0.618 (A) | 5.46±0.462 (B) | 4.94±0.348 (B) | 4.42±0.785 (B) |
| Fenchone | 15.69±1.725 (A) | 11.53±0.860 (AB) | 9.85±0.772 (B) | 10.20±1.505 (B) |
| Linalool | 369.10±34.288 (A) | 301.03±17.812 (AB) | 262.61±17.978 (B) | 244.43±24.164 (B) |
| C10H18O-154(93/79/99/121)-1 | 21.86±2.892 (A) | 14.36±1.431 (B) | 12.42±1.510 (B) | 12.87±1.458 (B) |
| Fenchol | 375.05±37.463 (A) | 285.36±20.753 (AB) | 228.82±20.218 (B) | 238.89±30.068 (B) |
| Terpinen-4-ol | 15.35±1.331 (A) | 11.74±1.061 (AB) | 9.87±1.093 (B) | 10.18±1.296 (B) |
| Borneol | 145.26±3.971 (A) | 136.49±2.446 (A) | 124.38±9.729 (A) | 131.24±3.330 (A) |
| α-Terpineol | 188.04±9.804 (A) | 167.76±5.095 (AB) | 156.16±6.491 (B) | 151.48±5.193 (B) |
| Geraniol | 30.44±5.099 (A) | 16.72±1.104 (B) | 13.32±1.260 (B) | 14.67±1.965 (B) |
| Bornyl acetate | 20.38±8.938 (A) | 10.40±3.542 (A) | 4.44±0.727 (A) | 16.78±7.215 (A) |
| β-Isocomene | 9.05±1.704 (A) | 6.44±0.835 (A) | 6.95±0.895 (A) | 9.95±1.751 (A) |
| Sesquithujene | 23.75±3.508 (A) | 18.00±1.980 (A) | 21.19±2.683 (A) | 30.56±5.353 (A) |
| cis-α-Bergamotene | 44.09±3.907 (A) | 35.48±4.057 (A) | 36.33±3.017 (A) | 48.82±3.441 (A) |
| α-Santalene | 24.37±1.892 (A) | 21.67±1.112 (A) | 21.99±1.078 (A) | 25.59±1.630 (A) |
| β-Caryophyllene | 422.44±91.678 (A) | 338.76±49.149 (A) | 375.49±59.262 (A) | 493.15±72.298 (A) |
| trans-α-Bergamotene | 195.92±36.204 (A) | 154.82±20.999 (A) | 154.13±18.818 (A) | 214.08±24.592 (A) |
| α-Guaiene | 306.44±90.744 (A) | 235.81±34.240 (A) | 228.07±26.989 (A) | 321.03±53.257 (A) |
| Guaia-6,9-diene | 3.68±1.983 (A) | 3.49±0.787 (A) | 3.94±0.351 (A) | 4.85±0.799 (A) |
| β-Santalene | 5.65±0.990 (A) | 4.52±0.633 (A) | 4.53±0.566 (A) | 6.34±0.794 (A) |

**Table. S19** continued on the next page

**Table. S19** continued from the previous page

| Terpenoid | Control | Mild drought | Moderate drought | Severe drought |
|---|---|---|---|---|
| α-Humulene | 178.43±37.485 (A) | 148.41±20.853 (A) | 161.78±23.973 (A) | 219.28±33.527 (A) |
| trans-β-Farnesene | 74.23±7.318 (A) | 62.02±4.832 (A) | 61.43±4.308 (A) | 73.19±5.164 (A) |
| C15H24-204(105)-1 | 18.42±1.260 (A) | 17.11±1.153 (A) | 16.82±1.114 (A) | 18.94±0.659 (A) |
| C15H24-204(133/189)-1 | 77.83±8.156 (A) | 74.97±7.994 (A) | 77.13±7.656 (A) | 87.00±10.727 (A) |
| γ-Muurelene | 22.58±2.152 (A) | 20.33±1.175 (A) | 20.37±1.061 (A) | 23.08±1.319 (A) |
| α-Amorphene | 19.65±1.191 (A) | 18.75±0.662 (A) | 18.50±0.634 (A) | 20.17±0.642 (A) |
| γ-Curcumene | 16.20±0.927 (A) | 18.35±0.970 (A) | 17.26±0.953 (A) | 16.92±0.513 (A) |
| C15H24-204(189/133)-2 | 59.44±9.344 (A) | 66.34±9.721 (A) | 64.95±5.585 (A) | 65.57±5.065 (A) |
| 4,5-di-epi-aristolochene | 14.16±1.553 (A) | 13.81±1.224 (A) | 13.26±1.026 (A) | 15.79±1.026 (A) |
| Sesquisabinene | 17.10±2.166 (A) | 13.74±1.557 (A) | 13.78±1.399 (A) | 18.03±1.854 (A) |
| α-Selinene | 62.62±9.409 (A) | 57.12±6.365 (A) | 55.07±4.720 (A) | 66.07±5.019 (A) |
| α-Curcumene | 5.01±0.667 (A) | 4.25±0.549 (A) | 3.93±0.385 (A) | 4.61±0.349 (A) |
| Valencene | 29.02±2.773 (A) | 25.98±1.595 (A) | 25.94±1.395 (A) | 29.05±1.475 (A) |
| β-Selinene | 143.82±25.061 (A) | 135.91±17.207 (A) | 135.07±16.270 (A) | 154.23±11.904 (A) |
| α-Zingiberene+C15H24-204(105) | 24.61±1.215 (A) | 22.74±0.885 (A) | 22.14±0.785 (A) | 23.86±0.811 (A) |
| δ-Guaiene | 393.89±71.261 (A) | 361.26±41.635 (A) | 340.71±32.585 (A) | 405.97±35.902 (A) |
| β-Bisabolene | 25.28±1.769 (A) | 24.86±1.496 (A) | 25.12±2.027 (A) | 25.28±2.102 (A) |
| C15H24-204(similar to Germarcene B) | 24.80±1.492 (A) | 22.27±0.889 (A) | 21.96±0.922 (A) | 23.52±0.971 (A) |
| α-Farnesene | 92.25±7.837 (A) | 73.63±4.586 (AB) | 69.25±4.864 (B) | 75.63±5.145 (AB) |
| C15H24-204(161/105)-Eremophilene | 25.00±1.809 (A) | 22.28±1.119 (A) | 21.73±0.948 (A) | 23.58±1.166 (A) |
| δ-Cadinene-C15H24-204(119/161/105/134) | 20.75±1.850 (A) | 20.16±1.605 (A) | 18.98±2.178 (A) | 19.19±0.805 (A) |
| Sesquicineole | 5.68±0.612 (A) | 5.06±0.442 (A) | 5.06±0.371 (A) | 5.98±0.396 (A) |
| C15H24-204(105)-2 | 27.24±1.714 (A) | 24.69±1.050 (A) | 24.23±1.022 (A) | 26.17±1.076 (A) |
| β-Sesquiphellandrene | 25.97±1.227 (A) | 25.09±0.909 (A) | 25.95±1.896 (A) | 27.73±0.864 (A) |
| C15H24-204(189/133)-3 | 111.01±10.744 (A) | 94.19±5.832 (A) | 95.82±6.018 (A) | 106.54±5.923 (A) |
| C15H24-204(161/133/105) | 1675.87±204.077 (A) | 1390.43±115.483 (A) | 1351.86±94.257 (A) | 1539.88±118.635 (A) |
| Selina-3,7(11)-diene | 2913.18±351.829 (A) | 2460.58±203.930 (A) | 2287.18±179.679 (A) | 2720.63±188.667 (A) |
| trans-α-Bisabolene | 25.50±1.296 (A) | 22.82±1.395 (A) | 24.98±1.303 (A) | 24.00±0.880 (A) |
| Cadala-1(10),3,8-triene-204(157/142) | 46.52±3.435 (A) | 44.80±4.143 (A) | 41.17±2.532 (A) | 46.36±1.508 (A) |
| Germacrene B | 5.43±1.020 (B) | 4.82±0.739 (B) | 5.95±0.868 (B) | 10.39±1.325 (A) |
| C15H24-204(189/133)-4 | 81.15±7.522 (A) | 70.43±6.690 (A) | 67.68±6.001 (A) | 78.33±6.104 (A) |
| trans-Nerolidol | 486.48±31.268 (A) | 437.91±42.134 (A) | 407.06±44.690 (A) | 402.55±43.860 (A) |
| Caryophyllene oxide | 26.21±0.705 (A) | 25.51±0.990 (A) | 24.56±1.013 (A) | 25.97±0.394 (A) |

**Table. S19** continued on the next page

**Table. S19** continued from the previous page

| Terpenoid | Control | Mild drought | Moderate drought | Severe drought |
|---|---|---|---|---|
| C15H22-202(178/163) | 9.48±0.590 (A) | 8.63±0.698 (A) | 7.75±0.631 (A) | 7.77±0.283 (A) |
| C15H22-202(187/202)-1 | 26.21±0.705 (A) | 25.51±0.990 (A) | 24.56±1.013 (A) | 25.97±0.394 (A) |
| C15H26O-222(93/69/41) | 27.15±1.711 (A) | 26.14±1.508 (A) | 25.15±1.622 (A) | 28.82±0.966 (A) |
| C15H24 -204(123/81) | 4.74±0.501 (A) | 4.12±0.319 (A) | 3.99±0.346 (A) | 4.83±0.456 (A) |
| Humulene oxide II | 7.13±0.356 (A) | 7.48±0.655 (A) | 7.29±0.667 (A) | 7.55±0.752 (A) |
| C15H26O-222(59/161/91)-1 | 5.66±0.494 (A) | 4.74±0.185 (AB) | 4.50±0.213 (AB) | 3.60±0.331 (B) |
| γ-Eudesmol | 21.82±1.626 (A) | 22.42±1.046 (A) | 14.46±1.043 (B) | 14.49±2.934 (B) |
| C15H26O-222(105/59/161)-1 | 4.47±0.483 (A) | 4.15±0.349 (AB) | 3.30±0.290 (AB) | 2.95±0.167 (B) |
| C15H22-202(187/202)-2 | 9.01±1.130 (A) | 7.34±0.729 (A) | 6.84±0.594 (A) | 7.49±1.089 (A) |
| β-Eudesmol | 58.30±2.362 (A) | 55.18±4.806 (A) | 50.65±3.512 (A) | 47.29±6.024 (A) |
| α-Bisabolol | 18.39±0.613 (A) | 17.26±0.908 (A) | 17.15±0.703 (A) | 17.58±1.353 (A) |
| Juniper camphor | 14.14±0.491 (A) | 13.71±0.615 (A) | 13.41±0.389 (A) | 13.37±0.654 (A) |
| Total Mono, ppm | 6467.64±1372.625 (A) | 5129.16±622.669 (A) | 4638.12±657.516 (A) | 5271.58±877.546 (A) |
| Total Mono, % | 0.65±0.137 (A) | 0.51±0.062 (A) | 0.46±0.066 (A) | 0.53±0.088 (A) |
| Total Sesqui, ppm | 8063.90±977.881 (A) | 6944.61±578.395 (A) | 6666.30±501.992 (A) | 7935.24±595.644 (A) |
| Total Sesqui, % | 0.81±0.098 (A) | 0.69±0.058 (A) | 0.67±0.050 (A) | 0.79±0.060 (A) |
| Total Terpenoids (ppm) | 14531.54±2345.449 (A) | 12073.76±1160.635 (A) | 11304.42±1126.214 (A) | 13206.81±1453.464 (A) |
| Total Terpenoids (%) | 1.45±0.235 (A) | 1.21±0.116 (A) | 1.13±0.113 (A) | 1.32±0.145 (A) |

**Table S19.** Effects of the drought treatments on terpenoid concentrations in secondary inflorescences of 'Odem' plants (Experiment 2). Terpenoid concentrations (mg/L; ppm) in the secondary inflorescences of 'Odem' plants subjected to 4 different irrigation treatments, as measured at the end of Experiment 2. Samples were taken from the secondary inflorescences. Values are means ± SE. Different letters represent significant differences between irrigation treatments, according to one-way ANOVA and Tukey's HSD test ($P < 0.05$, $5 \leq N \leq 6$).

| Terpenes | Çontrol | Mild drought | Moderate drought | Severe drought |
|---|---|---|---|---|
| Santolina triene | 0.87±0.074 (A) | 0.78±0.125 (A) | 0.59±0.070 (A) | 0.51±0.065 (A) |
| C10H16-136(93/43/121/136) | 1.17±0.163 (A) | 1.25±0.236 (A) | 0.88±0.116 (A) | 0.82±0.178 (A) |
| 2-Thujene | 1.94±0.209 (A) | 1.61±0.170 (A) | 1.40±0.208 (A) | 1.46±0.165 (A) |
| α-Pinene | 210.61±19.625 (A) | 205.81±25.871 (A) | 163.65±11.429 (A) | 158.73±23.025 (A) |
| Camphene | 96.36±9.741 (A) | 94.63±11.773 (A) | 72.63±5.453 (A) | 71.66±10.305 (A) |
| Sabinene | 1.02±0.182 (A) | 0.91±0.194 (A) | 0.77±0.138 (A) | 0.84±0.126 (A) |
| β-Pinene | 295.40±24.503 (A) | 290.55±37.176 (A) | 232.48±23.905 (A) | 216.01±28.217 (A) |
| β-Myrcene | 506.40±47.215 (A) | 468.97±49.938 (A) | 437.09±42.170 (A) | 424.75±71.946 (A) |
| α-Phellandrene | 3.82±0.257 (A) | 3.64±0.337 (A) | 3.20±0.083 (A) | 3.32±0.274 (A) |
| α-Terpinene | 26.93±2.851 (A) | 25.54±3.042 (A) | 18.14±1.693 (A) | 19.64±1.703 (A) |
| Limonene | 1732.96±105.662 (A) | 1578.10±98.460 (A) | 1541.88±196.240 (A) | 1374.87±91.500 (A) |
| p-Cymene | 0.22±0.036 (A) | 0.17±0.034 (A) | 0.27±0.155 (A) | 0.14±0.037 (A) |
| β-Phellandrene | 8.41±0.801 (A) | 8.31±0.806 (A) | 7.69±0.331 (A) | 7.63±1.010 (A) |
| Eucalyptol | 9.79±1.355 (A) | 7.74±0.482 (A) | 6.22±0.910 (A) | 8.82±0.861 (A) |
| trans-Ocimene | 7.48±0.543 (A) | 7.90±0.741 (A) | 7.04±0.211 (A) | 7.57±0.835 (A) |
| γ-Terpinene | 20.72±1.633 (A) | 18.25±2.255 (A) | 15.19±1.566 (A) | 14.67±1.491 (A) |
| Terpinolene | 47.63±4.849 (A) | 45.73±6.998 (A) | 37.93±3.908 (A) | 36.48±3.248 (A) |
| Fenchone | 18.74±1.490 (A) | 17.85±2.340 (A) | 13.76±1.438 (A) | 12.54±1.353 (A) |
| Linalool | 317.99±15.550 (A) | 335.37±31.940 (A) | 298.96±17.062 (A) | 265.26±20.596 (A) |
| C10H18O-154(93/79/99/121)-1 | 31.34±2.884 (A) | 28.60±3.531 (AB) | 19.48±2.376 (BC) | 14.45±1.695 (C) |
| Fenchol | 313.21±29.298 (A) | 305.75±49.300 (A) | 227.01±27.499 (A) | 199.64±21.542 (A) |
| Terpinen-4-ol | 12.36±0.879 (A) | 11.93±1.572 (A) | 10.15±1.106 (A) | 9.35±0.935 (A) |
| Borneol | 125.14±1.625 (A) | 126.38±3.718 (A) | 120.00±2.101 (A) | 115.92±1.925 (A) |
| α-Terpineol | 218.78±12.201 (A) | 219.76±21.159 (A) | 190.62±10.295 (A) | 182.12±9.119 (A) |
| Nerol | 10.62±1.085 (A) | 10.41±1.671 (A) | 8.94±1.099 (A) | 7.53±0.912 (A) |
| Citronellol | 30.29±10.335 (A) | 30.70±8.885 (A) | 30.08±10.322 (A) | 26.53±8.213 (A) |
| Geraniol | 57.22±7.832 (AB) | 64.72±3.583 (A) | 65.73±0.888 (A) | 33.09±11.430 (AB) |
| Geranyl ethyl ether 1 | 71.52±1.570 (A) | 75.16±3.262 (A) | 71.15±2.630 (A) | 66.58±3.098 (A) |
| Bornyl acetate | 4.27±0.556 (A) | 4.27±0.524 (A) | 4.24±0.657 (A) | 13.03±5.853 (A) |
| Ylangene | 10.33±0.713 (A) | 7.51±1.301 (A) | 6.51±0.734 (A) | 7.48±1.109 (A) |
| α-Copaene | 13.08±0.879 (A) | 10.53±1.412 (A) | 9.40±0.785 (A) | 10.95±1.399 (A) |
| 7-epi-Sesquithujene | 10.83±0.655 (A) | 9.23±1.085 (A) | 8.65±0.704 (A) | 9.65±1.137 (A) |
| β-Isocomene | 22.16±1.046 (A) | 19.37±1.714 (A) | 17.80±1.052 (A) | 18.75±1.164 (A) |
| Sesquithujene | 48.04±4.435 (A) | 41.28±3.364 (A) | 43.04±2.073 (A) | 42.71±5.492 (A) |
| cis-α-Bergamotene | 45.08±4.349 (A) | 52.69±3.576 (A) | 54.07±3.447 (A) | 43.89±3.776 (A) |

**Table. S20** continued on the next page

**Table. S20** continued from the previous page

| Terpenes | Çontrol | Mild drought | Moderate drought | Severe drought |
|---|---|---|---|---|
| C15H24-204(105/91/147/189) | 18.99±1.218 (A) | 14.77±2.161 (A) | 12.80±1.127 (A) | 15.38±2.169 (A) |
| α-Santalene | 28.24±0.683 (A) | 26.55±1.197 (A) | 25.67±0.794 (A) | 26.35±1.829 (A) |
| β-Caryophyllene | 1379.17±82.122 (A) | 1151.96±138.31 (A) | 973.36±41.921 (A) | 1069.98±115.262 (A) |
| C15H24-204(120/91/79/69) | 8.65±0.403 (A) | 7.05±0.788 (AB) | 6.35±0.212 (B) | 6.52±0.578 (AB) |
| γ-Elemene | 152.41±4.883 (A) | 126.67±10.653 (A) | 113.32±10.778 (A) | 173.12±28.934 (A) |
| trans-α-Bergamotene | 258.33±7.205 (A) | 235.08±17.705 (A) | 221.47±10.450 (A) | 243.41±33.028 (A) |
| α-Guaiene | 434.91±12.673 (A) | 396.29±27.454 (A) | 366.30±16.116 (A) | 415.04±55.743 (A) |
| Guaia-6,9-diene | 6.22±0.286 (A) | 4.96±0.679 (A) | 4.52±0.595 (A) | 5.21±0.740 (A) |
| C15H24-204(69/91/105/161) | 3.24±0.333 (A) | 2.85±0.172 (A) | 2.74±0.209 (A) | 3.05±0.316 (A) |
| C15H24-204(105/91/133/161/189)-1 | 11.52±0.555 (A) | 9.10±1.106 (A) | 8.28±0.751 (A) | 9.46±1.275 (A) |
| α-Humulene | 705.97±37.418 (A) | 603.24±55.423 (A) | 547.97±8.876 (A) | 583.49±57.101 (A) |
| trans-β-Farnesene | 98.26±3.424 (A) | 91.38±4.703 (A) | 92.40±5.026 (A) | 92.15±7.606 (A) |
| Acoradiene | 7.09±0.216 (A) | 6.32±0.470 (A) | 5.78±0.384 (A) | 6.03±0.436 (A) |
| C15H24-204(105)-1 | 22.31±0.301 (A) | 21.12±0.755 (A) | 20.57±0.517 (A) | 20.89±1.281 (A) |
| C15H24-204(133/189)-1 | 46.92±1.004 (A) | 62.61±10.029 (A) | 99.32±18.626 (A) | 72.76±13.673 (A) |
| β-Chamigrene | 75.19±18.967 (A) | 67.06±14.099 (A) | 38.31±17.231 (A) | 61.03±17.012 (A) |
| α-Amorphene | 31.81±0.796 (A) | 28.35±1.342 (AB) | 26.56±0.683 (B) | 27.73±1.942 (AB) |
| C15H24-204(189/133)-2 | 79.63±1.174 (A) | 73.73±3.012 (A) | 71.56±3.575 (A) | 76.52±7.927 (A) |
| 4,5-di-epi-aristolochene | 20.11±0.229 (A) | 19.14±0.535 (A) | 18.58±0.392 (A) | 18.75±0.743 (A) |
| Sesquisabinene | 25.66±1.231 (A) | 25.53±0.987 (A) | 24.86±1.311 (A) | 21.93±2.000 (A) |
| α-Selinene | 85.16±2.663 (A) | 75.88±4.553 (A) | 69.90±2.703 (A) | 77.89±8.718 (A) |
| Valencene | 57.43±2.175 (A) | 48.81±3.587 (A) | 47.23±2.142 (A) | 45.62±4.126 (A) |
| β-Selinene | 196.06±5.334 (A) | 175.38±9.056 (A) | 167.63±6.509 (A) | 194.50±14.718 (A) |
| α-Zingiberene | 18.75±0.343 (A) | 20.45±1.432 (A) | 21.38±1.997 (A) | 22.60±2.665 (A) |
| δ-Guaiene | 557.52±6.967 (A) | 504.41±25.721 (A) | 503.51±25.011 (A) | 486.26±32.124 (A) |
| Dihydroagarofuran | 32.21±5.852 (A) | 30.68±4.932 (A) | 20.71±6.503 (A) | 28.89±6.717 (A) |
| β-Bisabolene | 24.56±4.989 (A) | 25.96±2.780 (A) | 29.28±5.048 (A) | 21.45±2.948 (A) |
| C15H24-204(similar to Germarcene B) | 39.20±1.204 (A) | 34.95±2.144 (A) | 32.57±1.499 (A) | 31.75±2.394 (A) |
| α-Farnesene | 102.47±5.047 (A) | 92.76±8.597 (A) | 88.36±9.719 (A) | 78.34±12.967 (A) |
| C15H24-204(161/105)-Eremophilene | 46.48±2.245 (A) | 39.43±2.366 (AB) | 36.37±1.790 (B) | 34.29±2.895 (B) |
| δ-Cadinene-C15H24-204(119/161/105/134) | 19.89±0.274 (A) | 18.76±0.627 (A) | 17.93±0.666 (A) | 17.75±0.781 (A) |
| Sesquicineole | 10.50±0.276 (A) | 9.74±0.485 (A) | 9.40±0.659 (A) | 9.42±1.073 (A) |
| C15H24-204(105)-2 | 51.57±1.239 (A) | 44.21±2.182 (AB) | 41.79±1.797 (B) | 38.81±3.105 (B) |
| β-Sesquiphellandrene | 26.36±0.467 (A) | 26.20±0.472 (A) | 26.10±0.892 (A) | 24.74±0.812 (A) |
| C15H24-204(189/133)-3 | 291.35±9.169 (A) | 241.31±13.146 (AB) | 227.66±13.851 (B) | 216.41±19.600 (B) |

**Table. S20** continued on the next page

**Table. S20** continued from the previous page

| Terpenes | Çontrol | Mild drought | Moderate drought | Severe drought |
|---|---|---|---|---|
| C15H24-204(161/133/105) | 3261.92±129.24 (A) | 2658.06±156.2 (AB) | 2537.80±173.405(B) | 2326.01±238.541 (B) |
| Selina-3,7(11)-diene | 5749.80±218.31 (A) | 4925.88±303 (AB) | 4661.82±334.3 (AB) | 4197.11±434.368 (B) |
| trans-α-Bisabolene | 38.68±1.870 (A) | 37.87±1.127 (A) | 38.79±2.166 (A) | 35.77±3.427 (A) |
| C15H24-202(202/131/145/159) | 4.96±0.383 (A) | 4.41±0.491 (A) | 4.23±0.652 (A) | 3.21±0.529 (A) |
| Cadala-1(10),3,8-triene-204(157/142) | 51.46±1.524 (A) | 46.71±1.838 (A) | 47.40±3.388 (A) | 43.96±2.444 (A) |
| Germacrene B | 147.68±8.408 (A) | 118.91±12.015 (A) | 119.38±14.090 (A) | 163.01±24.793 (A) |
| trans-Nerolidol | 434.05±25.725 (A) | 457.11±24.767 (A) | 489.06±44.853 (A) | 470.62±34.835 (A) |
| Caryophyllene oxide | 37.69±1.400 (A) | 38.92±2.628 (A) | 37.01±3.585 (A) | 34.60±2.262 (A) |
| C15H22-202(178/163) | 14.57±0.931 (A) | 14.17±1.103 (A) | 12.97±1.085 (A) | 10.66±1.408 (A) |
| C15H22-202(187/202)-1 | 37.69±1.400 (A) | 38.92±2.628 (A) | 37.01±3.585 (A) | 34.60±2.262 (A) |
| Guaiol | 446.62±12.271 (A) | 445.73±18.744 (A) | 431.11±32.900 (A) | 401.63±16.128 (A) |
| α-epi-7-epi-5-Eudesmol | 14.67±0.574 (A) | 14.50±0.689 (A) | 13.91±0.968 (A) | 13.19±0.590 (A) |
| Humulene oxide II | 25.01±0.416 (A) | 25.58±1.082 (A) | 25.56±1.301 (A) | 23.51±1.191 (A) |
| C15H26O-222(similar γ-Eudesmol) | 2396.81±83.881 (A) | 2394.01±93.909 (A) | 2376.11±122.27 (A) | 2130.26±58.558 (A) |
| C15H26O-222(59/161/91)-1 | 37.81±3.038 (A) | 39.79±2.620 (A) | 40.06±1.414 (A) | 36.80±1.064 (A) |
| γ-Eudesmol | 172.08±3.972 (A) | 173.03±8.315 (A) | 171.02±11.405 (A) | 165.17±9.394 (A) |
| C15H26O-222(105/59/161)-1 | 37.89±0.662 (A) | 38.30±0.943 (A) | 38.17±1.000 (A) | 39.00±2.749 (A) |
| Agaruspirol | 40.78±1.483 (A) | 38.35±1.613 (A) | 34.90±1.882 (A) | 37.43±1.787 (A) |
| Cubenol | 32.03±0.839 (A) | 30.93±1.518 (A) | 30.57±1.180 (A) | 28.33±1.523 (A) |
| C15H24O-220(136/91/69) | 10.89±0.104 (A) | 11.43±0.276 (A) | 11.07±0.476 (A) | 10.15±0.379 (A) |
| C15H26O-222(59) | 12.61±0.438 (A) | 12.15±0.655 (A) | 12.04±0.824 (A) | 11.10±0.589 (A) |
| C15H22-202(187) | 4.89±0.561 (A) | 5.71±0.763 (A) | 5.23±1.017 (A) | 4.56±0.554 (A) |
| epi-γ-Eudesmol-C15H26O-222(105/59/161)-2 | 265.80±10.694 (A) | 252.97±15.040 (A) | 248.21±12.514 (A) | 229.17±8.924 (A) |
| α-Eudesmol-C15H26O-222(59/161/91)-2 | 17.92±2.372 (A) | 20.35±3.128 (A) | 17.91±3.356 (A) | 17.54±1.721 (A) |
| C15H22-202(187/202)-2 | 24.52±0.334 (AB) | 24.82±0.522 (A) | 24.32±0.572 (AB) | 22.56±0.436 (B) |
| β-Eudesmol | 310.75±125.485 (A) | 316.77±100.572 (A) | 503.49±28.880 (A) | 359.28±89.965 (A) |
| C15H24O-(similar Caryophyllene oxide)-1 | 25.10±0.413 (A) | 23.62±0.648 (A) | 23.79±0.906 (A) | 22.05±0.454 (A) |
| 7-epi-α-Eudesmol | 97.43±2.926 (A) | 97.34±5.007 (A) | 88.56±4.378 (A) | 93.69±3.174 (A) |
| Bulnesol | 130.27±6.117 (A) | 124.98±5.610 (A) | 119.50±6.823 (A) | 114.45±3.493 (A) |
| α-Bisabolol | 111.84±6.182 (A) | 107.62±5.089 (A) | 107.99±7.037 (A) | 110.71±8.637 (A) |
| Juniper camphor | 27.55±1.153 (A) | 23.68±1.591 (A) | 23.93±0.835 (A) | 23.73±0.793 (A) |
| Total MONO, ppm | 4191.18±266.91 (A) | 3998.81±330.92 (A) | 3615.19±288.26 (A) | 3301.98±268.861 (A) |
| Total MONO, % | 0.42±0.027 (A) | 0.40±0.033 (A) | 0.36±0.029 (A) | 0.33±0.027 (A) |
| Total Sesqui, ppm | 19209.73±479.1 (A) | 17125.85±847.3 (A) | 16532.52±838.6 (A) | 15676.70±1104.6 (A) |
| Total Sesqui, % | 1.92±0.048 (A) | 1.71±0.085 (A) | 1.65±0.084 (A) | 1.57±0.110 (A) |
| Total all, % | 2.34±0.034 (A) | 2.11±0.115 (AB) | 2.01±0.103 (AB) | 1.90±0.127 (B) |

**Table S20.** Effects of the drought treatments on terpenoid concentrations in primary inflorescences of 'MVA' plants (Experiment 3). Terpenoid concentrations (mg/L; ppm) in the primary inflorescences of 'MVA' plants subjected to 4 different irrigation treatments, as measured at the end of Experiment 3. Samples were taken from the primary inflorescences. Values are means ± SE. Different letters represent significant differences between irrigation treatments, according to one-way ANOVA and Tukey's HSD test ($P < 0.05$, $5 \leq N \leq 6$).

| Terpenoid | Control | Mild drought | Moderate drought | Severe drought |
|---|---|---|---|---|
| Santolina triene | 0.986±0.13027 (A) | 0.792±0.10367 (AB) | 0.601±0.03229 (B) | 0.517±0.05483 (B) |
| C10H16-136(93/43/121/136) | 1.359±0.20947 (A) | 1.252±0.26310 (A) | 0.756±0.15750 (A) | 1.062±0.24241 (A) |
| 2-Thujene | 2.251±0.28912 (A) | 1.766±0.12272 (AB) | 1.337±0.05201 (B) | 1.912±0.35185 (AB) |
| α-Pinene | 250.801±35.828 (A) | 216.633±21.070 (AB) | 167.308±11.03 (AB) | 149.630±11.16978 (B) |
| Camphene | 116.238±17.449 (A) | 100.124±10.446 (AB) | 74.267±6.775 (AB) | 70.374±6.21867 (B) |
| Sabinene | 1.191±0.16905 (A) | 1.056±0.16338 (A) | 0.818±0.10626 (A) | 0.938±0.14298 (A) |
| β-Pinene | 353.274±50.580 (A) | 304.187±30.951 (AB) | 244.300±17.63 (AB) | 214.950±17.84754 (B) |
| β-Myrcene | 526.986±45.553 (A) | 472.159±45.48182 (A) | 416.046±41.409 (A) | 442.161±55.85338 (A) |
| α-Phellandrene | 4.055±0.30524 (A) | 3.791±0.16761 (A) | 3.386±0.12721 (A) | 3.415±0.14953 (A) |
| α-Terpinene | 31.896±3.23413 (A) | 29.819±1.56104 (A) | 24.824±2.94525 (A) | 23.090±2.21844 (A) |
| Limonene | 1914.91±173.23 (A) | 1798.460±103.266 (A) | 1489.924±155.33 (A) | 1370.761±69.909 (A) |
| p-Cymene | 0.251±0.03668 (A) | 0.234±0.02696 (A) | 0.249±0.10276 (A) | 0.177±0.01694 (A) |
| β-Phellandrene | 9.292±0.94047 (A) | 9.065±0.57950 (A) | 8.022±0.54914 (A) | 7.611±0.50432 (A) |
| Eucalyptol | 10.979±1.15838 (A) | 7.975±0.38238 (A) | 7.243±0.62392 (A) | 13.681±4.31277 (A) |
| trans-Ocimene | 8.172±0.49953 (A) | 8.406±0.46897 (A) | 6.843±0.50125 (A) | 7.301±0.39812 (A) |
| γ-Terpinene | 23.046±2.45832 (A) | 19.119±1.44267 (A) | 16.604±1.56707 (A) | 17.133±1.93245 (A) |
| Terpinolene | 53.763±7.02605 (A) | 48.628±4.52508 (A) | 35.339±3.38775 (A) | 35.993±3.16309 (A) |
| Fenchone | 20.487±2.05693 (A) | 17.975±1.95314 (AB) | 12.697±1.31143 (B) | 12.743±0.94474 (B) |
| Linalool | 328.856±18.966 (A) | 338.909±20.88988 (A) | 278.767±20.3684 (A) | 272.089±9.51804 (A) |
| C10H18O-154(93/79/99/121)-1 | 34.933±5.02009 (A) | 27.278±3.82935 (AB) | 17.865±2.02691 (B) | 14.526±1.38300 (B) |
| Fenchol | 352.914±52.987 (A) | 305.921±39.97974 (A) | 220.394±21.7401 (A) | 210.011±19.29340 (A) |
| Terpinen-4-ol | 13.708±1.72227 (A) | 12.584±0.69152 (A) | 9.848±0.91723 (A) | 9.742±0.86323 (A) |
| Borneol | 122.694±3.983 (A) | 122.451±2.51938 (A) | 112.328±7.47698 (A) | 115.988±3.20630 (A) |
| α-Terpineol | 230.694±20.073 (A) | 218.600±14.64450 (A) | 181.083±8.63298 (A) | 180.562±7.88364 (A) |
| Nerol | 11.852±1.94085 (A) | 10.969±1.29822 (A) | 8.016±0.96940 (A) | 7.315±0.89393 (A) |
| Citronellol | 35.765±12.358 (A) | 37.476±9.97189 (A) | 26.126±8.79858 (A) | 26.302±9.61361 (A) |
| Geraniol | 53.986±9.008 (AB) | 65.505±1.68974 (A) | 64.284±1.89299 (A) | 31.276±10.32147 (B) |
| Geranyl ethyl ether 1 | 71.312±2.99312 (A) | 75.145±3.32756 (A) | 68.504±4.01659 (A) | 69.107±4.22892 (A) |
| Bornyl acetate | 4.821±0.52322 (AB) | 5.151±0.57149 (AB) | 4.133±0.46830 (B) | 7.139±0.80493 (A) |
| Ylangene | 9.991±1.12900 (A) | 7.625±0.56645 (AB) | 5.304±0.37338 (B) | 7.220±0.63912 (AB) |
| α-Copaene | 12.983±1.30257 (A) | 10.740±0.62110 (AB) | 8.106±0.58661 (B) | 10.678±0.81236 (AB) |
| 7-epi-Sesquithujene | 10.773±1.26270 (A) | 9.516±0.39448 (AB) | 7.367±0.35835 (B) | 9.042±0.39308 (AB) |
| β-Isocomene | 21.296±1.46803 (A) | 19.108±0.74017 (AB) | 15.879±1.26996 (B) | 18.686±0.34000 (AB) |
| Sesquithujene | 43.375±5.019 (AB) | 53.142±2.86911 (A) | 39.777±2.22693 (B) | 51.799±2.46270 (AB) |
| cis-α-Bergamotene | 45.776±3.25666 (A) | 44.899±3.09942 (A) | 47.567±1.83743 (A) | 41.169±3.17365 (A) |

**Table. S21** continued on the next page

**Table. S21** continued from the previous page

| Terpenoid | Control | Mild drought | Moderate drought | Severe drought |
|---|---|---|---|---|
| C15H24-204(105/91/147/189) | 18.596±2.06376 (A) | 14.936±1.08625 (A) | 11.219±0.80480 (A) | 15.788±1.23987 (A) |
| α-Santalene | 27.784±1.49312 (A) | 26.360±0.40951 (AB) | 23.797±0.47322 (B) | 26.385±0.67094 (AB) |
| β-Caryophyllene | 1193.164±96.729 (A) | 1115.968±75.5334(AB) | 897.334±28.2292(B) | 1020.413±64.56 (AB) |
| C15H24-204(120/91/79/69) | 7.831±0.45922 (A) | 6.758±0.27458 (AB) | 5.701±0.25406 (B) | 6.857±0.28269 (AB) |
| γ-Elemene | 140.969±11.253 (AB) | 121.870±8.830 (AB) | 113.607±8.95805 (B) | 167.350±19.63103 (A) |
| trans-α-Bergamotene | 260.878±21.326 (A) | 237.716±10.92965 (A) | 203.754±7.81101 (A) | 240.765±22.93132 (A) |
| α-Guaiene | 417.704±20.398 (A) | 396.800±22.70535 (A) | 354.367±26.046 (A) | 445.971±49.79918 (A) |
| Guaia-6,9-diene | 5.822±0.20587 (A) | 5.188±0.35494 (A) | 3.940±0.08060 (B) | 5.011±0.33436 (AB) |
| C15H24-204(105/91/133/161/189)-1 | 10.712±0.87341 (A) | 9.201±0.39240 (A) | 7.018±0.44932 (B) | 9.414±0.55319 (A) |
| α-Humulene | 667.955±48.807 (A) | 598.542±26.628 (AB) | 497.411±22.035 (B) | 570.330±43.893 (AB) |
| trans-β-Farnesene | 98.184±5.00153 (A) | 94.296±5.62809 (A) | 84.677±3.16152 (A) | 90.901±4.35052 (A) |
| Acoradiene | 6.532±0.41392 (A) | 6.316±0.22196 (A) | 5.216±0.13244 (B) | 6.291±0.34042 (AB) |
| C15H24-204(105)-1 | 21.050±0.43149 (A) | 21.002±0.52319 (A) | 19.717±0.26425 (A) | 21.318±0.59764 (A) |
| C15H24-204(133/189)-1 | 58.866±7.60923 (A) | 61.440±11.15472 (A) | 97.733±12.12290 (A) | 79.394±17.88339 (A) |
| α-Amorphene | 29.458±0.96028 (A) | 27.742±0.74003 (A) | 24.922±0.58206 (B) | 28.513±1.13778 (A) |
| C15H24-204(189/133)-2 | 75.266±2.31240 (A) | 75.071±3.90000 (A) | 61.710±3.91028 (A) | 75.629±4.34301 (A) |
| 4,5-di-epi-aristolochene | 18.813±0.36774 (A) | 18.752±0.30744 (A) | 19.246±0.95156 (A) | 18.718±0.46100 (A) |
| Sesquisabinene | 23.300±1.36349 (A) | 24.628±0.77996 (A) | 23.468±0.98956 (A) | 23.095±0.88297 (A) |
| α-Selinene | 79.959±3.11683 (A) | 75.911±2.30407 (AB) | 64.536±2.61445 (B) | 79.755±6.03550 (A) |
| Valencene | 55.789±3.67848 (A) | 49.678±1.97197 (AB) | 42.640±2.98423 (B) | 45.706±2.64060 (AB) |
| β-Selinene | 184.565±5.97989 (A) | 174.606±9.75622 (AB) | 149.495±6.30481 (B) | 183.303±10.59810 (A) |
| α-Zingiberene | 17.742±0.53313 (A) | 17.622±0.59160 (A) | 19.950±1.77155 (A) | 18.076±1.01733 (A) |
| δ-Guaiene | 526.264±7.75415 (A) | 498.482±20.18073 (A) | 460.068±18.1764 (A) | 508.116±36.84225 (A) |
| Dihydroagarofuran | 30.261±5.80088 (A) | 29.576±4.59693 (A) | 19.420±6.21502 (A) | 28.559±5.93871 (A) |
| β-Bisabolene | 20.664±3.64486 (A) | 28.982±3.53625 (A) | 29.683±3.53719 (A) | 23.028±3.77600 (A) |
| C15H24-204(similar to Germarcene B) | 37.101±1.90763 (A) | 34.108±1.53796 (AB) | 30.337±0.74565 (B) | 32.483±1.20956 (AB) |
| α-Farnesene | 92.520±8.24256 (A) | 94.653±9.44741 (A) | 81.640±9.49222 (A) | 77.584±10.21106 (A) |
| C15H24-204(161/105)-Eremophilene | 42.381±1.81103 (A) | 38.584±1.53074 (AB) | 33.571±1.40254 (B) | 35.269±1.45853 (B) |
| δ-Cadinene-C15H24-204(119/161/105/134) | 18.458±0.56416 (A) | 18.127±0.63631 (A) | 17.165±0.39105 (A) | 17.768±0.39322 (A) |
| Sesquicineole | 9.588±0.60538 (A) | 9.408±0.40111 (A) | 8.309±0.28809 (A) | 9.205±0.60476 (A) |
| C15H24-204(105)-2 | 47.794±1.36086 (A) | 43.852±1.26139 (AB) | 38.636±0.99490 (C) | 39.214±1.41573 (BC) |
| β-Sesquiphellandrene | 25.352±0.70565 (A) | 25.419±0.80600 (A) | 25.285±0.31211 (A) | 24.691±0.78074 (A) |
| C15H24-204(189/133)-3 | 267.203±10.079 (A) | 235.082±12.492 (AB) | 198.554±11.909 (B) | 220.423±11.294 (AB) |
| C15H24-204(161/133/105) | 2967.948±84.870 (A) | 2601.481±120.67 (AB) | 2463.889±144.71 (B) | 2329.158±105.21 (B) |
| Selina-3,7(11)-diene | 5550.251±127.66 (A) | 4786.593±234.08 (AB) | 4287.498±249.08 (B) | 4216.729±355.99 (B) |

**Table. S21** continued on the next page

**Table. S21** continued from the previous page

| Terpenoid | Control | Mild drought | Moderate drought | Severe drought |
|---|---|---|---|---|
| trans-α-Bisabolene | 36.608±0.62389 (A) | 36.125±1.85101 (A) | 35.774±1.64892 (A) | 35.953±1.68850 (A) |
| C15H24-202(202/131/145/159) | 4.445±0.44331 (A) | 3.976±0.37421 (A) | 3.527±0.55151 (A) | 3.592±0.40758 (A) |
| Cadala-1(10),3,8-triene-204(157/142) | 44.621±2.12763 (A) | 42.891±2.49773 (A) | 43.881±1.83642 (A) | 44.535±1.32009 (A) |
| Germacrene B | 133.052±9.177 (AB) | 121.820±9.766 (AB) | 114.761±10.533 (B) | 159.937±13.563 (A) |
| trans-Nerolidol | 427.309±19.444 (A) | 406.638±32.349 (A) | 517.015±41.514 (A) | 485.245±28.16 (A) |
| Caryophyllene oxide | 36.182±1.69062 (A) | 35.379±2.53723 (A) | 34.122±2.94357 (A) | 34.872±1.80904 (A) |
| C15H22-202(178/163) | 14.106±1.02049 (A) | 12.286±1.21029 (A) | 10.947±1.27534 (A) | 10.933±0.76935 (A) |
| C15H22-202(187/202)-1 | 36.182±1.69062 (A) | 35.379±2.53723 (A) | 34.122±2.94357 (A) | 34.872±1.80904 (A) |
| Guaiol | 421.734±10.568 (A) | 392.834±29.799 (A) | 392.229±30.063 (A) | 405.521±16.316 (A) |
| α-epi-7-epi-5-Eudesmol | 13.973±0.35925 (A) | 12.645±1.04154 (A) | 13.209±0.53043 (A) | 13.654±0.93857 (A) |
| Humulene oxide II | 23.495±0.65908 (A) | 23.955±1.16707 (A) | 22.699±2.07781 (A) | 23.328±0.61348 (A) |
| C15H26O-222(similar γ-Eudesmol) | 2237.077±55.832 (A) | 2142.536±147.725 (A) | 2241.165±117.36 (A) | 2186.382±125.31 (A) |
| C15H26O-222(59/161/91)-1 | 34.771±1.68149 (A) | 35.979±2.23369 (A) | 38.063±1.79084 (A) | 38.968±2.87298 (A) |
| γ-Eudesmol | 165.948±3.92292 (A) | 154.260±7.60005 (A) | 164.821±3.89218 (A) | 164.276±14.61531 (A) |
| C15H26O-222(105/59/161)-1 | 36.031±0.78117 (A) | 38.509±1.23322 (A) | 39.375±1.91336 (A) | 36.813±1.72191 (A) |
| Agaruspirol | 36.469±0.90761 (A) | 34.366±2.24848 (A) | 33.242±1.18097 (A) | 36.596±2.92608 (A) |
| Cubenol | 28.905±0.23191 (A) | 28.352±1.18553 (A) | 27.436±1.06838 (A) | 29.698±1.83224 (A) |
| C15H24O-220(136/91/69) | 10.379±0.21518 (A) | 10.546±0.42383 (A) | 10.552±0.26601 (A) | 10.471±0.51254 (A) |
| C15H26O-222(59) | 12.200±0.43109 (A) | 10.513±0.74709 (A) | 10.917±0.75294 (A) | 11.884±0.69049 (A) |
| C15H22-202(187) | 4.899±0.74040 (A) | 4.772±0.58913 (A) | 4.385±0.66037 (A) | 4.873±0.33044 (A) |
| epi-γ-Eudesmol-C15H26O-222(105/59/161)-2 | 246.749±7.24874 (A) | 218.883±13.20961 (A) | 230.546±12.4636 (A) | 244.810±18.24975 (A) |
| α-Eudesmol-C15H26O-222(59/161/91)-2 | 17.422±2.90090 (A) | 16.702±2.09133 (A) | 15.186±2.82825 (A) | 17.409±1.30600 (A) |
| C15H22-202(187/202)-2 | 23.535±0.95419 (A) | 21.852±1.31890 (A) | 23.968±0.53738 (A) | 23.291±1.15645 (A) |
| β-Eudesmol | 282.600±114.275 (A) | 260.463±81.568 (A) | 455.287±33.310 (A) | 367.182±94.41548 (A) |
| C15H24O-(similar Caryophyllene oxide)-1 | 22.961±0.83450 (A) | 21.862±1.22513 (A) | 22.261±0.97192 (A) | 22.851±1.35728 (A) |
| 7-epi-α-Eudesmol | 91.148±2.29740 (A) | 85.180±5.95242 (A) | 78.212±5.14805 (A) | 96.061±6.56756 (A) |
| Bulnesol | 121.787±3.44463 (A) | 111.167±4.96389 (A) | 109.246±6.20891 (A) | 117.448±7.90416 (A) |
| α-Bisabolol | 104.512±6.22075 (A) | 94.349±4.59436 (A) | 97.227±9.60254 (A) | 111.654±12.05569 (A) |
| Juniper camphor | 26.017±0.78914 (A) | 22.743±0.96287 (AB) | 21.292±0.60117 (B) | 24.293±1.77647 (AB) |
| Total MONO, ppm | 4599.500±435.02 (A) | 4269.43±245.59 (AB) | 3509.91±226.02 (AB) | 3325.5±137.545 (B) |
| Total MONO, % | 0.460±0.04350 (A) | 0.427±0.02456 (AB) | 0.351±0.02260 (AB) | 0.333±0.01375 (B) |
| Total Sesqui, ppm | 18034.61±263.33 (A) | 16332.37±578.9 (AB) | 15466.39±604.63 (B) | 15791.12±647.4 (AB) |
| Total Sesqui, % | 1.803±0.02633 (A) | 1.633±0.05789 (AB) | 1.547±0.06046 (B) | 1.579±0.06475 (AB) |
| Total Terpenoids, % | 2.263±0.06085 (A) | 2.060±0.06487 (AB) | 1.898±0.07928 (B) | 1.912±0.06512 (B) |

**Table S21.** Effects of the drought treatments on terpenoid concentrations in secondary inflorescences of 'MVA' plants (Experiment 3). Terpenoid concentrations (mg/L; ppm) in the secondary inflorescences of 'MVA' plants subjected to 4 different irrigation treatments, as measured at the end of Experiment 3. Samples were taken from the secondary inflorescences. Values are means ± SE. Different letters represent significant differences between irrigation treatments, according to one-way ANOVA and Tukey's HSD test ($P < 0.05$, $5 \leq N \leq 6$).
